# Location dependence of protein intrinsic disorder in *Drosophila melanogaster*

**DOI:** 10.64898/2026.07.02.732782

**Authors:** Hassan Sibroe Abdulla Daanaa, Shigehiro Kuraku, Hiroshi Akashi, Kuniaki Saito

## Abstract

The relevance of protein structural flexibility in function remains contested, but experimental and computational evidence continues to accumulate. Many efforts to address this investigate intrinsic disorder, which commonly refers to peptide segments or entire protein sequences that presumably lack structure and exhibit high flexibility/conformational heterogeneity under physiological conditions. These efforts face challenges such as conflicting computational predictions and ambiguous relationships among intrinsic disorder locations and other protein properties. We address these challenges at a genome-wide scale in *Drosophila melanogaster* using residue-level predictions for various protein properties. We employ single and consensus approaches to quantify the prevalence of intrinsic disorder and attempt to infer function by testing for differences along protein sequences. Intrinsic disorder is likely more common at terminals than internal regions, and amino acid frequencies can vary substantially between regions in a manner that plausibly reflects functions of intrinsic disorder, rather than only proteome-wide effects. Tertiary structure potentially underlies the prevalence of intrinsic disorder along protein sequences; this prevalence varies more in a putatively solvent-exposed context than a solvent-buried one. Protein-binding appears to be a main function of intrinsic disorder, and we find support consistent with the notion that structural flexibility fosters binding plasticity, and show that location and protein length are factors in this relationship. Nucleic acid-binding and linker are ostensibly less common disorder functions than protein-binding, but nucleic acid-binding seems more localized at terminals. Residue-level estimates of selection pressure indicate that disordered regions generally evolve under weaker sequence constraints than structured regions, except at the N-terminal region. Biases in disorder prediction are a considerable factor in many of the observations, but unlikely a full explanation. The findings strengthen support for functional relevance of flexibility, offer insight into protein architecture and function, and lend impetus for experimental inquiry.

## Introduction

Understanding the relevance of protein structural flexibility in function and evolution is an ongoing challenge. Addressing this challenge often involves tests of the disorder-function paradigm, which posits that protein function can arise without a well-defined structure. This paradigm has not yet been firmly established despite substantial evidence that continues to accumulate [reviewed in 1–3]. Tests of the paradigm focus on intrinsic disorder, which are peptide segments or whole proteins that are presumably unstructured under physiological conditions and exhibit high flexibility/conformational heterogeneity. Experimental studies identifying intrinsic disorder – using methods such as X-ray crystallography and nuclear magnetic resonance spectroscopy – have revealed compositional biases that favor charged and/or hydrophilic residues, and high solvent accessibility (i.e., increased surface exposure). Such studies support roles of intrinsic disorder in functional molecular interactions [e.g., 4–6], and in allowing proteins to adopt multiple functional conformations in 3D space [e.g., 7–9]. Despite these insights, experimental annotations of intrinsic disorder are scant, motivating computational approaches for prediction.

Early computational approaches for intrinsic disorder prediction were grounded in experimental observations and/or theoretical principles [reviewed in 10]. Since the primary structure of a protein largely determines its biophysical (and functional) properties, many computational approaches [10–12] relied on amino acid composition and physicochemical properties for intrinsic disorder prediction, with net charge and/or hydrophobicity being among main properties of focus [13,14]. Later computational approaches – leveraging advances in machine learning algorithms and/or protein structure prediction – improved prediction accuracy by incorporating various information [reviewed in 10,15]. Such information includes residue-residue contacts [16,17], 3D structure information [18], and homology profiles [e.g., 19–22]. Many of these approaches allow prediction of various classes of intrinsic disorder such as short and long disordered regions or disordered binding sites. The programs implementing these approaches typically require an input protein sequence or structure and return residue-level “disorder scores” specifying propensities. Analysis of the resulting predictions have generally corroborated findings from experimental data, and critically, have fostered large-scale studies of many organisms including fruit fly and human. These studies present evidence that terminal residues are ostensibly frequently disordered [23] and may be enriched in both disorder and protein-binding functions [24]. In addition, compared to structured regions, disordered regions are associated with faster evolution [25] and more protein adaptive evolution [26–28], but have also been associated with purifying selection [25,29].

Despite insights from experimental and computational studies, relationships between intrinsic disorder and protein architecture, which can inform on functional relevance, have received less attention. Disordered regions can occur at any location (terminal or internal) within a protein sequence, and previous studies – confined to a subset of residue positions – support prevalence/enrichment of intrinsic disorder near terminal residues in multiple taxa such as fruit fly, human and *Escherichia coli*. However, the challenge remains to account for the full length of protein sequences and to investigate candidate causal factors in the prevalence of intrinsic disorder along protein sequences, and address the possible main functions of intrinsic disorder. Hurdles to this challenge include conflicting predictions among disorder prediction programs, owing to differing optimizations, and include controlling for confounding factors in evolutionary analysis (e.g., incorporation of homology profiles for disorder prediction). Addressing these hurdles, at a genome-wide scale may benefit from considering protein length since the upper limit of a disordered region/protein will depend on protein length.

We address the challenges above primarily in *Drosophila melanogaster*. Publicly available databases provide high quality genome assemblies and gene annotation data for *D. melanogaster* and its close relatives. We develop a consensus approach to predict intrinsic disorder using flDPnn, AlphaFold-based approaches and IUPred3. The resulting predictions are used to quantify the prevalence of disordered residues and test associations with several factors (e.g., location within the protein and amino acid composition), while controlling for protein length. Differences in the prevalence of intrinsic disorder along protein sequences may hint at biological significance, and we present new support consistent with the disorder-function paradigm. Intrinsic disorder is likely more prevalent at protein terminals than at internal regions, an observation supported in multiple other taxa including human and *E. coli*. This apparent location dependence is associated with regional variation in the frequency of amino acids, and some of this variation can plausibly be attributed to intrinsic disorder presence rather than a proteome-wide effect. Interestingly, tertiary structure context (i.e., solvent-exposed and -buried states) is a major factor in the extent of location dependence. Analysis of predicted disorder functions reveals that protein-binding may be the most prevalent function of intrinsic disorder relative to domain-linker, RNA-binding, and DNA-binding functions, but the latter two appear more strongly associated with the location dependence. Interestingly, hubs, proteins with many protein interactors, generally have more intrinsic disorder than non-hubs, consistent with previous studies and with the notion that structural flexibility allows functional interactions with multiple structurally diverse partners. However, we find that protein length and disorder locations can have a considerable effect on this relationship. We obtain estimates of protein evolutionary rates and selection pressure at disordered and non-disordered residues, and find support for faster evolution and relaxed purifying selection at disordered residues, along protein segments but with a notable exception to the N-terminal region, where these estimates are generally similar between the residue classes.

## Results

### Uneven distribution of disordered residues along proteins

Differences in the prevalence of intrinsic disorder along protein sequences can hint at function. We test for such differences by comparing N-terminal, internal and C-terminal segments for the fraction of predicted disordered residues, which is a commonly-used statistic to quantify prevalence and is the ratio between counts of disordered residues and all residues. We calculate this fraction separately for bins representing short, medium and long protein sequences (explained in *Material and Methods*), and find substantially higher fractions at terminal segments than internal ones across all protein length bins (Fig 1, S3, S4 Figs and S7 Table). We observe the greatest extent of difference in the medium length bin followed by the long and short ones (S7 Table). These results support genome-wide prevalence of disordered regions at protein terminals relative to internal regions, and hint that intrinsic disorder may have a general functional role in many proteins, but also possibly, specified roles depending on location within the protein.

**Fig 1.**
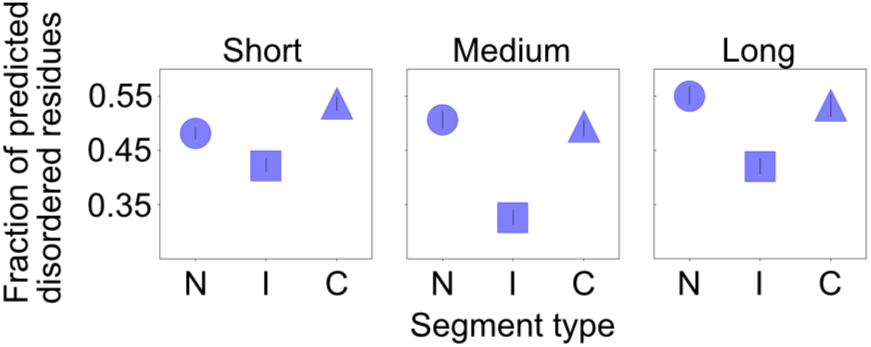
Prevalence of intrinsic disorder along protein sequences. The fraction of predicted disordered residues is the ratio between total counts of predicted disordered residues and all residues. This fraction was calculated separately for N-terminal (N), Internal (I), and C-terminal (C) segments of protein sequences with 80 residues or above. Error bars indicate 95% confidence intervals calculated among 500 replicates created by sampling genes with replacement. n=3361, 2315 and 2146 genes for the short, medium and long protein length bins, respectively. Data binning is explained in *Materials and Methods* and statistical test results are presented in S7 Table.

### Amino acid compositional heterogeneity among disordered residues

Compositional trends of disordered regions relative to non-disordered regions or among classes of disordered region lengths have been studied extensively. We test whether such trends reflect proteome-wide patterns or functional differences attributable to presence of intrinsic disorder. Differences between protein segments for amino acid frequencies among disordered residues may suggest a biological signal.

We calculate a statistic, the scaled amino acid frequency difference (sAA_freq_diff), to analyze amino acid frequencies among disordered residues considering the frequencies for non-disordered residues (*Materials and Methods*). sAA_freq_diff of several amino acids such as Met, Lys and Thr vary substantially among segments, but sAA_freq_diff for most other amino acids, including Cys, Glu, Ile, Leu, Phe, Pro and Val varies between two segments or more and such variation can be particular to protein length bins. Patterns for Met are notable; sAA_freq_diff for N-terminal is substantially higher than those of internal and C-terminal segments across all protein length bins (Fig 2 and S8 Table) and amino acid frequency trends clearly differ between predicted disordered and non-disordered residues (S5–7 Figs, S8 and S9 Tables). Initiation Met underlies these patterns; the frequency of Met at the N-terminal segment plummets drastically, when initiation Met is filtered (S8 Fig). These compositional analyses suggest that proteome-wide effects dominate amino acid frequency distributions at disordered and non-disordered regions, but support exists for effects attributable to intrinsic disorder presence, consistent functional differences with respect to binding targets and/or extent of flexibility.

**Fig 2.**
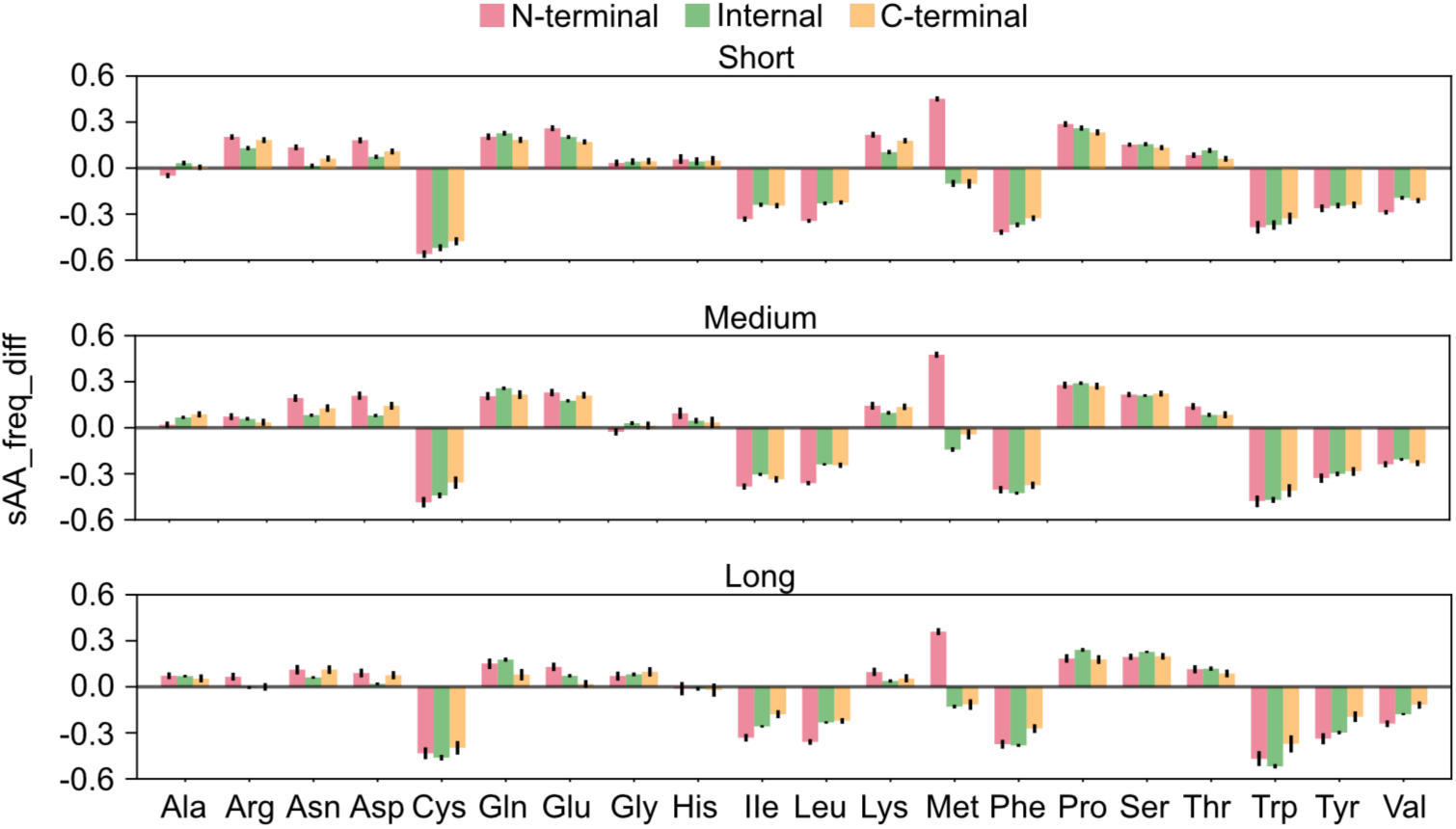
Scaled amino acid frequency differences (sAA_freq_diff) of disordered residues. sAA_freq_diff is used to compare protein segments for amino acid frequencies at predicted disordered residues, accounting for the frequencies at predicted non-disordered residues (see *Materials and Methods*). sAA_freq_diff>0 and sAA_freq_diff<0 indicate higher and lower frequencies, respectively, among the disordered residues. Error bars indicate 95% confidence intervals calculated among 1000 replicates that were created by sampling genes with replacement. n=3361, 2315 and 2146 genes for the short, medium and long protein length bins, respectively. Statistical tests results are shown in S8 and S9 Tables.

### Solvent accessibility associations with the distribution of disordered residues

High solvent accessibility is a known general feature for intrinsic disorder, yet how this relates to the prevalence of intrinsic disorder along protein sequences is unclear. We address this by analyzing the fraction of predicted disordered residues for putatively solvent-exposed and solvent-buried classes, determined based on relative solvent accessibility (RSA) values obtained from DSSP program (*Materials and Methods*). The distribution of predicted disordered residues differs between classes (Fig 3, S11 and S12 Tables). The fraction of predicted disordered residues is significantly higher for terminal segments than internal ones for the solvent-exposed class, across all protein length bins. This pattern differs from patterns for the solvent-buried class where the fraction is only marginally higher at terminal segments than at internal ones for the medium and long protein length bins and, notably, significantly lower at the N-terminal segment than the remaining segments for the short protein length bin. These results may be due to incorporation of RSA values (i.e., AlphaFold-RSA) for intrinsic disorder prediction. However, this is unlikely because disorder scores are generally higher for terminal segments than internal ones (S10–12 Figs), for each of the disorder prediction approaches that do not use RSA (i.e., AlphaFold-pLDDT, flDPnn and IUPred3).

**Fig 3.**
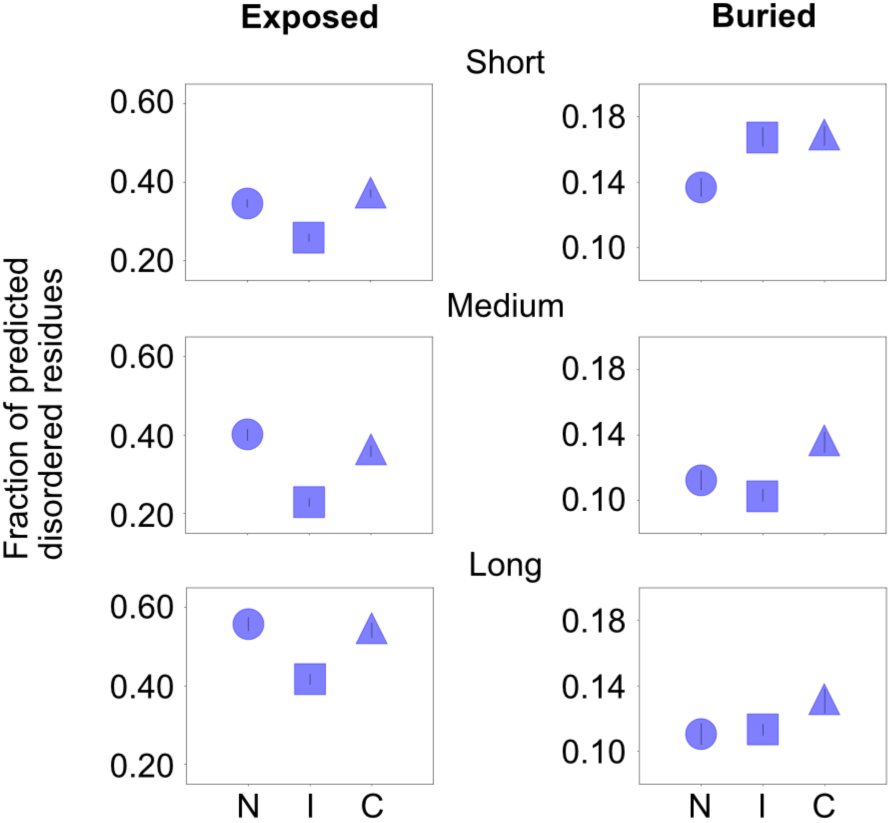
Prevalence of intrinsic disorder in putative solvent accessibility classes (exposed, buried). Residues were classified for relative solvent accessibility (RSA) values, as described in *Materials and Methods*. The exposed and buried residue classes have RSA≥0.5 and RSA<0.5, respectively. The fraction of predicted disordered residues was calculated for each class, separately for N-terminal (N), Internal (I) and C-terminal (C) segments. Error bars indicate 95% confidence intervals calculated among 1000 replicates that were created by sampling genes with replacement. n=3274, 2171 and 1737 genes for the short medium and long protein length bins. Statistical test results are presented in S11 and S12 Tables.

### Variation in the prevalence and locations of disorder functional classes

We attempt to identify primary functions of intrinsic disorder and relate them to location, and thus leverage flDPnn, which assigns protein-, RNA-, or DNA-binding, or linker functions (*Materials and Methods*). We compare those functional classes for the fraction of predicted disordered residues, focusing on putative solvent-exposed residues, which constitute the majority of predicted disordered residues and strongly capture the uneven distribution of such residues along protein sequences (results for putative solvent-buried residues are presented in S16 Fig and are generally flatter). The fraction of predicted disordered residues is highest for protein-binding followed by the DNA-binding (Fig 4, S14 Fig and S13 Table), with lower fractions for RNA-binding and linker. This pattern is generally consistent across protein segments and length bins. Across all functional classes, terminal segments have higher fractions than internal segments, with a tendency of higher fractions at N-terminal than C-terminal segment. However, the extent of difference between terminal and internal segments differs among classes (Fig 4 and S14 Table); RNA-binding and linker classes exhibit the steepest and flattest patterns, respectively. For example, in the long protein length bin, log-odds ratio (LOR) comparisons reveal greater differences between N-terminal and internal segments for the RNA-binding (0.83) than DNA-binding (0.80), protein-binding (0.59), and linker (0.47). A potential confounding factor in these analyses is that a given disordered region (or residues) may have multiple functions, limiting ability to distinguish independent and combined effects. We address this by analyzing residues assigned a single predicted function such as protein-only binding, and find patterns consistent with the initial analyses (S15 Fig and S14 Table), suggesting existence of independent relationships. Overall, the results suggest that intrinsic disorder primarily fosters interactions among proteins, and that disorder-nucleic acid interactions may be more localized at terminals.

**Fig 4.**
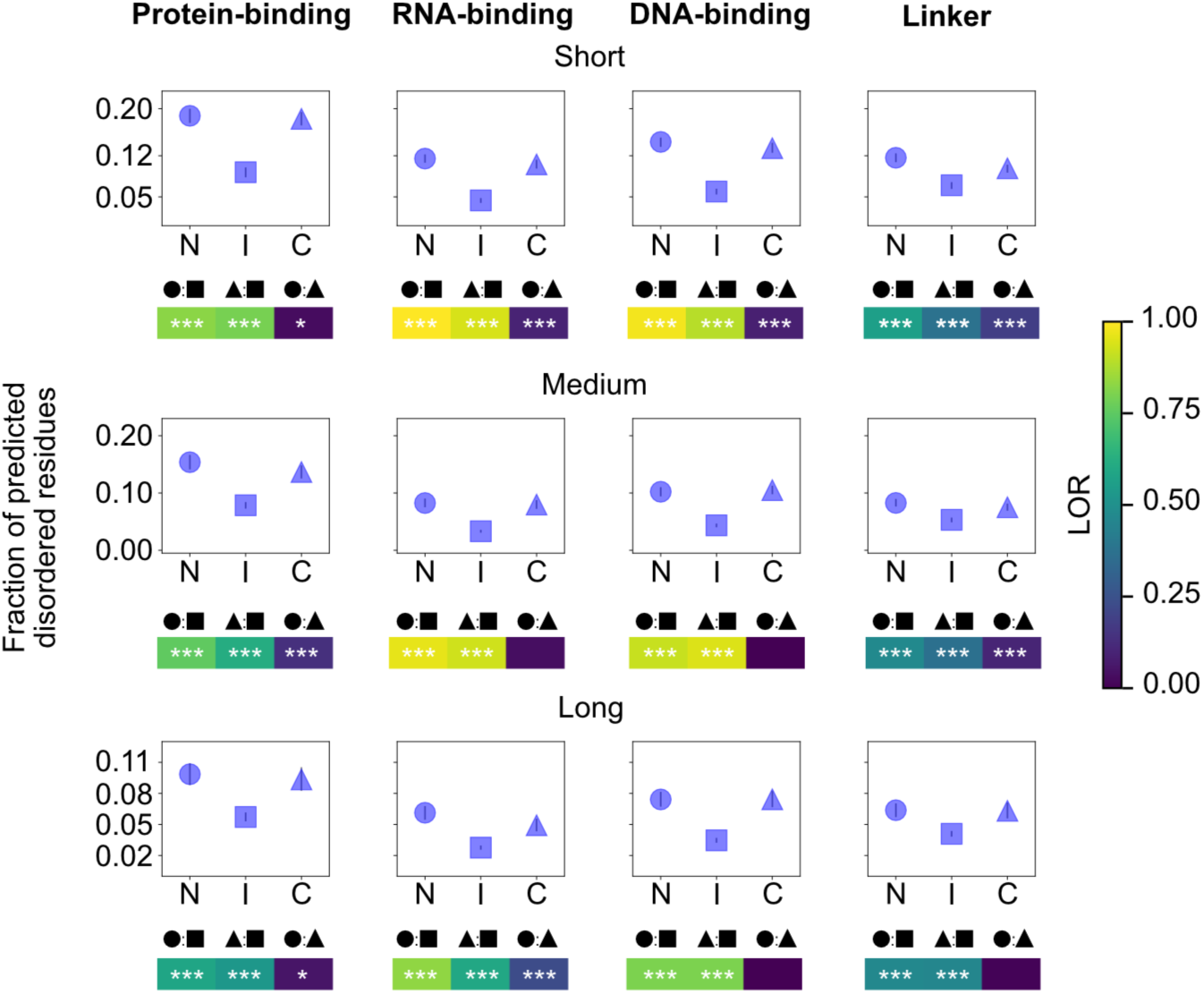
Prevalence of putative functional classes of intrinsic disorder. The fraction of predicted disordered residues was calculated for putatively solvent-exposed residues (RSA≥0.5), separately for residues assigned a function such as protein-binding (see *Materials and Methods*) and separately for N-terminal (N), Internal (I) and C-terminal (C) segments. Error bars indicate 95% confidence intervals calculated among 500 replicates that were created by sampling genes with replacement. LORs are employed for comparisons between segments. Given “A vs B” notation, LOR>0 indicates a higher fraction of predicted disordered residues for A than B. LOR<0 and LOR=0 indicates a lower fraction for A and equal fractions, respectively. n=1947, 1851 and 1558 genes for the short, medium and long protein length bins, respectively. Segment pairings were tested (Z-test) for significant differences, and the Bonferroni sequential method was employed for multiple test corrections across segment pairings and length bins (i.e., nine tests were corrected). * and *** indicate *p<*0.05 and *p*<0.001, respectively and LORs without asterisks are non-significant. Further statistical test results are presented in S13 and S14 Tables.

### Possible length and location effects in the protein connectivity-intrinsic disorder relationship

The analyses above suggest that protein-binding is a main function of intrinsic disorder, and we further investigate its relevance by comparing data for classes of protein connectivity. We obtain reports of experimentally-verified protein-protein interaction data from STRING database and use a cut-off of four interactors (i.e., the median count) to define hub and non-hub proteins (explained in *Materials and Methods*). Previous studies support greater intrinsic disorder in hub proteins, and we tested this pattern, controlling for protein length and disorder locations. The fraction of predicted disordered residues is higher for hubs than non-hubs, across all segments for the short and medium protein length bins, but not the long one (Fig 5, S17 Fig, S16 and S17 Tables). For the latter bin, hubs have a substantially higher fraction than non-hubs at the internal segment but not at terminal segments, where the fractions are comparable. A notable pattern is that the extent of difference between hubs and non-hubs is greater for the medium protein length bins than the short one (S17 Table). These observations are consistent with the idea that intrinsic disorder may have an important role in facilitating multiple functional interactions for a given protein and we find support that such a role is associated with location and length.

**Fig 5.**
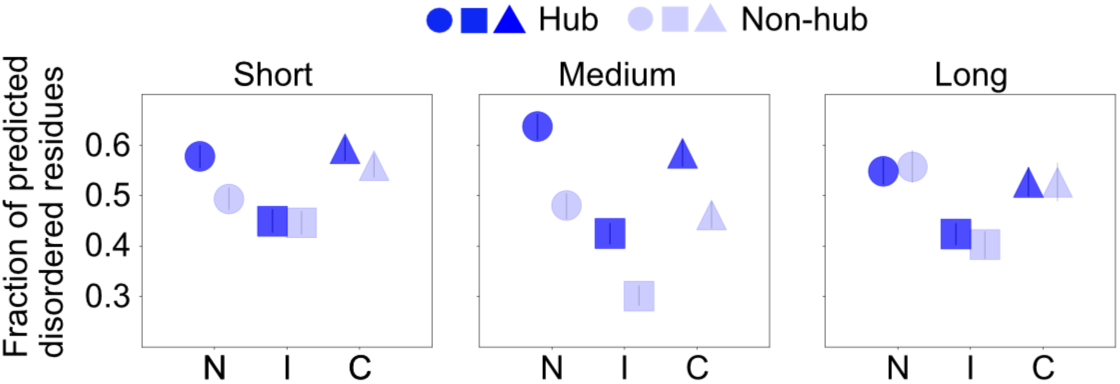
Prevalence of intrinsic disorder in protein connectivity classes. The fraction of predicted disordered residues was calculated separately for hubs (four interactors or above) and non-hubs (below four interactors), and separately for N-terminal (N), Internal (I) and C-terminal (C) segments. See *Materials and Methods* for details. Error bars indicate 95% confidence intervals calculated among 1000 replicates that were created by sampling genes with replacement. n= 943, 805 and 1132 genes for the short, medium and long protein length bins, respectively, for the hub class and n=1043, 800 and 685 for the corresponding bins for the non-hub class. Statistical test results are presented in S16 and S17 Tables.

### Distinct molecular evolutionary patterns of disordered residues

Analyzing evolutionary processes associated with intrinsic disorder can inform on function. One analysis approach is through comparisons of evolutionary rates [30–33], namely, the replacement and silent substitution rates, commonly referred to as *d_N_* and *d_S_*, but see [31]. The *d_N_* /*d_S_* ratio serves as a measure of selection pressure on the protein level, assuming that silent sites evolve neutrally or under relatively weaker selective constraints than replacement sites. *d_N_* /*d_S_* <1 and *d_N_* /*d_S_* >1would be consistent with purifying selection and adaptive evolution respectively, and *d_N_* /*d_S_* =1 would be consistent with neutral evolution. We obtain estimates of these evolutionary parameters – under a simplified implementation of the Yang and Nielsen [34] codon substitution model – for predicted disordered and non-disordered residues and separately for autosomal and X-linked genes (*Materials and Methods*).

Estimates of *d_N_* are generally higher at predicted disordered residues than non-disordered ones, across all segments, length bins, and autosomal and X-linked loci (Fig 6). An exception to this pattern is the *d_N_* at the N-terminal segment, which is comparable between predicted residue types, for the short and medium protein length bins but not the long one, where higher *d_N_* is visibly higher for predicted disordered residues than non-disordered ones. *d_N_/d_S_* values are below one for all data categories and the patterns along proteins are similar to those for *d_N_*, with no apparent major differences among segments. The differences between predicted disordered versus non-disordered residues could arise from the fact that flDPnn, AlphaFold-RSA and AlphaFold-pLDDT approaches incorporate evolutionary information for intrinsic disorder prediction. We address this issue by repeating analysis for residues that have been classified as disordered based on IUPred-short, an approach that does not incorporate evolutionary information. We observe the same general patterns (S19 Fig) as in the initial analysis; higher evolutionary rates at predicted disordered than non-disordered residues and no readily discernable *d_N_* and *d_N_/d_S_* differences among predicted disordered residues, with exception to the long protein length bin.

**Fig 6.**
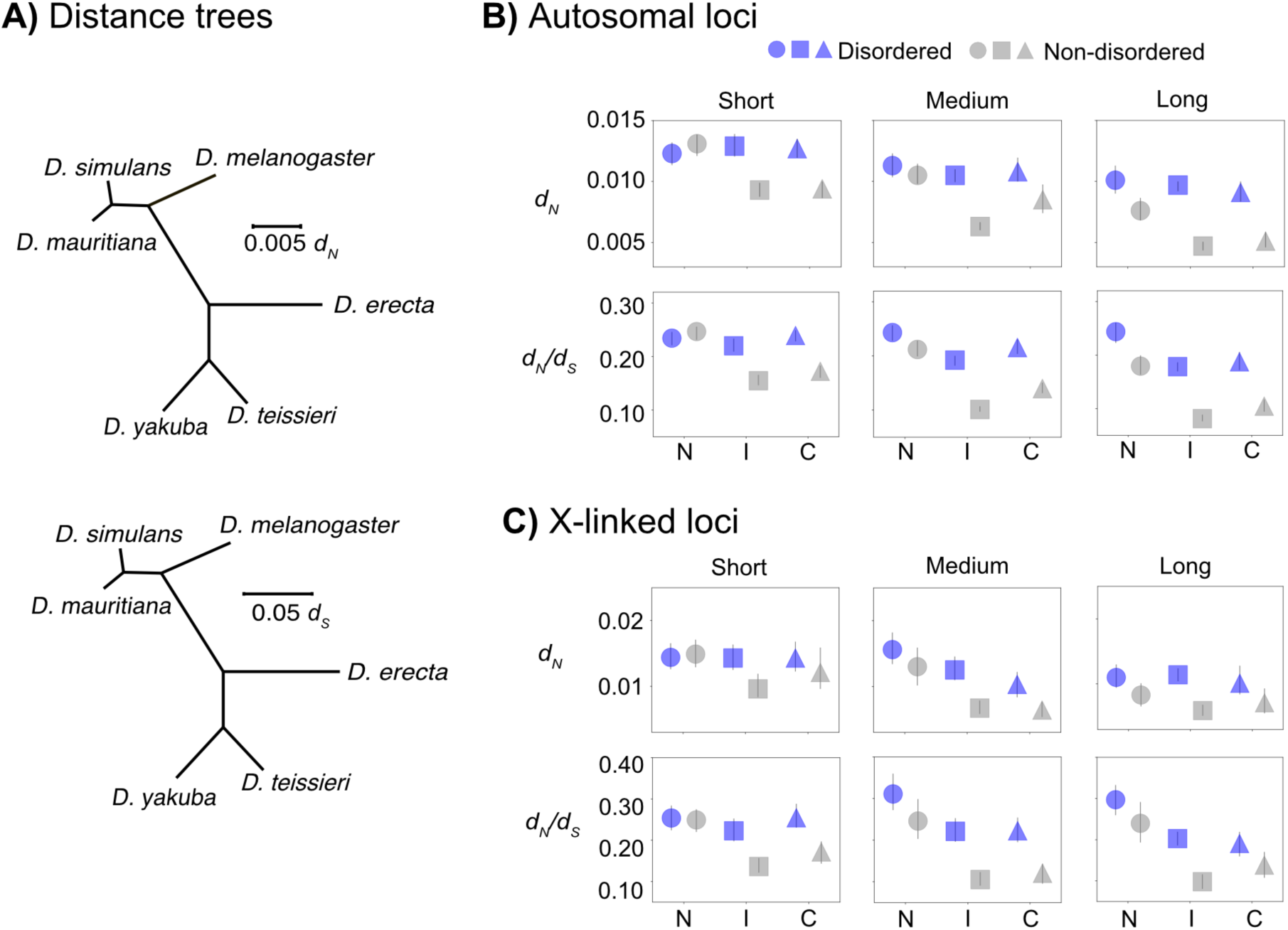
Estimates of divergence (*d_N_*) and selection pressure (*d_N_*/*d_S_*) at disordered residues. Estimates of evolutionary parameters were obtained for members of the *D. melanogaster* subgroup and those estimates for the *D. melanogaster* species branch were analyzed (see details in Materials and Methods). *d_N_* and *d_N_*/*d_S_* were calculated under the Yang and Nielsen (1998) codon substitution model implemented in codeml program, separately for N-terminal (N), internal (I) and C-terminal (C) segments. **A)** Unrooted tree, based on autosomal loci. **B)** and **C)** show *d_N_* and *d_N_/d_S_* values for autosomal and X-linked loci, respectively, for the *D. melanogaster* branch. n=2202, 1694, and 1550 genes for the short, medium and long protein length bins of the autosomal loci and n=378, 282, and 355 for the corresponding bins for X-linked loci. Error bars indicate 95% confidence intervals calculated among 100 replicates that were created by sampling genes with replacement.

### Evident but limited contribution of disorder prediction biases

The results above suggest that intrinsic disorder is more common at protein terminal segments than at internal ones, and this pattern is associated with amino acid composition and molecular interactions. We test whether these patterns are the result of biases in disorder prediction by comparing data from observed protein sequences (“Actual data”) and data from randomized versions of these sequences (“Shuffled data”). The latter is created by arbitrarily changing residue positions to decouple relationships – between amino acid sequence and location – that may be biologically relevant (*Materials and Methods*). If disorder prediction biases can fully explain the apparent biological patterns, then results from the Actual and Shuffled data should be indistinguishable. Analysis in this section uses flDPnn disorder predictions because the uneven distribution of disordered residues is observed independently for each program, and not a result of our consensus criteria or of incorporating evolutionary information. In addition, flDPnn is among the most reliable disorder prediction programs [15,35].

We compare the fraction of predicted disordered residues for Actual and Shuffled data and find substantially higher fractions for the former data across all segments (Fig 7A; S19 Table). The extent of difference between datasets is greater for comparisons between terminal segments than between internal segments (Fig 7B), with exception to the long protein sequence bin. For example, in the short protein length bin, the fraction of predicted disordered residues is ∼0.35 and ∼0.25 on the Actual and Shuffled data, respectively, for the N-terminal segment. For the internal segment, the fractions are ∼0.20 and ∼0.16 for Actual and Shuffled data, respectively. LORs for differences between Actual and Shuffled data are higher for the N-terminal (0.29) than internal (0.19) segment (*p*<0.001, Z-test. Fig 7B). These results suggest that disorder prediction biases are unlikely to explain the overall prevalence of intrinsic disorder, but the patterns suggest a role of these biases in the distribution of intrinsic disorder. Indeed, terminal segments have higher fractions of predicted disordered residues than internal segments in both Actual and Shuffled data, but with ostensibly flatter patterns in latter data (Fig 7A). We scrutinize this observation by checking whether the Actual and Shuffled data differ for LORs that compare the fraction of predicted disordered residues among segment pairs (Fig 7C). LORs from the Actual data are generally significantly higher than those from the Shuffled data (*p*<0.001, Wald test. S20 Table), but only for the short and medium protein length bins. For the long protein length bin, LORs are significantly higher for the Shuffled data. In addition, the fraction of predicted disordered residues is higher for the N-terminal than C-terminal segments across all protein length bins in the Actual data (Fig 7A). We observe a similar pattern in the Shuffled data but only for the short protein length bin, and for the remaining bins, the fraction is substantially lower for the N-terminal than C-terminal segment, and clearly differs from patterns of the Actual data. These observations further support the notion that disorder prediction biases can only partially explain the distribution of disordered residues, and indicate that the explanatory scope would depend on location and length. We emphasize that the biases cannot explain differences between Actual and Shuffled data for distribution shapes (Fig 7A).

**Fig 7.**
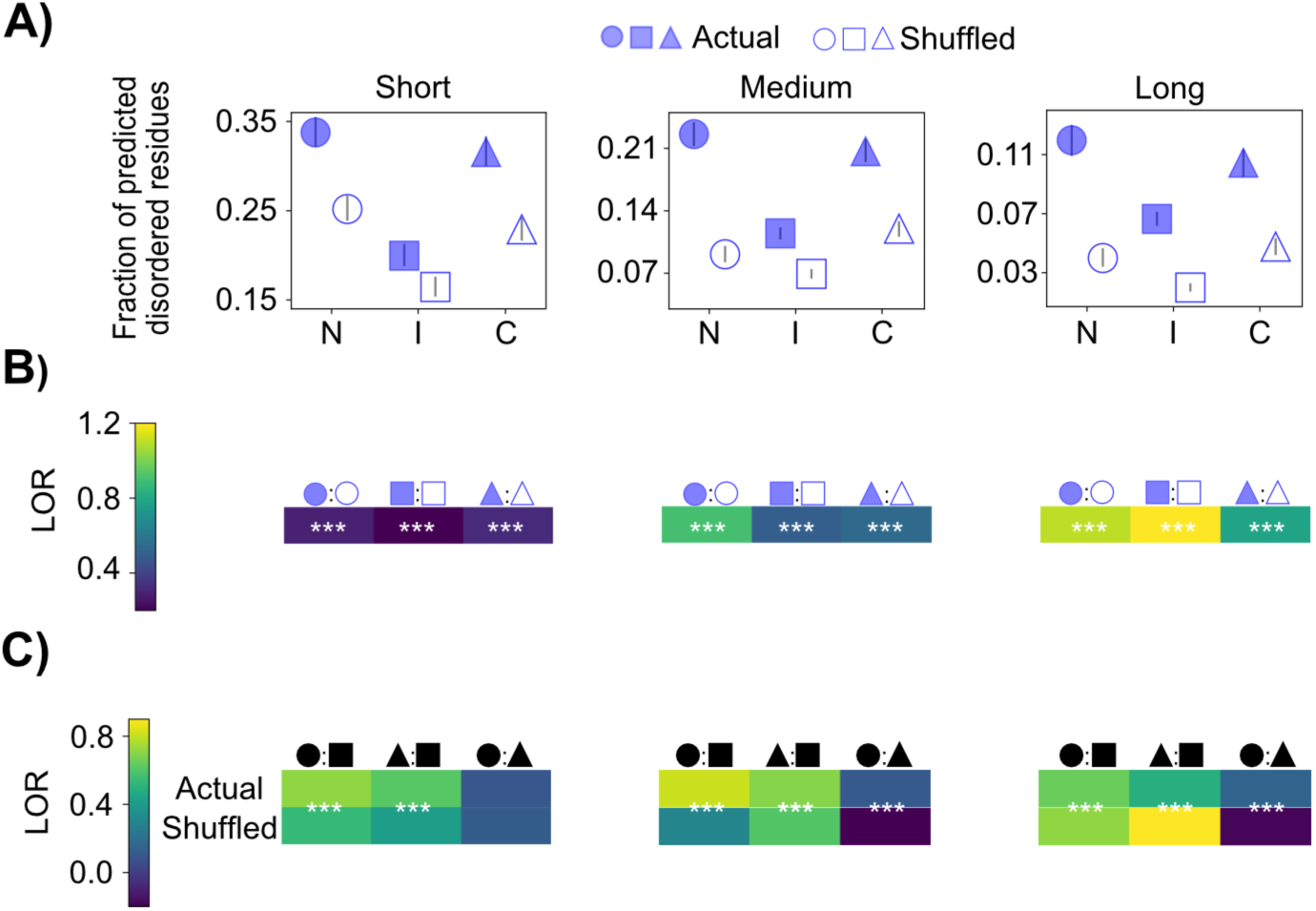
Actual and Shuffled data comparisons for the fraction and distribution of predicted disordered residues. flDPnn was employed for intrinsic disorder prediction in *D. melanogaster* protein sequences (“Actual” data) and in randomized versions of those protein sequences (“Shuffled”data). The resulting predictions were tested for differences between data types. **A)** Prevalence of intrinsic disorder along Actual and Shuffled protein sequences. Data are shown separately for N-terminal (N), Internal (I) and C-terminal (C) segments. n=2005, 1981 and 1956 genes for the short, medium and long protein length bins, respectively. **B)** LORs for differences between Actual and Shuffled data for the fraction of predicted disordered residues at terminal and internal segments. LORs>0 and <0 indicate higher and lower fractions, respectively, in the Actual data. The data types were tested (Z-test) for significant differences. **C)** LORs for differences between segments for the fraction of predicted disordered residues. Interpretation of LOR values follows the same logic as in panel B. The data types were tested (Wald test) for significant differences. In panels B and C, *** indicates *p*<0.001. Absence of asterisks indicates non-significance. Detailed statistical test results are shown in S19 and S20 Tables.

We test whether biases in disorder prediction can explain the apparent regional heterogeneity in amino acid composition. Actual and Shuffled data differ for the frequency distributions of many amino acids including Lys, Leu, Pro, Ser, Tyr and Val. (Fig 8, S21–23 Figs. S21 and S22 Tables). For example, in the short protein length bin, the frequency of Lys is higher at terminal segments than internal segments in the Actual data but not the Shuffled data, where the opposite pattern is observed (S21 Fig). In the medium protein length bin, Lys frequency is higher at terminal than internal segments in the Actual data, which differs from the Shuffled data where Lys frequency at the N-terminal, and not at the C-terminal, is higher than the frequency at the internal segment (S22 Fig), and we observe similar patterns in the long protein length bin. Another notable aspect of these observations is that the frequency distributions of Lys at non-disordered residues differ between Actual and Shuffled data, a pattern challenging to explain as an effect of disorder prediction bias. Such an effect is also an unlikely explanation for the patterns of Leu, Pro, Ser, Tyr, and Val (S21-S23 Figs). For other amino acids, clear support exists for a role of disorder prediction biases, but the explanatory scope depends ostensibly on length and location.

**Fig 8.**
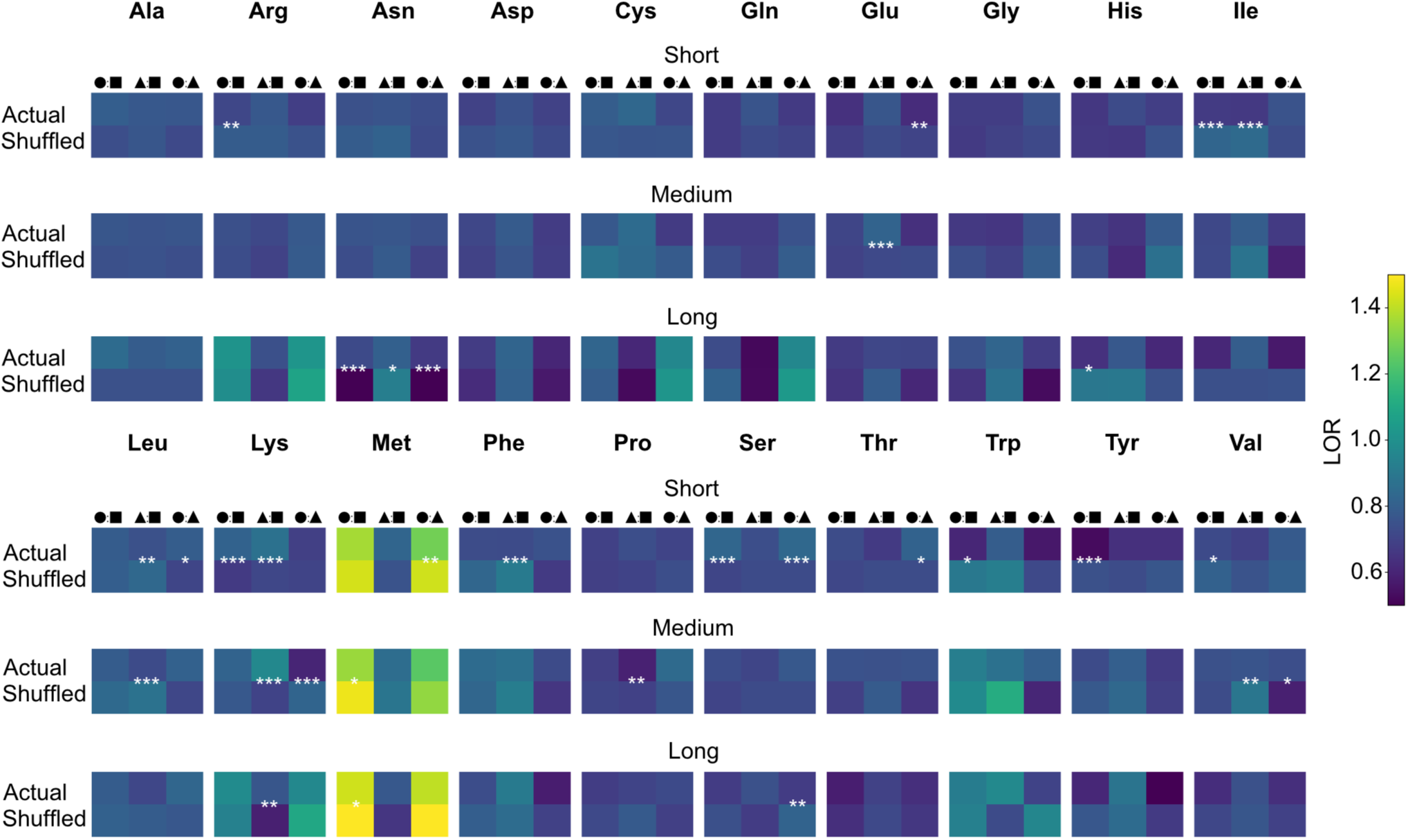
Actual and Shuffled data LOR comparisons for amino acid frequency differences among segment pairs. flDPnn was employed for intrinsic disorder prediction in *D. melanogaster* protein sequences (“Actual” data) and in randomized versions of those protein sequences (“Shuffled”data). Amino acid frequencies of the resulting predicted disordered residues are compared among N-terminal (circle), Internal (square) and C-terminal (triangle) segments, separately for each data type. Given “A:B” symbol, LOR>0 and LOR<0, indicate a higher and lower frequency, respectively, for segment A. Wald test was employed to check whether Actual and Shuffled data exhibit the same extent of amino acid frequency differences among segment pairings (i.e. LOR differences). The Bonferroni sequential method was employed for multiple test correction across segment pairings and protein length bins (i.e., nine tests were corrected for each amino acid). Asterisks specify *p*<0.05 (*), *p*<0.01 (**) or *p*<0.01 (***). Absence of asterisks indicates non-significance. n=2005, 1981 and 1956 genes for the short, medium and long protein length bins, respectively. Plots of amino acid frequency trends are presented in S21–S23 Figs and statistical test results are provided in S21 and S22 Tables.

We compare Actual and Shuffled data for their amino acid frequency distributions by employing Wald tests that contrast LORs. These LORs reflect differences among segment pairs for amino acid frequencies (Fig 8). LORs differed significantly for ten amino acids. For Lys, these differences are observed consistently across protein length bins, in particular, for comparisons of C-terminal versus internal segment pairs. For the remaining amino acids, we observe significant differences but mostly for one or two protein length bins. Interestingly, we did not observe strong support that Met frequencies distributions are attributable to disorder presence, in contrast to the analyses based on the consensus criteria. Met distributions are highly comparable between Actual and Shuffled data, and LORs (N-terminal vs Internal) differ significantly only for the medium and long protein length bins. Overall, these results support a role of biases in disorder prediction in regional heterogeneity, including for Met, but reveal that the biases cannot fully explain the heterogeneity. We note that the evidence for regional heterogeneity is clear, even in experimental data (S26 and S27 Figs), and note many instances where the heterogeneity can plausibly be attributed to intrinsic disorder presence.

We test whether disorder prediction biases underlie the relationship between disorder functions and location, and thus compare the Actual and Shuffled data for the fraction of predicted disordered residues for putative functional classes. The fractions are significantly higher in the Actual data than the Shuffled data across all functional classes and segments (Fig 9, S23 Table). Interestingly, these differences scale with protein length. For example, in the protein-binding class, LORs for the difference between N-terminal segments of Actual versus Shuffled data are 0.27 and 0.99, respectively, for the short and long protein length bins (Fig 9A and S23 Table). Across all functional classes, LOR differences (Actual versus Shuffled data comparisons) for terminal segment comparisons tend to be greater than those for internal segment comparisons. This pattern is observed in the short and medium protein length bins, but an opposite pattern is observed in the long protein length bin.

**Fig 9.**
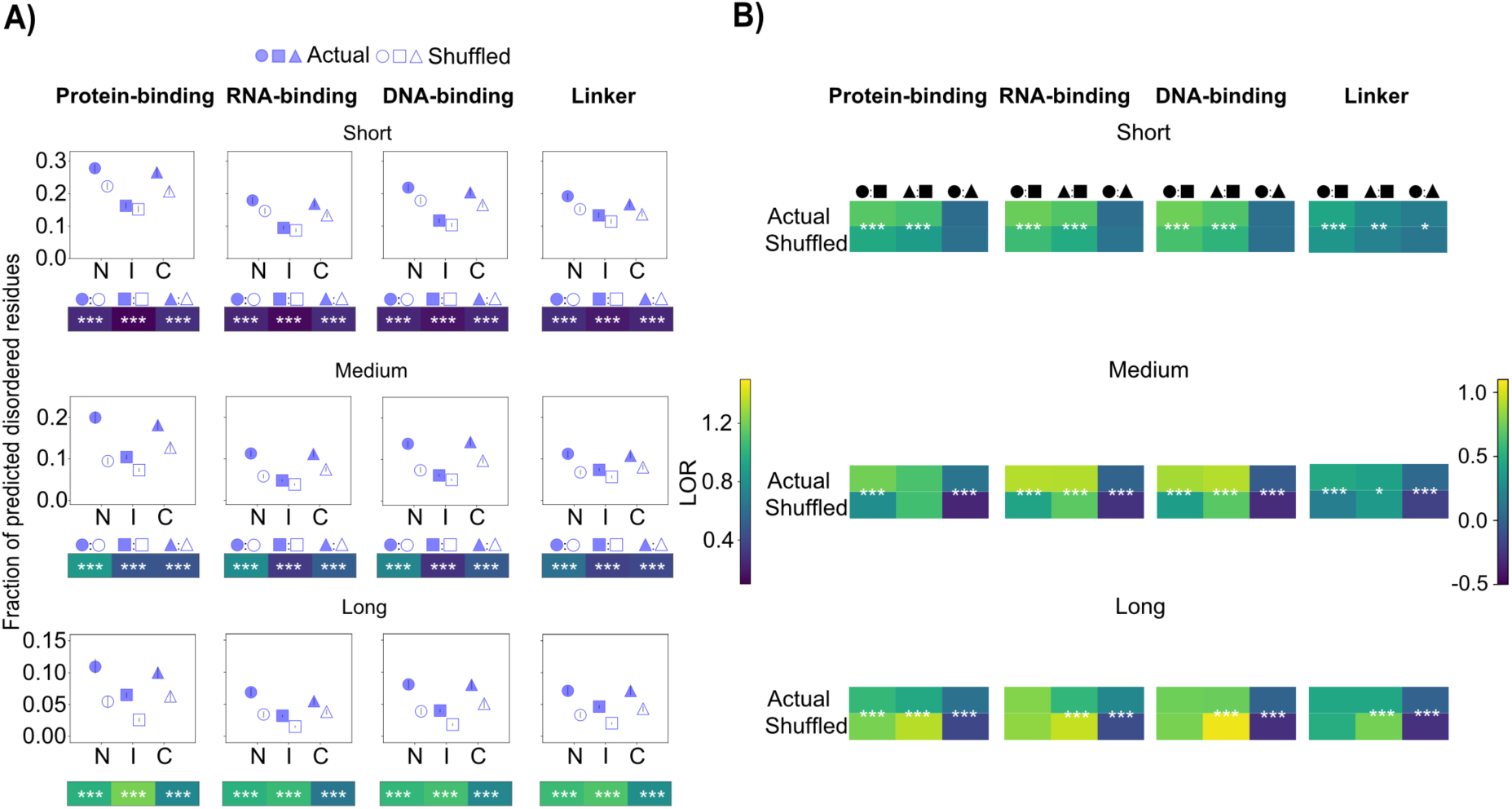
Actual and Shuffled data comparisons for the prevalence and location of functional classes of intrinsic disorder. flDPnn was employed for intrinsic disorder function prediction in *D. melanogaster* protein sequences (“Actual” data) and in randomized versions of those protein sequences (“Shuffled”data). The fraction of predicted disordered residues was compared among functional classes (See details in *Materials and Methods*), and among N-terminal (N or circle), Internal (I or square) and C-terminal (C or triangle) segments. **A)** Prevalence of disordered residues for each functional class. Error bars are 95% confidence intervals calculated among 500 replicates that were created by sampling protein sequences with replacement. n=2005, 1981 and 1956 genes for the short, medium and long protein length bins. LORs were employed for comparing Actual and Shuffled data for each segment. Given “A:B” symbol, LOR>0 indicates a higher fraction of predicted disordered residues for A than B. LOR<0 and LOR=0 indicate a lower fraction for A and equal fractions, respectively. Actual and Shuffled data were tested (Z-test) for significant differences for the fraction of predicted disordered residues. All differences are significant at *p*<0.001. **B)**. LOR for comparisons between segment pairings. Wald test was employed to check whether Actual and Shuffled data exhibit the same extent of difference for each segment pairing (i.e. LOR differences). *, ** and *** indicate *p*>0.05, *p*>0.01 and *p*>0.001, respectively. Absence of asterisks indicates non-significance. In both panels, the Bonferroni sequential method was employed for multiple test corrections among segments and protein length bins (i.e., nine tests were corrected). Statistical tests results are presented in S23 and S24 Tables.

We checked whether Actual and Shuffled data differ for the distribution of disordered residues among each functional class by comparing LORs calculated for segment pairings (Fig 9B and S24 Table). Across all putative functional classes, the LORs tend to be significantly higher in the Actual than Shuffled data, but only in the short and medium protein length bins. In the long protein length bin, the LORs tend to be significantly lower in the Actual data. For instance, in the protein-binding class, LORs comparing N-terminal versus internal segments are 0.69 and 0.47 for the Actual and Shuffled data, respectively, in the short protein length bin (*p*<0.001 for a Wald test. Fig 9B, S24 Table). In the long protein length bin, the corresponding LORs are 0.57 and 0.78 for the Actual and Shuffled data, respectively. LORs for the RNA-binding class tend to be higher than those for other functional classes.

We also test whether disorder prediction biases can explain the relative prevalence of disorder functions. In the Actual and Shuffled data, the protein-binding class tends to be significantly more prevalent than the remaining functional classes, across all protein length bins (Fig 10 and S25 Table), and the RNA-binding class tends to be least prevalent. For example, in the short protein length bin, we observe positive LORs when comparing the protein-binding class relative to each of the remaining classes, and observe, in most cases, negative LORs when comparing the RNA-binding class relative to other functional classes (Fig 10 and S25 Table). The similarity of patterns between the Actual and Shuffled data suggests that disorder prediction biases can explain the relative prevalence of disorder function. However, the LORs in the Actual data tend to differ significantly from those in the Shuffled data, in particular in the short and medium protein length bins, suggesting differential enrichment of functional classes whereby the protein-binding and RNA-binding classes tend to be the most and least enriched, respectively, relative to other classes.

**Fig 10.**
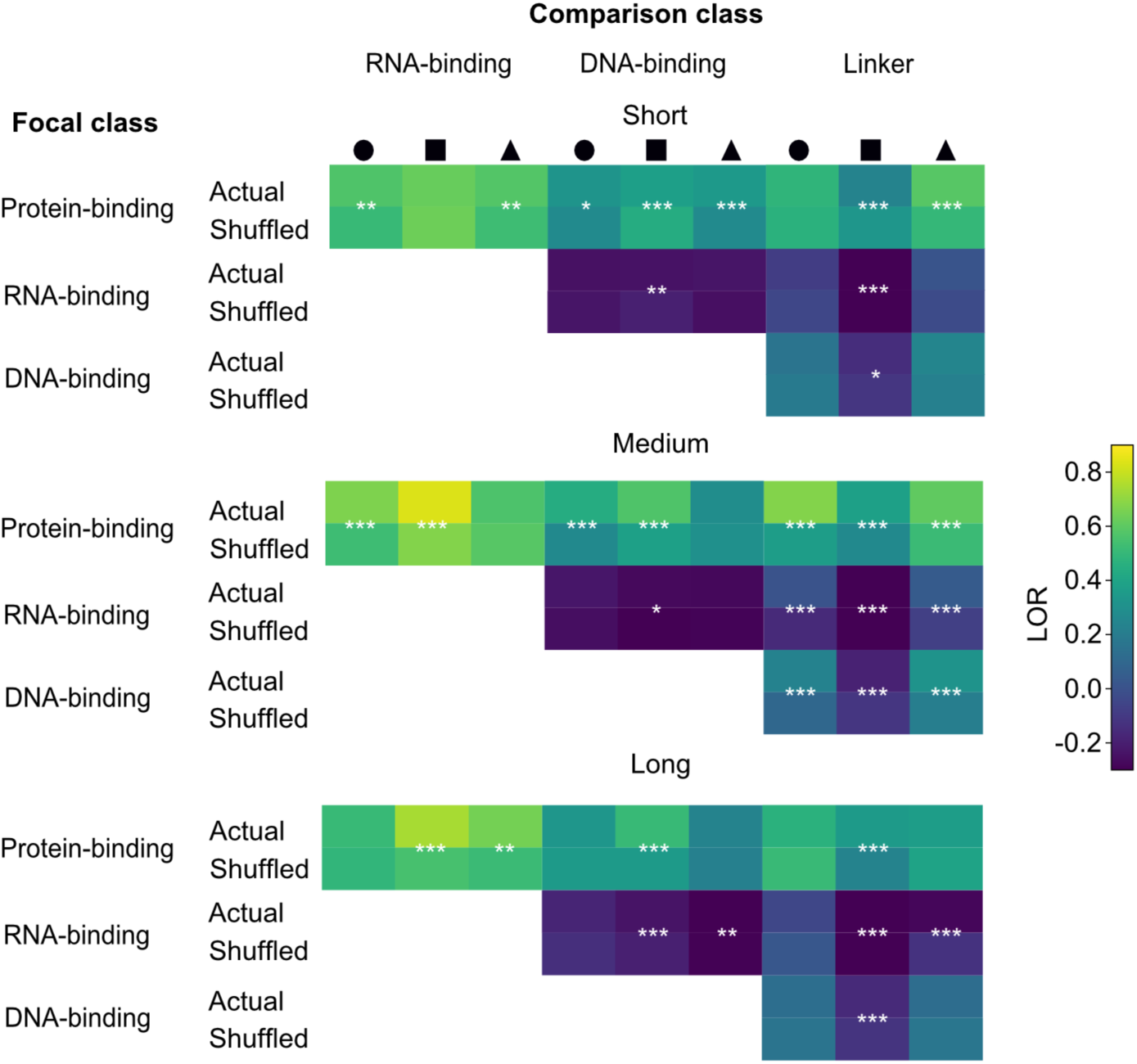
Actual and Shuffled data comparisons for the relative prevalence of intrinsic disorder functions. flDPnn was employed for intrinsic disorder function prediction in *D. melanogaster* protein sequences (“Actual” data) and in randomized versions of those protein sequences (“Shuffled”data). LORs were used to test for differences between “focal” and “comparison” classes, separately for N-terminal (circle), Internal (square) and C-terminal (triangle) segments. LOR>0 indicates a higher fraction of predicted disordered residues for the focal class than the comparison class and LOR<0 and LOR=0 indicate a lower fraction and equal fractions, respectively. Wald test was employed to check whether Actual and Shuffled differ for LORs. *, ** and *** indicate *p*>0.05, *p*>0.01 and *p*>0.001, respectively. Absence of asterisks indicates non-significance. The Bonferroni sequential method was employed for multiple test corrections among segments and protein length bins (i.e., nine tests were corrected). n=2005, 1981 and 1956 genes for the short, medium and long protein length bins. Detailed statistical tests results are provided in S25 Table.

## Discussion

This study investigated the prevalence of intrinsic disorder in biophysical and functional contexts, with a general aim of testing the disorder-function paradigm. We demonstrate the utility of employing single and consensus approaches for intrinsic disorder prediction, as well as employing descriptive and enrichment statistics to address – at a genome-wide scale – the relevance of intrinsic disorder in function and evolution. A key underlying assumption in the analyses is that biological relevance can be inferred from variation in the prevalence of intrinsic disorder – and related properties – along protein sequences. We find new support for the paradigm; disordered regions are likely unevenly distributed along protein sequences in a manner associated with amino acid composition, solvent accessibility and function. Evolutionary processes may generally differ between disordered and non-disordered regions, but may be similar along protein sequences. Biases of disorder prediction approaches do not appear to be a sufficient explanation of the patterns, but can be a considerable factor that complicates interpretations, especially with respect to the relevance of amino acid composition and the main functions of intrinsic disorder. We summarize the significance and limitations of this study in the sections below.

### Prevalence of intrinsic disorder along protein sequences

Efforts to address the locations of intrinsic disorder often focus on terminal residues [e.g., 24]. Here, we considered the full length of proteins, and observed support for greater prevalence of disordered regions at protein terminals than internal regions in *D. melanogaster* (Fig 1), a pattern similar to that Lobanov et al., [36] who analyzed data from PDB database. We found that this pattern is mainly caused by higher prevalence of intrinsic disorder at the first/last 10–20 residues (S4A Fig), is generally robust to protein length (Fig 1 and S3 Fig), is a feature of the disorder score distribution along protein sequences (S10–S12 Figs), and is observed in many other organisms including human and *E. coli* (S28 and S30 Figs). This apparent location dependence of intrinsic disorder plausibly reflects biases from the prediction programs, but the biases do not appear to be a sufficient explanation for the pattern, at least in *D. melanogaster*. Evaluation of the biases in *D. melanogaster* revealed that intrinsic disorder is likely distributed in a manner that differs, to varying extents, from predictions on randomized data. Results from experimental data, despite being scant, show some consistency with location dependence in *D. melanogaster* and the non-*Drosophila* species (S25 and S30 Figs). The apparent location dependence of intrinsic disorder may reflect a biological signal. Since many disorder prediction approaches, including those employed in this study, incorporate information on experimentally-determined protein structures/disordered regions, the resulting predictions may thus reflect genuine biological signals that are exaggerated or swamped. Addressing this issue may require evaluating positional biases among disorder prediction programs, as well as more experimental data.

### Amino acid composition and intrinsic disorder

We observed compositional differences between predicted disordered and non-disordered residues (Fig 2), consistent with many previous studies across multiple taxa including *D. melanogaster* [13,14,25,37]. Such differences are generally expected due to known compositional trends of intrinsic disorder, but the fact that these patterns can differ along proteins and can differ among protein length categories was not necessarily expected. Our analyses also revealed support for regional compositional heterogeneity that can plausibly be attributed to intrinsic disorder presence and highlighted the relevance of disorder prediction biases.

The distribution of Met frequencies was notable throughout the amino acid frequency analysis based on the consensus criteria for disorder prediction. Met frequencies were higher at the N-terminal segment than the internal and C-terminal segments, at predicted disordered residues only, and the corresponding distribution at non-disordered residues was comparatively flat, which suggested a possible role of intrinsic disorder associated with Met presence. We found that initiator Met underlies the high Met frequencies at disordered residues in N-terminal (S8 Fig). This observation, as well as observations in randomized data, suggest an effect of disorder prediction bias and thus complicate interpreting the relationships between intrinsic disorder and genome-wide frequencies of Met. Critically, however, the findings do not rule out a functional role between initiator Met and intrinsic disorder. First, the distribution of Met frequencies along medium and long proteins differs, although marginally, between Actual and Shuffled data (e.g., Fig 8 and S21–S23 Figs). Second, analysis of experimentally-determined disordered regions revealed that Met frequencies are substantially higher at N-terminal segment than at the internal and C-terminal segments (S26 and S27 Figs), a pattern that is observed when data from multiple organisms are pooled [36]. Third, initiator Met and intrinsic disorder can be plausibly related, at least, with respect to post-translational modifications. Excision of initiator Met can be essential for protein stability, subcellular localization and/or further modifications [38–42]. The process is thought to occur in many organisms including *D. melanogaster*, and is catalyzed by Met aminopeptidase (MetAP) enzyme for which activity is associated with the identity of the second residue position. Future investigations may benefit from analyses of Met frequencies at the first residue while controlling for the amino acid identity at the second residue. Oxidation is another plausible link between initiator Met and intrinsic disorder. Met oxidation can be important for regulating molecular interactions, and is enriched in proteins that are sensitive to redox conditions including disordered regions [43–45]. Whether this pattern is associated with initiator Met remains is unclear. Experimental investigations relating initiator Met and intrinsic disorder may provide further insight on functional relevance. Despite these limitations, regional heterogeneity in amino acid composition – including for Met – is clear from observations of this study. Lys is among the compelling cases that seem to associate amino acid compositional heterogeneity and intrinsic disorder. The frequency ranks (among segments) of Lys are inconsistent between predicted disordered and non-disordered residues, a pattern robust to the disorder prediction approach (e.g., compare Lys data in S5–7 Figs). Lys frequency ranks are also inconsistent between Actual and Shuffled data (S21–S23 Fig). Such results tether amino acid composition and intrinsic disorder in a manner difficult to explain by proteome-wide patterns and/or disorder prediction biases. The associations may reflect amino acid compositional requirements of disordered binding regions along protein sequences, consistent with previous support relating amino acid composition to specific functions [reviewed in 1,36]. For instance, disordered regions containing polyelectrolytes (e.g., stretches of Glu, Ser, Lys) are reported to have roles in molecular interactions associated with transcription and chromatin formation.

### Solvent accessibility and intrinsic disorder

Previous studies have established deterministic relationships between solvent accessibility and intrinsic disorder; high solvent accessibility is a known general feature of disordered regions as it allows for increased structural flexibility [46]. We attempted to clarify relationships between solvent accessibility and intrinsic disorder by analyzing the distribution of disordered residues along proteins among classes of solvent accessibility. Interestingly, we found that the apparent location dependence of intrinsic disorder seems associated with solvent accessibility; disordered regions are likely more common near terminal than internal segments, in a solvent-exposed context but not in a buried one, across multiple taxa (Fig 3, S13 Fig), where location dependence is weaker or non-existent. Overall, these observations possibly hint that solvent-exposure is an essential property for functions of most disordered regions – at terminal disordered regions in particular – and that putative functions of intrinsic disorder generally involve intermolecular interactions.

### Possible main functional roles of intrinsic disorder

The putative functional roles of intrinsic disorder are often studied with respect to protein-protein interactions and/or gene ontology (GO) terms [19,24,25,47]. Here, we delved into the relationship between intrinsic disorder and function by focusing on the main functions of intrinsic disorder, taking location into account. First, we analyzed the prevalence of multiple classes of disorder functions such as protein-binding and RNA-binding and the “single-function” classes and found evidence that most disordered regions are involved in binding rather than linking domains (Fig. 4 and S14–16 Tables). Among the putative disorder binding classes, protein- and RNA-binding appear to be the most and least common, respectively. These patterns are consistent among *D. melanogaster*, human and *E. coli*. Disorder prediction biases complicate interpretations because patterns can be similar between Actual and Shuffled data (Fig 9), but the biases do not appear to explain disorder prevalence within each functional class; the fraction of predicted disordered residues is consistently higher in observed than randomized data, across the protein length bins. The biases, however, account for the relative prevalences among binding and linker functions, but to limited extents depending on the protein length bin, function and location. Future efforts to address primary functional roles of intrinsic disorder may benefit from employing multiple programs for function prediction.

We attempted to relate disorder locations to function by analyzing the distribution of disordered residues for putative functional classes. Putative binding functions, compared to linker functions, seem to contribute more strongly to the unevenness of disorder locations (Fig 4 and S14 Table), and RNA-binding and DNA-binding appear to be main factors. These patterns can plausibly be explained by disorder prediction biases, but with limited explanatory power; the extent of unevenness in disorder locations differs significantly between the Actual and Shuffled data (Fig 9 and S24 Table), a pattern observed consistently across protein length bins (but with some exceptions to the segment pairs analyzed). We found support that these patterns plausibly result from individual effects of a given function (e.g., protein-only binding), rather than combined effects, which had not been observed in previous studies. A challenge is to address how functions (e.g., ion-binding) other than the ones employed in this study relate to the prevalence and locations of intrinsic disorder.

We found support that intrinsic disorder is more common in protein hubs than non-hubs, consistent with the view that intrinsic disorder allows protein hubs to interact with multiple structurally diverse partners [9,47]. An unaddressed aspect is whether/how this pattern relates to the locations of intrinsic disorder, and we found that protein hubs have a higher fraction of disordered residues than non-hubs (Fig. 5 and S16 Table), in a manner that is associated with location and length. The challenge remains to understand relationships between connectivity and intrinsic disorder, considering interactions of the latter with non-proteins.

### Relating evolutionary processes to disorder locations

Relationships between protein evolution and intrinsic disorder have been studied in many organisms [27,28,49], but have not distinguished disorder locations. Here we attempted to clarify relationships by analyzing *d_N_* and *d_N_*/*d_S_*, separately for terminal and internal segments. *d_N_* and *d_N_*/*d_S_* are generally higher at predicted disordered residues than non-disordered ones, which is consistent with previous studies, but are similar when N-terminal is examined. These results suggest that faster protein evolution, and possibly relaxed constraints, may be a common feature of intrinsic disorder, and that regional differences in evolutionary rates can be a factor in the evolutionary rates of disordered residues (Fig 6 and S18 and S19 Figs). Understanding relationships between disorder functions and protein evolution will benefit from controlling for disorder functions, and known factors in evolutionary parameter estimation.

## Limitations

Despite the relatively detailed analysis, predicting intrinsic disorder, establishing causal relationships, and understanding evolutionary processes at disordered regions are among the main limitations of this study and are discussed in the next sections.

### Uncertainties in intrinsic disorder prediction

Quantifying the prevalence of disordered residues has been an ongoing issue in intrinsic disorder research and was among the goals of this study. Here, we developed consensus criteria sets to identify disordered residues with strong support, but found that disorder prevalence can be highly sensitive to criteria set. For example, the fraction of predicted disordered residues in *D. melanogaster* is 0.414 and 0.117 under non-strict and strict implementations of the consensus criteria sets developed in this study (S2 Fig). The former value is within the range previously predicted for multicellular eukaryotes (∼0.35-0.45. [50,51]), but differs from the values predicted for *D. melanogaster* in other studies (Panda and Tuller, [24] reported ∼0.280 and Ward et al., [52] reported 0.216). The value under the strict consensus criteria set is likely conservative given the low recall rate on experimental data (S2 Fig and S3 Table). Nonetheless, the inconsistencies among studies, including the current one, highlight the withstanding challenge to quantify disorder prevalence accurately. Large-scale experimental determination of intrinsic disorder may be essential to address this challenge.

Accurate prediction of the full-length of disordered regions is another ongoing challenge even when consensus approaches are employed (e.g., [53]). The consensus approaches employed in this study are not designed/optimized to predict entire disordered regions, instead, the predictions are residue-level and mitigate the uncertainty of determining appropriate lengths of disordered regions. Hence, these consensus approaches may not be appropriate analyzing a few proteins and/or disordered regions. Future studies may benefit from employing strategies/models that incorporate length distributions of experimentally-determined disordered regions.

### Establishing causal relationships

This study attempted to address the factors underlying disorder prevalence and locations, by analyzing the distribution of amino acid frequencies at predicted disordered residues, and by comparing the fraction of predicted disordered residues among various putative biophysical and functional classes. The resulting observations provided clues about possible causal relationships, but strong conclusions remain challenging. Controlling for other factors (e.g., intra-molecular interactions, base composition), and experimental analyses may yield further insights on cause. DSSP program [54] assumes a static structure for solvent accessibility estimation, and future investigations may benefit from estimates that integrate structural dynamics over time.

### Identifying primary functions of intrinsic disorder

This study attempted to determine the primary functional roles of intrinsic disorder, and thus focused on general classes such as “protein-binding”. An alternative approach would be to relate intrinsic disorder with motifs that have known functions such as protein targeting signal motifs. These signals constitute different classes of peptide motifs such as signal peptides [55,56], nuclear localization signals [57], and mitochondrial presequence [58], and involve various critical processes including secretory pathways and organellar targeting. Many of these motifs are composed of charged and/or polar amino acids, likely rendering a dynamic nature. Indeed, many studies, across multiple taxa including *Drosophila*, support roles of intrinsic disorder in functions of signal peptides [59–62] and nuclear localization signals [63,64]. If presence of protein targeting signals is a factor in intrinsic disorder prevalence, then most disordered regions should contain such signals. Signal peptides are of particular interest to understand possible functions of N-terminal disordered regions because, by definition, they are exclusive to the N-terminal, unlike other protein targeting signals. A signal peptide is composed of a short motif, 16–30 residues long [56]. A subset of the signal peptide motif, 1–5 residues long, typically contains the initiator Met and charged residues such as Lys, Arg and His. This subset serves roles in interactions with lipid membranes, signal recognition and protein transport, and its charged composition could suggest intrinsic disorder presence. Hence, analysis of signal peptides may allow testing the biological significance of N-terminal disorder, and its relationship to initiator Met, at a genome-wide scale. Mechanistically, a potential role of N-terminal disorder would be to foster recognition of signal peptidase, which catalyzes the cleavage of signal peptide.

### Uncertainties in evolutionary analysis

Non-stationary evolutionary processes can affect divergence estimation (e.g., [65]) but were not considered. Our primary goal was to test patterns, rather than determine specific values (e.g., *d_N_*). Potential confounding effects of ignoring non-stationary evolution may cancel out, if these effects are similar along proteins. We tested whether this assumption is plausible by obtaining nucleotide substitution rate estimates under a model that accounts for non-stationary evolutionary processes “GTR-NH_b_” and comparing these rates for the second codon position of predicted disordered and non-disordered residues. The substitution rates are higher for predicted disordered residues than non-disordered ones, with no major discernable differences among locations (data not shown), which is consistent with the pattern based on *d_N_* and *d_N_/d_S_* estimates. As the analyses assume site independence in evolution, future work may benefit from accounting for possible effects of context-dependence, given the existing support in *Drosophila* (e.g., [66]).

Alignment quality and tests of natural selection can both be sensitive to the alignment algorithm employed (e.g., [67–69]). Terminal regions can be more prone to misalignment than internal regions, ostensibly, owing to high divergence. This study attempted to mitigate alignment quality issues by employing a dataset of closely related species, which is known to allow for improved alignability compared to datasets with higher divergence (e.g., [70,71]). In addition, the alignment algorithm we employed, E-INSI, is designed to address gaps and regions that are challenging to align, a feature of disordered regions, by employing an iterative approach that builds multiple sequence alignments based on the consistency of local pairwise alignments, a process that incurs penalties for gap and indel presence. Such iterative refinement algorithms tend to perform more reliably than methods that are non-iterative and do not check alignment consistency ([72,73]). Analyzing the subset of sites that are consistently aligned across multiple algorithms (e.g., [74]) will allow testing robustness of the evolutionary analysis to alignment quality.

## Materials and methods

### Protein sequences and structures

We obtained protein sequence and predicted structures from publicly available databases. Coding sequences (CDS) for members of the *Drosophila melanogaster* subgroup were obtained from FlyBase or NCBI and translated, and 3D protein structure predictions for *D. melanogaster* were obtained from AlphaFold Protein Structure database. We mapped data from FlyBase and AlphaFold Protein Structure database by comparing protein sequence identity. Note that unless indicated otherwise, protein sequence counts correspond to gene counts (one protein sequence per gene is used).

We extracted CDS of *D. melanogaster* annotation 6.62 from “dmel-all-CDS-r6.62.fasta.gz” file downloaded on April 14, 2025 (http://flybase-ftp.s3-website-us-east-1.amazonaws.com/genomes/Drosophila_melanogaster/dmel_r6.62_FB2025_01/index.html), and extracted CDS of the following *Drosophila* species annotations from “cds_from_genomic.fna” downloaded on April 14, 2025 (https://www.ncbi.nlm.nih.gov/datasets/genome): *D. simulans* (103), *D. mauritania* (100), *D. yakuba* (102), *D. teissieri* (100), *D. erecta* (103)*, D. elegans* (102), *D. ficusphila* (102) and *D. kikkawai* (GCF_030179895.1-RS_2024_12). For each species, we used the longest CDS isoform for each gene and translated such CDS using the standard genetic code. These CDS and the resulting protein sequences were primary inputs for evolutionary analysis and the ones from *D. melanogaster* were also inputs for both prediction of intrinsic disorder and other protein properties. The following CDS counts were processed: *D. melanogaster* (13,937), *D. simulans* (14,109), *D. mauritania* (13,891), *D. yakuba* (14,039), *D. teissieri* (14,052), *D. erecta* (13,514)*, D. elegans* (13,741), *D. ficusphila* (13,199) and *D. kikkawai* (14,976)

We extracted AlphaFold 2 protein structure predictions of *D. melanogaster* from “UP000000803_7227_DROME_v4.tar” file downloaded on April 14, 2025 (https://ftp.ebi.ac.uk/pub/databases/alphafold/latest/UP000000803_7227_DROME_v4.tar), and relied on full sequence identity to successfully map these protein structures to the protein sequences obtained from FlyBase. Each gene from FlyBase was considered successfully mapped if its representative protein sequence was identical to one or more protein sequences from the PDB files from AlphaFold Protein Structure database. Of the 13,937 CDS from FlyBase, 12,899 and 23 sequences were successfully mapped to single and multiple PDB files, respectively, and 1,015 genes were not successfully mapped. We selected one PDB file randomly, if a given gene was mapped to multiple PDB files. Successfully mapped genes were inputs for AlphaFold-based approaches for intrinsic disorder prediction.

### Intrinsic disorder prediction

Numerous programs have been developed for intrinsic disorder prediction, and we leverage such programs to develop a strategy to classify residues into disordered and non-disordered. Inconsistency of disorder predictions among programs may create uncertainty in genome-wide patterns. Incorporation of evolutionary information can improve predictions [20] but renders downstream evolutionary analysis of disordered regions relatively challenging to interpret and might render spurious predictions. Hence, we have employed disorder prediction approaches that incorporate evolutionary information and that do not incorporate such information, and require sufficient support from both sets of approaches to annotate a given residue as disordered.

We considered programs that are widely-used and have been assessed as reliable previously [15,35]. flDPnn and AlphaFold-based approaches incorporate evolutionary information. We downloaded flDPnn program on April 14, 2025 from https://gitlab.com/sina.ghadermarzi/fldpnn. We ran flDpnn for 13,937 input protein sequences and 13,917 were processed successfully (i.e., outputs contain both disorder and disorder function predictions). The remaining 20 cases were not processed because their residue counts were 5,000 or above, which exceeds max count for flDPnn. Among the successful processes, we filtered low quality predictions as indicated by flDPnn. This filtering process retained outputs for 10,629 protein sequences. We employed AlphaFold-based approaches that rely on the only confidence scores of structure prediction (i.e., predicted local distance difference test [pLDDT])– AlphaFold-pLDDT –or both these scores and scores for relative solvent accessibility (AlphaFold-RSA). AlphaFold-pLDDT approach was implemented as described in equation 1 in Wilson et al., [75]. AlphaFold-RSA program was downloaded on April 14, 2025 (https://github.com/BioComputingUP/AlphaFold-disorder) and ran with default settings. We obtained intrinsic disorder predictions from AlphaFold-pLDDT and AlphaFold-RSA for the input 12,922 protein sequences that were successfully linked to PDB files.

We employed IUPred3 as an approach that does not incorporate evolutionary information. We downloaded IUPred3 program on April 14, 2025 from https://iupred3.elte.hu/download_new, and ran the program with optimizations for predicting short (IUPred-short) and long (IUPred-long) disorder. All other parameter settings were defaults. For both optimizations, 13,930 sequences were successfully processed, and the remaining seven cases were not processed because they contained below 20 residues which is less than the min count required by IUPred3 (when other parameters are defaults). Among the successfully processed cases, we excluded residues unexpected values (below zero or above one).

We developed a consensus strategy for disorder prediction and analyzed experimentally validated data from DisProt database to decide cut-offs for prediction approaches. The strategy assigns residues as disordered if the disorder score is equal to or above cut-offs for both the approaches that incorporate evolutionary information (i.e., any of flDPnn, AlphaFold-RSA, AlphaFold-pLDDT) and that do not incorporate such information (i.e., any of IUPred3-short or IUPred3-long). The remaining residues are classified as non-disordered. We exclude protein sequences that do not have sufficient data to make a prediction (i.e., disorder scores are not available for one approach or both). We leveraged the experimental annotations of DisProt to analyze the fraction of true positives of the consensus criteria under various sets of disorder score cut-offs (S1 Fig, S2 Table and S9 Text). We present results for a “non-strict” criteria set with the following cut-offs: 0.15 (flDPnn), 0.40 (AlphaFold-RSA), 0.25 (AlphaFold-pLDDT), 0.25 (IUPred3-short), and 0.25 (IUPred3-long). The resulting predictions achieve a true positive fraction of 0.944, which is higher (0.897) than when using commonly-used/default cut-offs such as 0.50.

We obtained residue-level disorder predictions under the consensus criteria for 13,910 of 13,937 protein sequences in *D. melanogaster*. We note that the predicted disordered residues analyzed in this study can belong to a disordered region of any length.

### Solvent accessibility

We obtained residue-level solvent accessibility values from DSSP version 3.0 (downloaded from https://anaconda.org/salilab/dssp on April 21, 2025). Inputs were 12,922 PDB files, and the resulting values were divided by theoretical max solvent accessibility values from Tien et al., [76] to obtain RSA. We classify residues as solvent-exposed and solvent-buried if RSA≥0.5 and RSA<0.5, respectively.

### Protein connectivity

We obtained protein-protein interaction counts from the STRING database [77]. We downloaded “7227.protein.links.full.v12.0.txt” from https://string-db.org/cgi/download on April 15, 2025. We retained only protein-protein interactions that have support from the “experimental” evidence type column (i.e., score above zero). We mapped STRING entries to those of FlyBase by comparing FBpp IDs. A given FBpp ID from FlyBase considered successfully mapped if there was an identical ID in STRING. Otherwise, it was considered unmapped. We successfully mapped 9,408 protein sequences and used the median count of protein interactors to classify proteins as hub (four interactors or more) and non-hub (less than four interactors).

### Disordered residue function analysis

We used outputs from flDPnn to assign putative functions for the residues that had been predicted to be disordered under the consensus criteria. A given residue was required to have intrinsic disorder support from both flDPnn and a consensus criteria set. Residues which passed this requirement were classified as putatively functional if there was function support from flDpnn and classified as non-functional otherwise.

### Molecular evolution analysis

We analyzed protein evolutionary rates at predicted disordered and non-disordered residues to relate intrinsic disorder and genome evolution. We obtained a set of candidate orthologs among members of the *Drosophila melanogaster* subgroup, and used this set to create CDS alignments. These alignments were filtered to exclude gaps before using PAML (Phylogenetics Analysis with Maximum likelihood) package to obtain estimates for replacement and silent substitution rates.

### Candidate ortholog set identification

We identified candidate ortholog sets among members of the *Drosophila melanogaster* subgroup using OrthoFinder version 2.5.5 downloaded from https://anaconda.org/bioconda/orthofinder/2.5.5/download/noarch/orthofinder-2.5.5-hdfd78af_2.tar.bz2 on May 30, 2025 [78]. We ran OrthoFinder using input protein sequences from the following nine *Drosophila* species: *D. melanogaster, D. simulans*, *D. mauritania*, *D. yakuba*, *D. teissieri*, *D. erecta*, *D. elegans*, *D. ficusphila*, *D. kikkawai*. Among the resulting candidate ortholog sets, we focused on sets containing a single gene from six relatively closely related members from the *Drosophila melanogaster* subgroup: *D. melanogaster, D. simulans*, *D. mauritania*, *D. yakuba*, *D. teissieri*, *D. erecta*. We extracted sets of candidate orthologs from the “N0.tsv” file which specifies candidate ortholog sets and each set has one gene per species. We retained candidate ortholog sets having one gene from each member of the *Drosophila melanogaster* subgroup. The resulting data contained 11,528 candidate orthologs sets. 9,696 and 1,831 ortholog sets belonged to genes found on autosomal and X-linked loci, respectively. Total residue counts were 5,110,229 and 1,069,670 for autosomal and X-linked loci, respectively.

### Multiple sequence alignments

We created CDS alignments for the candidate orthologs using MAFFT version 7.525 downloaded from https://anaconda.org/bioconda/mafft/7.525/download/osx-64/mafft-7.525-h2413b67_1.tar.bz2 on May 30, 2025 [72]. For each candidate ortholog set, we ran MAFFT with the E-INS-i algorithm setting. The resulting aligned protein sequences were back-translated to codons to create CDS alignments. Filtered CDS alignments were created by eliminating aligned columns that have gaps.

### Evolutionary rate estimation

We used codeml, implemented in PAML v4.9 [79], to obtain estimates of the replacement (*d_N_*), silent (*d_S_*) substitution rates and their ratio (*d_N_/d_S_*) along the branch of *D. melanogaster*. These estimates were obtained under a simplified implementation of the Yang and Nielsen [80] codon substitution model. This implementation allows separate parameters for transition-transversion rate ratio (κ) and equilibrium codon frequencies. We set κ to be fixed at two based on experimentally determined mutation rates [81,82] *Drosophila* species. Defaults were used for all other parameter settings. codeml was run for input filtered codon alignments and using an unrooted species tree. We ran codeml separately for autosomal and X-linked loci data.

### Randomized protein sequence analysis

Employing randomized protein sequences can allow investigating whether the prevalence and locations of intrinsic disorder can arise from biases in disorder prediction. Such biases are plausible if there are similarities between patterns on observed and randomized data. Differing patterns would imply that biases in disorder prediction are an unlikely explanation. This analysis employs flDPnn for disorder prediction because predicted disordered residues are more prevalent near terminals than internal segments, regardless of consensus criteria set or individual disorder prediction approaches employed in this study. In addition, previous studies support flDPnn as a relatively more reliable approach than the remaining ones employed in this study and has a shorter processing time than AlphaFold-based approaches [15,35].

We created randomized sequences by shuffling residue positions for observed protein sequences. This process was repeated 1,000 for a given observed sequence and is designed primarily to randomize relationships between amino acid composition and location. The resulting “Shuffled” sequences were used as input for flDPnn program. We analyzed a shuffled sequence only if its counterpart (i.e., Actual sequence) was successfully processed by flDPnn. We analyzed data for 9,018 protein sequence pairings and tested for differences in genome-wide patterns using LOR statistics and non-parametric approaches.

### Binning and statistical analysis

We focus on genome-wide patterns rather than gene/protein-specific ones, and thus employ data binning approaches to reduce statistical noise and simplify both data visualization and interpretation. Most analyses involve large sample sizes, and we employ both parametric and non-parametric approaches to evaluate the robustness of the inferences.

### Protein length bins

Protein length is a factor in the length (and number) of disordered regions, solvent accessibility, and evolutionary rates (e.g., [27,50]). We therefore created protein length bins after analyzing the distribution of protein sequence lengths and using the mean, 523 residues, to assign length categories (S3 Fig and S5 Table). We present results for bins with length ranges [19,100], [401,550], and [851,4976] and refer to these bins as “short”, “medium”, and “long” based on whether the range overlaps with the mean. Total gene/protein sequence counts were 3361, 2315 and 2146 for the short, medium and long protein length bins, respectively. Note that these bins contain protein sequences from autosomal and X-linked loci. For molecular evolution analysis, We created subsets of data for each protein length bin. Autosomal and X-linked loci were analyzed separately given their known distinct evolutionary patterns (e.g., [66]).

### Segment bins

We split protein sequences into segments that correspond to terminal and internal regions to be able to investigate the genome-wide distribution. Unless indicated otherwise, protein sequences were split into segments containing roughly 40 residues per segment following Lobanov et al., [36]. The number of segments per length bin was adjusted to allow for similar residue counts in all segments across protein length bins. The resulting first and last segments are assigned as “N-terminal” and “C-terminal” segments, respectively, while the remaining segments are assigned as “internal”. Results for internal segments are highly comparable therefore we present results (and statistical analysis) pooling data among all internal segments. We provide relatively detailed visualizations in the *Supplementary Materials*.

### Confidence interval estimation

We obtained 95% confidence intervals by sampling the representative protein sequence of each gene with replacement to create 100 to 1000 replicates depending on analysis.

### Odds ratios

We employed 2×2 contingency tables to calculate odds ratios for the following two types of comparisons: 1. fraction of predicted disordered residues between pairs of protein segments (N-terminal, Internal, C-terminal) and/or between classes of residues (e.g., exposed versus buried). 2. frequency of amino acids at predicted disordered/non-disordered residues between pairs of protein segments. We used odds ratios to calculate LORs, Z-statistics, and *p*-values, and tested for LOR differences using Wald test [83,84], calculating the Z-statistic as the difference between LORs values divided by the square root of the sum of squared standard errors.

### sAA_freq_diff

We used *sAA_freq_diff* to compare amino acid frequencies at disordered residues among segments, while accounting for frequencies at non-disordered residues.

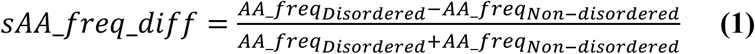

*sAA_freq_diff* ranges from −1 to 1 and sAA_freq_diff>0 and sAA_freq_diff<0 indicate higher and lower frequencies for disordered residues, respectively, and sAA_freq_diff=0 indicates identical frequencies.

### Correction for multiple tests

We used the Bonferroni sequential method [85], as implemented in the Python SciPy package [86].

