## Supplementary Material for "Location dependence of protein intrinsic disorder in *Drosophila melanogaster*"

### Supplementary Results and Discussion

#### *1. Development and implementation of intrinsic disorder prediction criteria*

Disorder prediction programs typically return residue-level scores. We use such scores from various programs to develop a consensus strategy for assigning disordered residues, accounting for some uncertainties of prediction. First, spurious predictions and ambiguous results may arise as a consequence of incorporating evolutionary information for disorder prediction, even though incorporating such information has improved prediction accuracy [1]. Second, disorder predictions can be conflicting among programs [2], partly due to program optimizations, implicitly or explicitly, for particular types of intrinsic disorder [3,4]. We employ multiple disorder prediction approaches that have been assessed as relatively reliable, and have been implemented as programs that are widely-used (discussed in *Materials and Methods*). flDPnn, AlphaFold-RSA or AlphaFold-pLDDT approaches incorporate evolutionary information, and the latter two have been assessed previously as relatively reliable predictors of conditionally-folded disordered regions. IUPred3-short and IUPred3-long approaches do not incorporate evolutionary information and are optimized for predicting short and long intrinsic disorder, respectively. We require sufficient support both from approaches that incorporate evolutionary information and that do not incorporate evolutionary information (S1A Fig). To determine disorder score cut-offs, we analyze disorder scores for experimentally-determined data obtained from DisProt database (S9 Text), and create instantiations of the consensus criteria that are permissive or stringent, relative to an instantiation in which commonly-used/default disorder scores (e.g., 0.5). Below, we describe disorder score distributions, consensus criteria development, and application to the *D. melanogaster* genome.

***Broad distribution of disorder scores for experimentally-determined data.*** We ran disorder prediction programs for input protein sequences (n=62, S1 and S2 Tables), including those known to have intrinsic disorder. Disorder scores are non-uniformly distributed (S1B Fig), and for most prediction approaches, scores of 10% or more residues have a score below 0.5. The distribution of disorder scores for flDPnn is roughly flat but peaks at lower (~0.1) and higher (~0.7) disorder scores, while distributions for AlphaFold-based and IUPred3 approaches are left-skewed and roughly bell-shaped/flat, respectively. These distributions indicate that commonly-used or default cut-offs for disorder scores can result in a high proportion of false negatives and that the specific proportion likely depends on the prediction approach.

**Differing recall of consensus criteria sets.** We determine cut-offs for disorder scores by comparing instantiations of the consensus criteria strategy for the prediction accuracy (i.e., in correct predictions among DisProt annotations). We refer to the main instantiation used throughout as "non-strict" as it is designed to allow for a higher fraction of true positives than a "comparison" criteria set in which default/commonly used disorder scores are used (0.314 for flDPnn and 0.5 for all other approaches). This non-strict consensus criteria set uses the following cut-offs: 0.15 (flDPnn), 0.40 (AlphaFold-RSA), 0.25 (AlphaFold-pLDDT), 0.25 (IUPred3-short), and 0.25 (IUPred3-long). These cut-offs lead to fractions of true positives of between 0.77 and 0.91, which are higher than values under "comparison" cut-offs (0.50–0.85 range) for each disorder prediction approach. The non-strict consensus criteria set may allow, in principle, for more false positives, and we create "strict" consensus criteria with the following cut-offs: 0.50 (flDPnn), 0.80 (AlphaFold-RSA), 0.60 (AlphaFold-pLDDT), 0.60 (IUPred3-short), and 0.70 (IUPred3-long). The resulting fractions of true positives is between 0.53 and 0.60, which, as expected, is lower than the values under the cut-offs for the comparison criteria. Overall, we observe that the fraction of true positives is 0.944 under the non-strict consensus criteria set, which is 1.4 times higher than the fraction under the strict set, 0.673 (S1C Fig and S3 Table). These fractions differ from the fraction for the comparison criteria (0.897).

**Sensitivity of genome-wide disorder statistics to consensus criteria sets.** We employed the non-strict and strict consensus criteria sets to assign disordered residues in the *D. melanogaster* genome. The fraction of predicted disordered residues is 0.414 and 0.117 under the non-strict and strict consensus criteria sets, respectively (S2A Fig and S3 Table), and length distributions of predicted disordered segments for both sets are right-skewed (S2B Fig and S4 Table); shorter segments are more common than longer ones. We focus on the non-strict consensus criteria set due to its higher recall rate on the DisProt dataset.

**Uneven distribution of disordered residues.** We analyze the distribution of predicted disordered residues along protein sequences, and use "within-terminal" comparisons to analyze the fraction of predicted disordered residues among the first or last 40 residue positions, and use "among-segment" comparisons to analyze that fraction along the full length of protein sequences among various protein length bins (discussed in Materials and Methods. S5 Table). Both comparisons indicate that intrinsic disorder is more prevalence at protein terminals than internal parts; the fraction of predicted disordered residues associates with the number of residue positions relative to the terminal residue (S6 Table), for N-terminal ($r_s$=-0.169) and C-terminal ($r_s$=-0.999). Patterns for the among-segment comparisons are consistent with those in Fig 1. Internal segments appear to have similar fractions of disordered residues, but a notable exception is the medium protein length bin where fractions are higher for internal segments closer to protein ends than middle parts, and we note that this apparent location dependence is generally robust to protein length (S3 Fig and S7 Table). Overall, the apparent location dependence of intrinsic disorder is associated with higher fractions of predicted disordered residues at the first and last 10-20 residue positions, and observed under the non-strict and strict consensus criteria sets (data not shown). Previous studies based on within-terminal comparison have shown support for prevalence of intrinsic disorder at terminals, especially at the first or last ten positions [5,6]. This is consistent with our observations, but the among-segment comparisons extend the scope of observations.

##### *2. Biophysical and functional factors in intrinsic disorder presence*

We found that the apparent location dependence of intrinsic disorder is associated with amino acid compositional variation that reflects proteome-wide patterns and/or possible functional differences among disordered regions (Fig 2, S5–8 Figs, S8 and 9 Tables). We investigate the location dependence by analyzing relationships between protein tertiary structure and intrinsic disorder. We first analyze relative solvent accessibility values (RSA) and disorder scores along protein sequences, and then analyze the distribution of predicted disordered residues in putative solvent-exposed, solvent-buried and functional (e.g., protein-binding, RNA-binding) contexts.

**High RSA at protein terminals.** Within-terminal and among-segment comparisons of mean RSA support higher exposure of protein terminals than internal parts. Terminal segments have higher mean RSA (>~0.5) than internal segments, with highest values observed for the N-terminal segment (S9 Fig). These patterns are associated with substantial increases in RSA values at the first and last five residues. Mean RSA values and disorder scores for most approaches exhibit similar trends along protein sequences. An exception to this pattern is the disorder scores IUPred3-long, where association with mean RSA values appears weaker

(S10–S12 Figs). These results suggest that disordered regions near terminal residues tend to be short, and that the relationship between RSA values and disorder scores is unlikely fully explained by use of RSA for disorder prediction; RSA values are associated with disorder scores of flDPnn, AlphaFold-pLDDT, and IUPred3-short, which do not employ RSA for disorder prediction.

**RSA-associated changes in disorder prevalence along proteins.** We analyzed the distribution of predicted disordered residues in solvent-exposed and solvent-buried contexts (as described in Materials and Methods). Both within-terminal and among-segment comparisons show support for location dependence of intrinsic disorder, revealing substantial differences in solvent-exposed and solvent-buried contexts. The fraction of predicted disordered residues is strongly associated with the distance relative to the terminal residue, for the solvent-exposed class ($r_s$= -0.320 and -0.996 at $p$<0.05, for the N and C terminals, respectively. S10 Table and S13 Fig). Such an association is in the opposite direction or weaker for the solvent-buried class ($r_s$= 0.508 and -0.526 at $p$<0.05 for the N and C terminals, respectively). Interestingly, the fraction of predicted disordered residues summits at the terminal residue for the solvent-exposed class, but plummets at that residue for the solvent-buried class (S13A Fig).

The fraction of predicted disordered residues is higher for the solvent-exposed class than the solvent-buried one (Fig 3, S13 Fig and S11 Table). Among-segment comparisons reveal that the fraction is significantly higher for terminal segments than internal ones for the solvent-exposed class, but not the solvent-buried class (S13B Fig and S12 Table). These results are consistent with the notion that solvent accessibility is a general feature of intrinsic disorder, and suggest that it may shape the locations of disorder along protein sequences.

**Functional class differences for total disorder prevalence.** We found support that, in a solvent-exposed context, protein-binding is a common function of intrinsic disorder, relative to RNA-binding, DNA-binding and linker functions (Fig 4, S14 Fig and S13 Table). The protein-binding class shows the highest fraction of disordered residues compared to the remaining classes, a pattern observed for each protein segment (S14 Fig and S13 Table). For example, in the short protein length bin, LORs are 0.58, 0.79, and 0.64 for comparisons between protein-binding versus DNA-binding classes for N-terminal, internal and C-terminal segments, respectively ($p$<0.05 for these LOR differences. S14 Fig and S13 Table). The DNA-binding class has a significantly higher fraction of predicted disordered residues than both the linker and RNA-binding classes, with the latter generally having the lowest fraction among all classes, in particular, at the internal segment (note that LORs<0 in S13 Table). These patterns are generally observed across protein segments and length bins and suggest intermolecular protein-binding may be a possible primary function of intrinsic disorder.

***Apparent functional class-specific changes of intrinsic disorder prevalence along proteins.*** Our previous analysis hints at a functional specificity of intrinsic disorder that depends on location, in particular for RNA-binding and DNA-binding functions, but whether these patterns reflect individual effects of the putative functions remain unclear. We address this issue by analyzing the subset of predicted disordered that have been assigned only one function by flDPnn, and the following classes were employed: “protein-only binding”, “RNA-only binding”, “DNA-only binding” and “linker-only”. We find general support for individual effects; RNA-only binding and DNA-only binding exhibit steeper patterns than protein-only binding and linker-only (S15 Fig, S13 and S14 Tables). For example, in the long protein length bin, LOR comparisons for N-terminal versus internal segments reveals a higher value for RNA-only binding class (1.88) than protein-only binding (0.53), DNA-only binding (0.34), linker-only (0.41) classes ($p<0.05$ for Wald tests of LORs for RNA-only binding class versus all other classes). We observe a similar pattern for the short protein length bin, but not the medium protein length bin (LOR for RNA-only binding class, 1.37, is second only to that for the DNA-only binding class, 1.39, but $p>0.05$ for a Wald test). These results suggest that, compared to protein-binding and linker functions of disordered regions, nucleic acid-binding functions are disproportionately more likely to occur at protein terminals than at internal regions.

We find that protein-only and DNA-only/RNA-only binding classes tend to be the most and least prevalent “single-function” classes, respectively. LORs are above zero for comparisons of the fraction of predicted disorder residues between protein-only binding and each of the remaining single-function classes. LORs are below zero for comparisons between the RNA-only binding class and each of the remaining classes. The LORs can be above zero or below for other comparisons. Within-terminal analysis shows that the fraction of predicted disordered residues tends to increase towards the terminal residues, with the steepest and flattest patterns observed for the protein-only binding class and linker-only class respectively (data not shown).

Compared to the putative solvent-exposed class, patterns for the solvent-buried class are generally flatter (S16 Fig), but show support for location dependence.

##### *3. Protein connectivity and intrinsic disorder at terminals*

We employed within-terminal comparisons to investigate the role of intrinsic disorder in protein connectivity. The fraction of predicted disordered residues is significantly higher for the hub than non-hub proteins (LOR=0.380 and 0.230 at $p<0.001$ for hub versus non-hub comparisons at N- and C-terminals, respectively. S15 Table). These observations, along with observations from among-segment comparisons (Fig 5, S17B Fig, S16 and S17 Tables), are consistent with the view that flexibility facilitates binding plasticity, but show that protein length and the location of a disordered region are factors.

##### *4. Functional enrichment and intrinsic disorder prevalence*

We further tested relationships between intrinsic disorder locations and function by performing GO (gene ontology) terms analysis, leveraging experimentally verified annotations from FlyBase (S10 Text). Multiple GO terms are enriched at one segment or more. For example, “male germ cell nucleus” is enriched at terminal segments but not internal segments, and “haltere development” is enriched only at the latter segment. GO terms “negative regulation of autophagy” and “neuropeptide receptor activity”, are enriched at internal and C-terminal segments, respectively. The GO term “RCAF complex” is a notably strong case of enrichment at the N-terminal. This complex consists of multiple histone proteins and a high fraction of residues are predicted to be disordered (96.4%; S19 Table). Since this complex enables nucleosome assembly on newly replicated DNA and constitutes histone 3, histone 4 and the anti-silencing factor 1 [7], intrinsic disorder at N-terminals may serve roles in nucleosome assembly, which would be consistent with previous studies in human [8,9]. These results serve as further support for location-specific functions of intrinsic disorder, and we note that some of these functions may involve long disordered regions that span multiple protein sequences.

##### *5. Accounting for evolutionary information in intrinsic disorder predictions*

We showed support for faster evolution and weaker constraints at disordered residues than non-disordered residues (Figs 6B, 6C and S18 Fig), with exception pattern N-terminal where such differences are observed for the long protein length bin but not the short and medium protein length bins. Such a pattern may arise from the fact that flDPnn, AlphaFold-RSA and AlphaFold-pLDDT incorporate evolutionary information, implicitly or explicitly, for intrinsic disorder prediction. We attempt to address this issue by performing evolutionary analysis based on predictions from IUPred3-short. We observe the same general patterns as in the initial analysis: higher evolutionary rates at predicted disordered than non-disordered residues and no readily discernable $d_N$ and $d_N/d_S$ differences among predicted disordered residues (S19 Fig), with exception to the long protein length bin. Interestingly, for the N-terminal segment, $d_N$ and $d_N/d_S$ values are similar between predicted disordered and non-disordered residues, across all protein length bins and autosomal and X-linked loci, which suggests an evolutionary process independent of intrinsic disorder presence.

##### 6. *Inability of disorder prediction biases to fully explain regional trends*

We found that disorder prediction biases can be a relevant factor in the uneven distribution of disordered residues and amino acid frequencies (Figs 7, 8. S19–25 Tables). We further investigated such biases using non-parametric approaches. To quantify the extent of unevenness in the distribution of disordered residues, we calculate the "magnitude of disorder distribution unevenness", MDDU, for segment pairings (e.g., N-terminal and internal). MDDU values above and below zero respectively, indicate greater and lesser unevenness in the Actual data (S11 Text). We found that MDDU values are non-zero in all comparisons. For example, MDDU for N-terminal and internal segments is substantially above zero (i.e., error bars do not overlap with zero) in the short and medium protein length bins, but marginally below zero in the long protein length bin (S20 Fig). MDDU values for C-terminal versus internal segments are substantially above zero for the short protein length bin, but below zero for the long protein length bin and MDDU value for N-terminal and C-terminal segments is substantially lower than zero for the medium length protein length bin, but not the remaining bins. These results are generally consistent with comparisons based on the fraction of predicted disordered residues (Fig 7, SFig 20), and show that disorder prediction biases are a factor in the distribution of disordered residues, and that the magnitude and direction of these biases is associated with protein length.

The amino acid analyses demonstrated considerable effects of disorder prediction biases, but also revealed support for regional differences in amino acid frequencies that cannot be fully explained by the biases (Fig 8). We further investigated this by visual analysis of amino acid frequency distributions for Actual and Shuffled data. We found six instances –

Lys, Leu, Pro, Ser, Tyr and Val – where those datasets clearly differ with respect to frequency and/or frequency distribution, in one or more protein length bins (S21–23 Figs). These observations suggest that disorder prediction biases are unlikely to explain the regional heterogeneity in Lys frequencies in the Actual data, as well as patterns for Leu, Pro, Ser, Tyr, and Val. For other amino acids, support exists for a role of disorder prediction biases (i.e., LORs are non-significant in many cases), but the explanatory scope is associated with the protein length bin. We further tested for differences between the Actual and Shuffled data by calculating the magnitude of amino acid distribution unevenness, MADU (S11 Text). MADU is similar to MDDU, and quantifies the extent to which the Actual and Shuffled datasets exhibit the same unevenness in amino acid frequencies for a given segment pair, accounting for amino acid frequencies at disordered and non-disordered residues. We observe many instances – such as Ala, Cys, Gly, Ile, Lys, Met, and Val – where MADU is substantially above or below zero and error bars do not overlap zero, for one or more segment pairs, suggesting that amino acid frequency differences in the Shuffled data unlikely explain those differences in the Actual data (S24 Fig). Overall, these amino acid compositional analyses, similar to the LOR-based analysis, support a role of disorder prediction biases in regional heterogeneity, including for Met, but reveal that the biases cannot explain fully the heterogeneity. Length-associated effects are also apparent from these analyses.

##### *7.* Support for disorder functions in experimentally-determined data

We analyzed intrinsic disorder annotations from DisProt database to gauge whether the genome-wide patterns are biologically plausible. The annotations are scant, and only 62 protein sequences in *D. melanogaster* have annotations. We perform analysis for the entire dataset (All) as well as subset bins. The distribution of protein sequence lengths is right-skewed and we use the median, 592 residues, to delimit shorter (<592) and longer (>592) protein sequences (S25A Fig). The fraction of disordered residues appears to differ marginally among segments, across the three data categories, and generally, higher fractions of disordered residues are observed for terminal segments than internal ones (S25B Fig). LOR-based comparisons of these fractions support significant differences among segments. These patterns support an uneven distribution of disordered residues, but the lack of data appears to weaken statistical support.

We also analyzed amino acid frequencies at disordered residues and found that frequencies of multiple amino acids – such as Ile, Tyr, Pro, Asp and Val – differed substantially among segments in one or more data categories (S26 and S27 Figs). Interestingly, Met frequencies were consistently higher for the N-terminal segment than the internal segment, across all data categories. These differences were significant in “All” and “<592” data categories. These observations are consistent with biological plausibility of the regional heterogeneity of amino acid composition related to intrinsic disorder. Due to the lack of annotations, the challenge remains to distinguish disorder-associated effects from proteome-wide effects.

##### ***8. Consistency of genome-wide patterns among non-Drosophila species***

Our results in *D. melanogaster* support that most disordered regions exist in protein terminals and that solvent accessibility and functions are important factors in the distribution of disordered regions. Ubiquity of these observations across other taxa can hint at evolutionary conservation. We explored this possibility by employing AlphaFold-pLDDT predictions to analyze prevalence of intrinsic disorder in the following organisms used as models in many studies: human, mouse, zebrafish, *C. elegans*, *A. thaliana*, *S. cerevisiae* and *E. coli* (S12 Text). Terminal segments have higher fractions of predicted disordered residues than internal segments, in both solvent-exposed and solvent-buried contexts (S28 Fig). However patterns are steeper for the latter context, seemingly due to lower disorder prevalence at C-terminals.

We also investigated whether intrinsic disorder may serve the same main functions across taxa. We thus analyzed flDPnn predictions for human and *E. coli*, which we employ as representatives of complex and simpler organisms, respectively. For both organisms, the protein-binding class has the higher fraction of predicted disordered residues, and the fractions are comparable among the remaining functional classes (S29 Fig). The steepest and flattest patterns appear to be for protein-binding and RNA-binding classes respectively, in human and protein-binding and linker in *E. coli*. Future analysis will require testing for effects of disorder prediction biases in these patterns. Overall, these results are compatible with the possibility of conserved locations of disordered regions, and hint at lineage-specific relationships between intrinsic disorder and solvent accessibility. We note that the former possibility is supported in analysis of experimental data of several organisms (S29 Fig); the fraction of *disordered* residues tends to be higher for terminal segments than internal ones in *D. melanogaster* and the seven other organisms analyzed. The differences can be considerable in all organisms analyzed except in zebrafish and *C. elegans*, which have the least data available.

### Supplementary Methods

#### *9. Experimentally-determined intrinsic disorder analysis*

We analyzed disorder scores at experimentally-determined disordered residues to decide cut-offs for classifying residues as disordered or non-disordered and to check whether patterns based on genome-wide predictions of intrinsic disorder are also observed in experimental data. We obtained data from DisProt which is widely used and contains manually curated annotations for multiple organisms including *D. melanogaster*. We downloaded the database file "DisProt release_2024_12.tsv" (https://disprot.org/download, accessed on May 7, 2025), which tabulates various types of information including annotations for intrinsic disorder in protein segments for multiple organisms. Each row includes a UniProt ID and, in the cases of intrinsic disorder annotations, includes positions of disordered residues within the protein, but does not include the full protein sequence. We implemented a sequential process to link disorder annotations and disorder scores. First, we retain rows for an organism of interest (i.e., rows that specify *D. melanogaster* for the "Organism" column). Next, for a given protein sequence, we retain rows that specify annotations of intrinsic disorder (i.e., rows having "disorder" for the "term_name" column). Note that this step requires obtaining full length protein sequences. We obtained such sequences from UniProt database and relied on ID and sequence comparisons to link data from DisProt to UniProt. A given protein was considered successfully mapped if the ID from DisProt had an identical ID in UniProt database and if sequences for disordered segments from DisProt and UniProt are identical (note that DisProt provides residue position information for disorder annotations, and we use such information to extract disordered segments from UniProt data and compare those segments with corresponding segments in DisProt data).

We successfully mapped most or all available protein sequences for each organism (≥98% or above. S26 Table).

#### *10. Gene ontology terms analysis*

We aimed to test associations between locations and possible functions of intrinsic disorder by employing FlyBase gene ontology (GO) data. We downloaded the "gene_association.fb" file (https://flybase-ftp.s3.us-east-1.amazonaws.com/releases/FB2025_01/precomputed_files/index.html, last accessed on April

15, 2025). This file contains gene ontology terms 8,677 terms for 13,789 genes (protein- and non-protein coding genes), and has specific annotations for objects such as "gene" and "protein". We retained data for protein objects, and excluded annotations if the only evidence was based on computational prediction or curator statements. Of the remaining GO terms annotations, we excluded cases of indirect association. Cases with the following key words were excluded: "colocalizes", "acts_upstream_of", "acts_upstream_of_positive_effect", "acts_upstream_of_negative_effect". We obtained 5,693 terms across 6,575 genes.

We were interested in testing associations between GO term enrichments and short IDRs within a particular segment or long IDR that span terminal and internal segments. Therefore, we split protein sequences into four segments in proportion to length, and employed a hypergeometric approach to test for enrichment of GO terms in relation to intrinsic disorder prevalence, separately for each segment. For a given segment, we sorted units (i.e., gene IDs) for the fraction of predicted disordered residues. We then included a given unit, and thus its GO term, in a "test" sample if its fraction of predicted disordered residues belonged to the 90th percentile (calculated among units for the same segment class). Each test sample GO term was checked for enrichment, using the hypergeometric model implemented in "hypergeom" in the Python scipy package [10]. As inputs, we used the unit count for the GO term being tested as the sample and the total unit count among all GO terms as the reference set.

##### *11. Comparisons between Actual and Shuffled data*

We employed MDDU to compare Actual and Shuffled data for scaled differences in the fraction of predicted disordered residues among segment pairings such as N-terminal and internal segments. We first obtain the difference for a given segment pair separately for Actual and Shuffled data.

$$X_{AB} = \left|\frac{DR_A - DR_B}{DR_A + DR_B}\right| \qquad \textbf{(S1 Eq)}$$

where "X" specifies Actual or Shuffled data and AB specifies the segment pair being compared and "DR" specifies the fraction of predicted disordered residues. We then obtain the scaled difference between the two datasets:

$$\text{MDDU}_{AB} = \frac{Actual_{AB} - Shuffled_{AB}}{Actual_{AB} + Shuffled_{AB}} \qquad \textbf{(S2 Eq)}$$

MDDU ranges from -1 to 1 and MDDU>0 and MDDU<0 indicate greater and less unevenness, respectively in the Actual data than the Shuffled data, and MDDU=0 indicates similar unevenness. We note that MDDU informs only on magnitude of differences and not differences in distribution shapes (e.g., MDDU can be zero even though distributions differ between Actual and Shuffled data). In addition, the approach of MDDU is likely conservative since multiple difference statistics are used for calculation.

We aimed to check whether Actual and Shuffled data exhibit the same extent of difference for disordered amino acid frequencies between segment pairs and employed MADU statistic. MADU allows such checks while accounting for non-disordered amino acid frequencies. Calculation and interpretation of MADU is analogous to MDDU (explained in Materials and Methods), with the only difference being that absolute values for sAA_freq_diff are used to solve S2 Eq. MADU can take up values from -1.0 to 1.0 and MADU>0 and MADU<0 indicate higher and lower extents of difference, respectively in the Actual data than the Shuffled data while MADU=0.0 indicates identical extents of difference. MADU limitations are the same as MDDU (e.g., it does not account for the shape of the distribution, is a conservative analysis approach because it is calculated from other difference statistics, which accumulates variance).

##### *12. Disorder prediction analysis in non-Drosophila species*

We aimed to gauge whether the genome-wide patterns in *D. melanogaster* are widespread across taxa. We downloaded AlphaFold protein structure predictions (https://ftp.ebi.ac.uk/pub/databases/alphafold/latest, last accessed on June 1, 2025) for the following taxa: Human (UP000005640_9606_HUMAN_v4.tar), Mouse (UP000000589_10090_MOUSE_v4.tar), Zebrafish (UP000000437_7955_DANRE_v4.tar), *Caenorhabditis elegans* (UP000001940_6239_CAEEL_v4.tar), *Arabidopsis thaliana* (UP000006548_3702_ARATH_v4.tar), *Saccharomyces cerevisiae* (UP000002311_559292_YEAST_v4.tar), and *Escherichia coli* (UP000000625_83333_ECOLI_v4.tar).

The protein structure predictions (PDB files) were inputs for AlphaFold_pLDDT approach (S2 and S26 Tables). For each species, one or more PDB files contain protein structure predictions for the same protein sequence, and we randomly selected one file in such cases. For a given organism, we analyzed the non-redundant set of protein sequences. We split protein sequences as described in *Materials and Methods*.

### Supplementary Figures

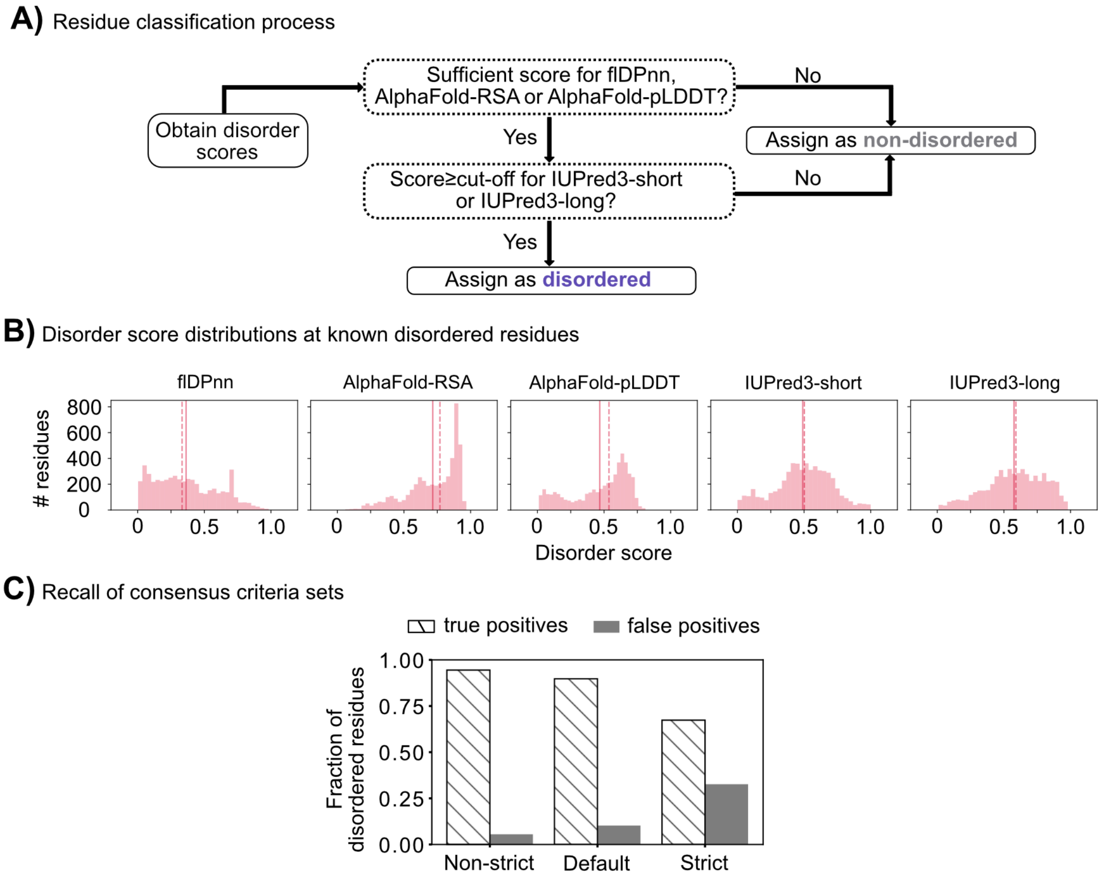

**S1 Fig. Developing consensus criteria for assigning disordered residues.**
**A)** A strategy was developed to assign disordered residues considering the predictions of existing approaches that rely on evolutionary information and that do not rely on such information. Residues having support for both approaches are referred to as "disordered". Because the programs are designed for disorder prediction, the remaining residues are referred to as "non-disordered" rather than "ordered" or "structured". Instantiations of the consensus criteria were created and differ *only* for the disorder score cut-offs of each disorder prediction approach. **B)** To determine cut-offs, predictions of each approach were obtained for *D. melanogaster* protein sequences (n=62) that have one or more disorder annotations in DisProt (discussed in S1 and S9 Text). The resulting distributions are shown and solid and dashed lines specify mean and median disorder scores, respectively. **C)** Results for residue assignments based on, "non-strict" and "strict" criteria sets are shown. These are defined relative to a set in which commonly-used/default cut-offs are used (see S1 Text). The criteria sets were evaluated by analyzing the fraction (residues) of accurate predictions. Further descriptive statistics are presented in S1 and S2 Tables.

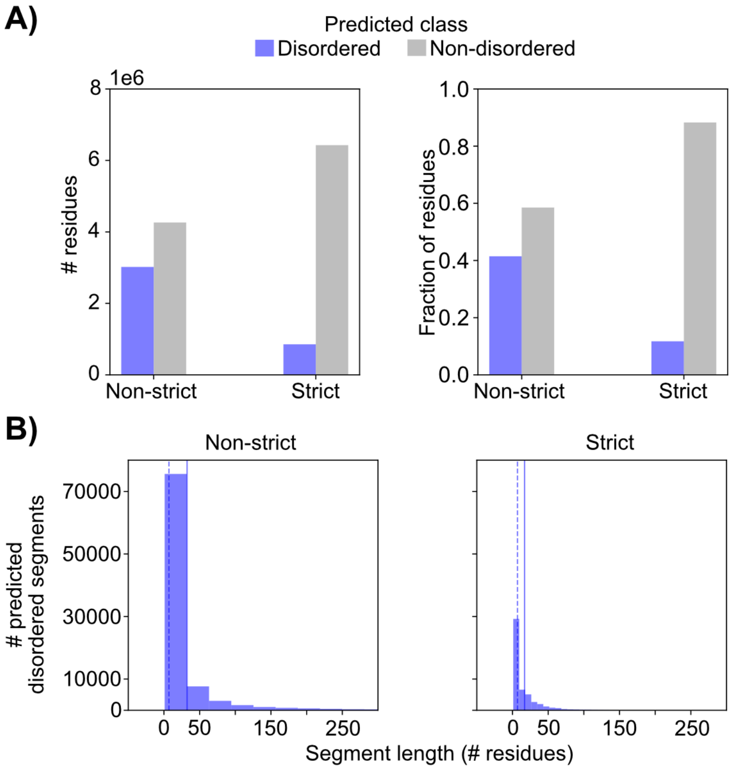

**S2 Fig. Prevalence and length of predicted intrinsic disorder.**
The non-strict and strict consensus criteria sets were employed for assigning intrinsic disorder in 13910 protein sequences (one per gene). **A)** Total counts and fractions of predicted disordered and non-disordered residues. **B)** Distribution of predicted disordered segment lengths. Solid and dashed lines specify mean and median lengths, respectively. Descriptive statistics are shown in S3 and S4 Tables.

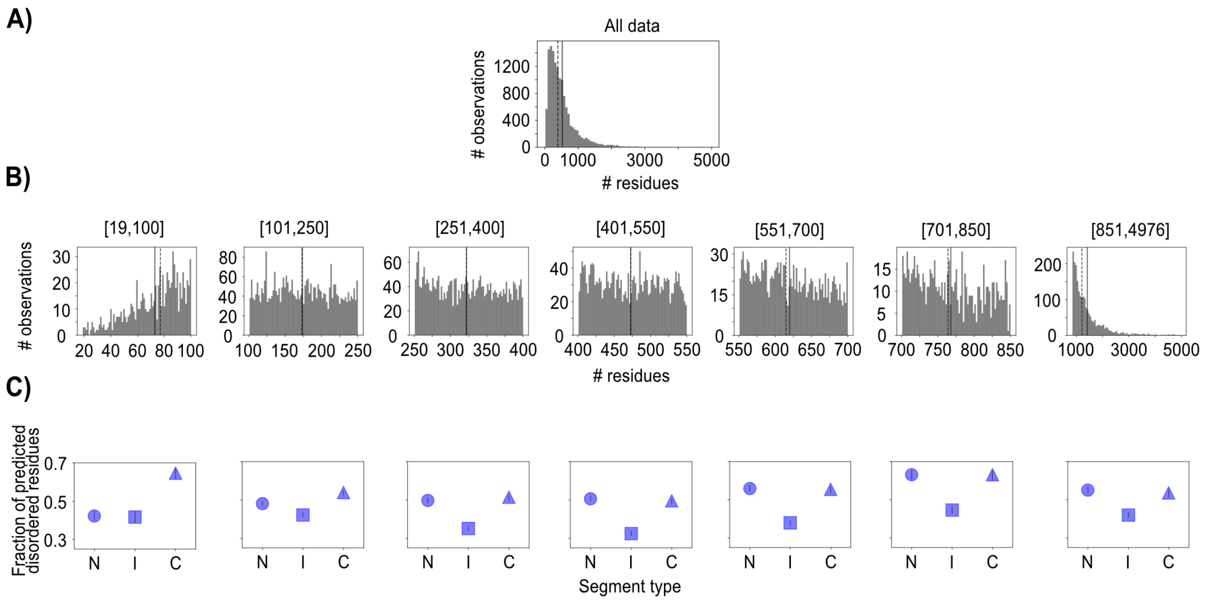

**S3 Fig. Protein length bin residue counts and fraction of predicted disordered residues.** Bins were created following analysis of the distribution of protein lengths in *D. melanogaster*. The fraction of predicted disordered residues was then analyzed for each bin, under the non-strict consensus criteria set. These fractions are shown separately for N-terminal (N), internal (I) and C-terminal (C) segments. **A)** Distribution of protein lengths. Only sequences used for assigning intrinsic disorder were included (n=13910). **B)** Distribution of protein sequence lengths for each bin. The residue count range is shown for each bin. Samples sizes and descriptive statistics are shown in S5 Table. **C)** The fraction of disordered residues along protein sequences. Protein sequences were split to contain roughly 40 residues for the terminal segments (i.e., first/last residues) and the remaining residues were used to create the internal segment, following the approach of Lobanov et al., (2010). The exception to this process is bin [19,100] for which terminal segments contain roughly 20 residues. Note that these patterns remain even when protein sequences are split in proportion to length. For downstream analyses, bins [101,250], [401,550], and [851,4976] were used as representatives for short, medium and long proteins. These bins have above 2000 genes and above 500,000 residues. Error bars are 95% confidence intervals from 500 bootstrap replicates created by sampling genes with replacement.

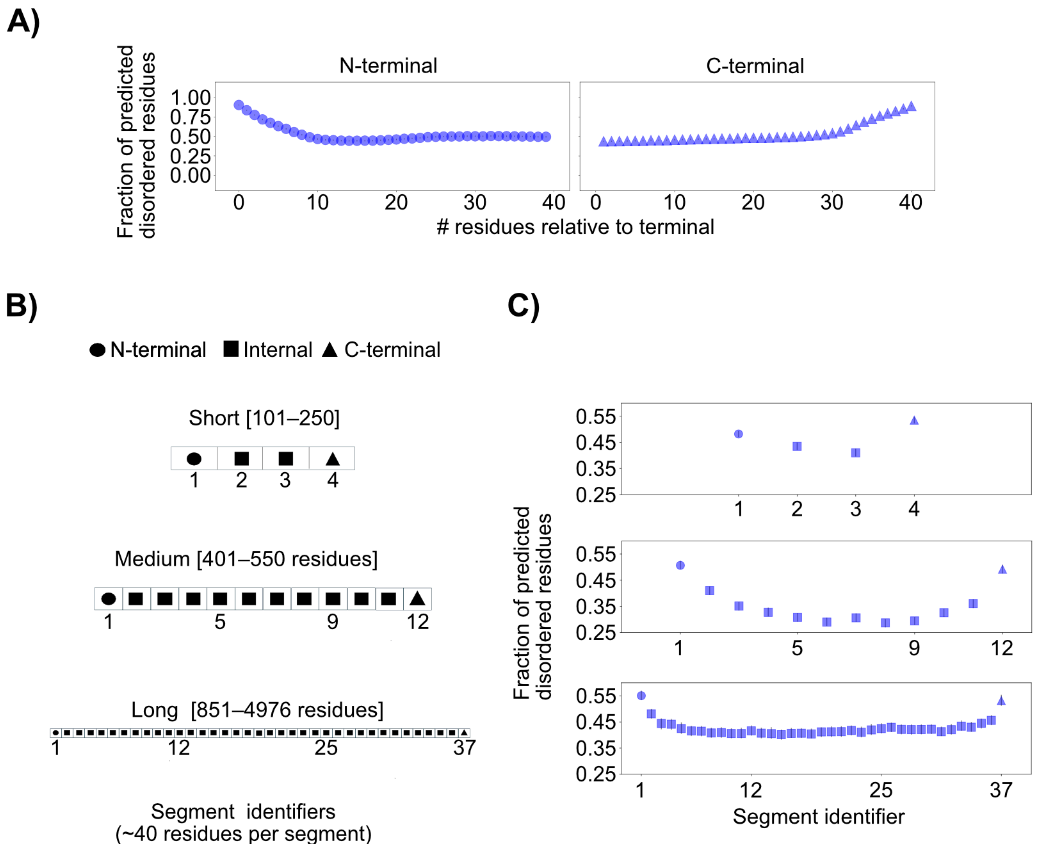

**S4 Fig. Fraction of predicted disordered residues along protein sequences.** Values under the non-strict consensus criteria set are shown. These values are calculated as the total counts of predicted disordered residues (among protein sequences) at a given segment divided by the total count of residues at that segment. **A)** Prevalence of intrinsic disorder for N- and C- terminal positions. Protein sequences with 80 residues or above are analyzed. **B)** Protein segments and length bins. Protein sequences were grouped for their residue counts and split into parts corresponding to terminal and internal segments, as described in *Materials and Methods*. **C)** Prevalence of intrinsic disorder for N-terminal, internal and C-terminal segments for each bin. All error bars are 95% confidence intervals calculated among 500 or 1000 replicates for the within-terminal and among-segment analyses, respectively. Replicates were created by sampling genes with replacement. n=3361, 2315 and 2146 genes for the short, medium and long protein length bins, respectively. Statistical test results are presented in S6 and S7 Tables.

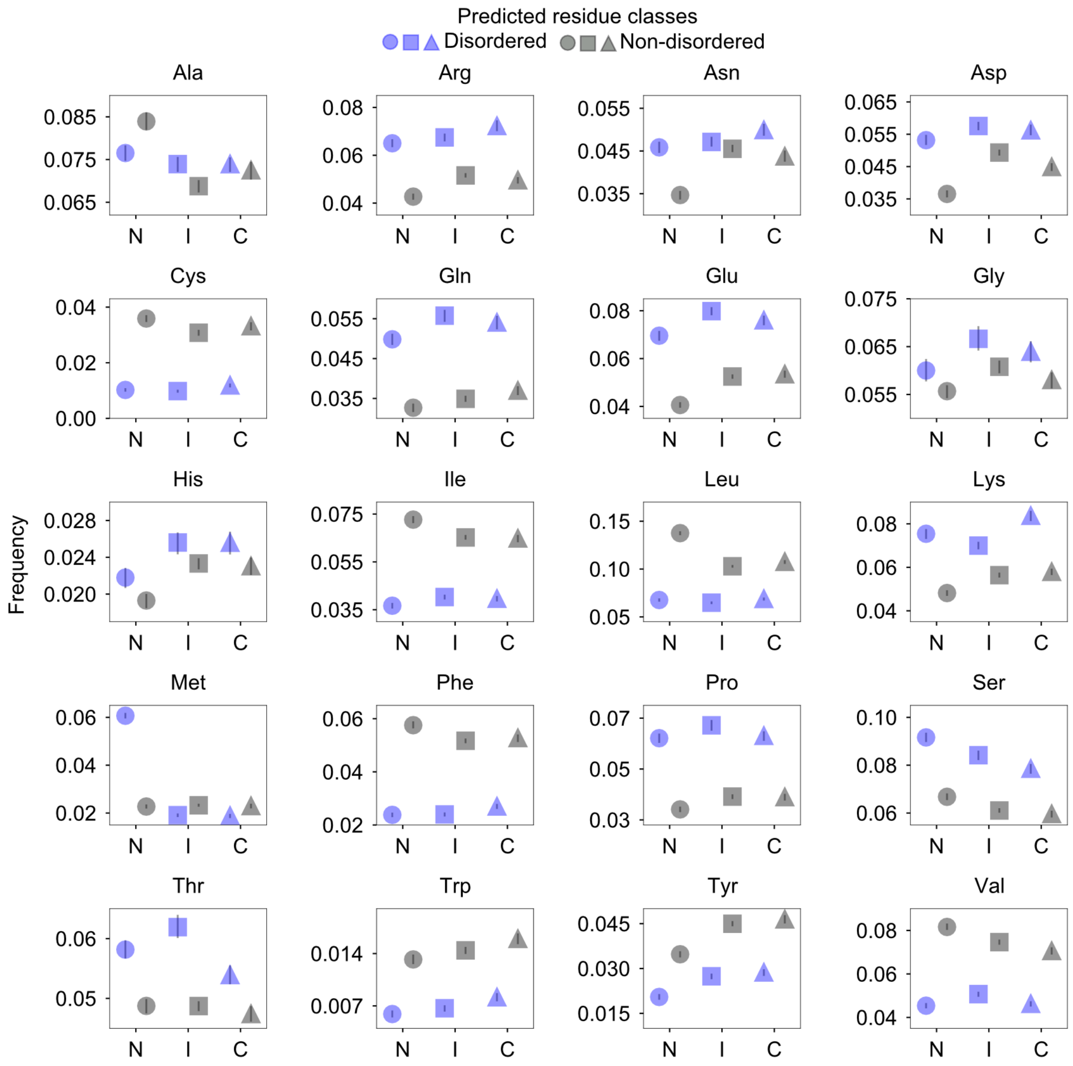

**S5 Fig. Amino acid frequencies at predicted disordered and non-disordered residues along short protein sequences.**

Frequencies at terminal (N, C) and internal (I) segments are shown. All error bars are 95% confidence intervals calculated among 1000 replicates. n=3361 genes were used. Replicates were created by sampling genes with replacement. Statistical test results are presented in S8 and S9 Tables.

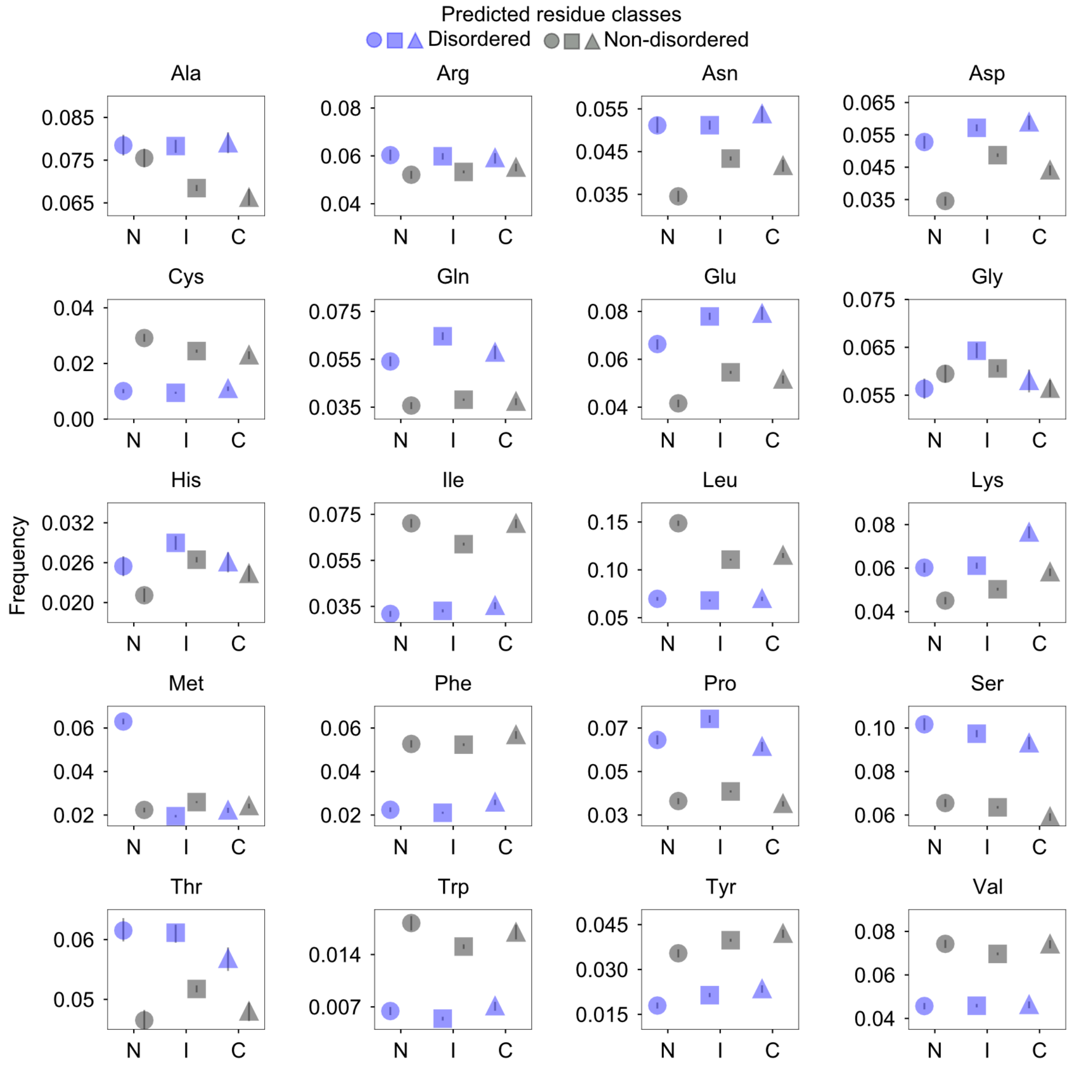

**S6 Fig. Amino acid frequencies at predicted disordered and non-disordered residues along medium-length protein sequences.**

Frequencies at terminal (N, C) and internal (I) segments are shown. All error bars are 95% confidence intervals calculated among 1000 replicates. n=2315 genes were used. Replicates were created by sampling genes with replacement. Statistical test results are presented in S8 and S9 Tables.

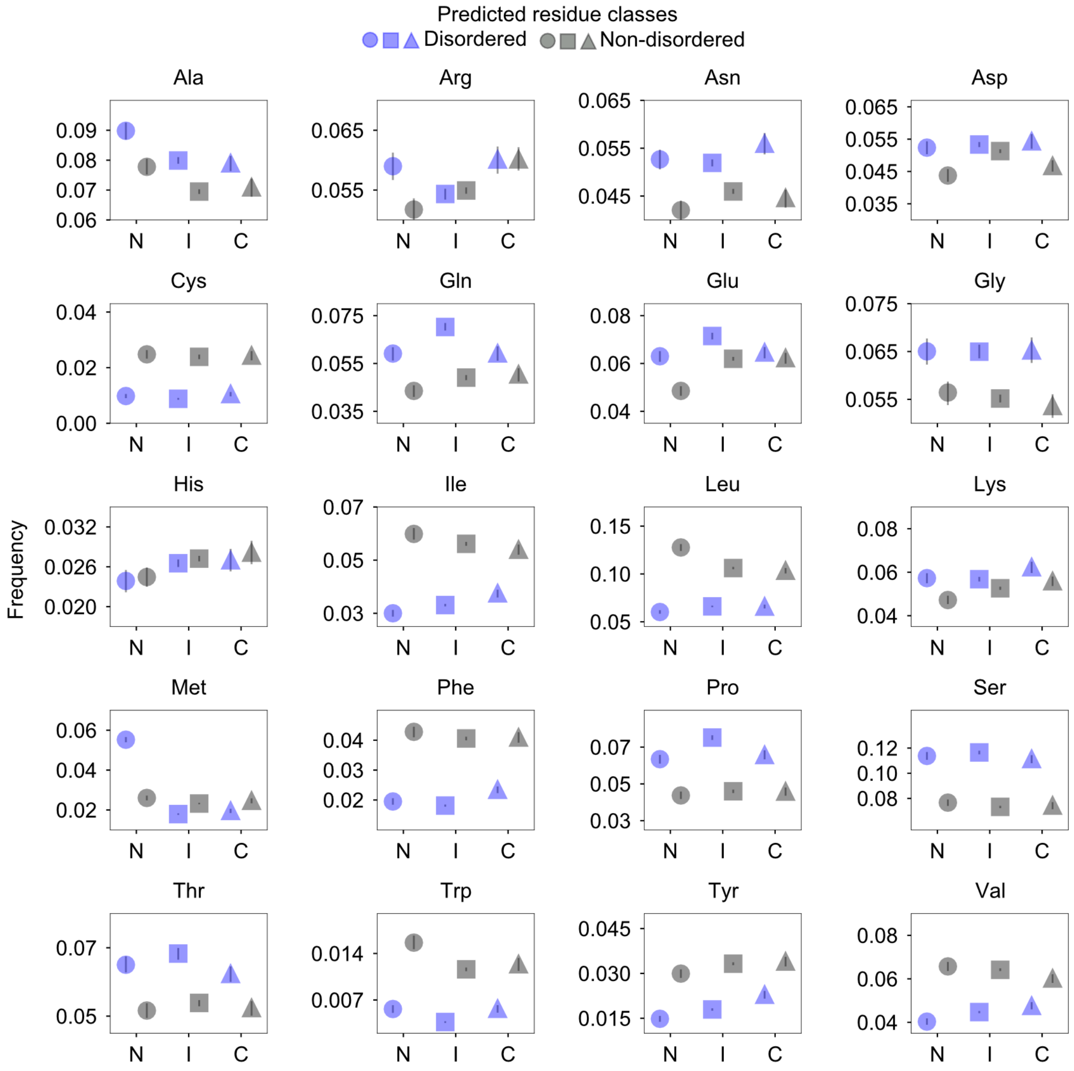

**S7 Fig. Amino acid frequencies at predicted disordered and non-disordered residues along long protein sequences.**

Frequencies at terminal (N, C) and internal (I) segments are shown. All error bars are 95% confidence intervals calculated among 1000 replicates. n=2146 genes were used. Replicates were created by sampling genes with replacement. Statistical test results are presented in S8 and S9 Tables.

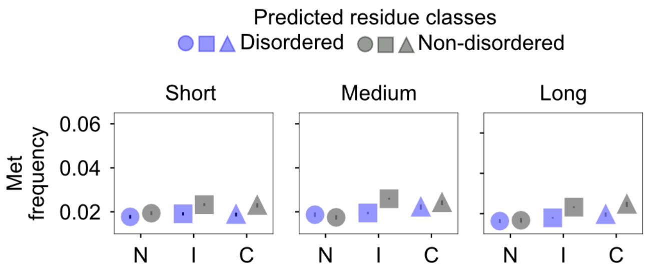

**S8 Fig. Methionine (Met) frequencies after filtering initiation Met.**
Frequencies at terminal (N, C) and internal (I) segments are shown. Note that a given residue at the first position was filtered only if that residue was Met. Otherwise, the residue was retained. All error bars are 95% confidence intervals calculated among 1000 replicates. Replicates were created by sampling genes with replacement. n=3361, 2315 and 2146 genes for the short, medium and long protein length bins, respectively.

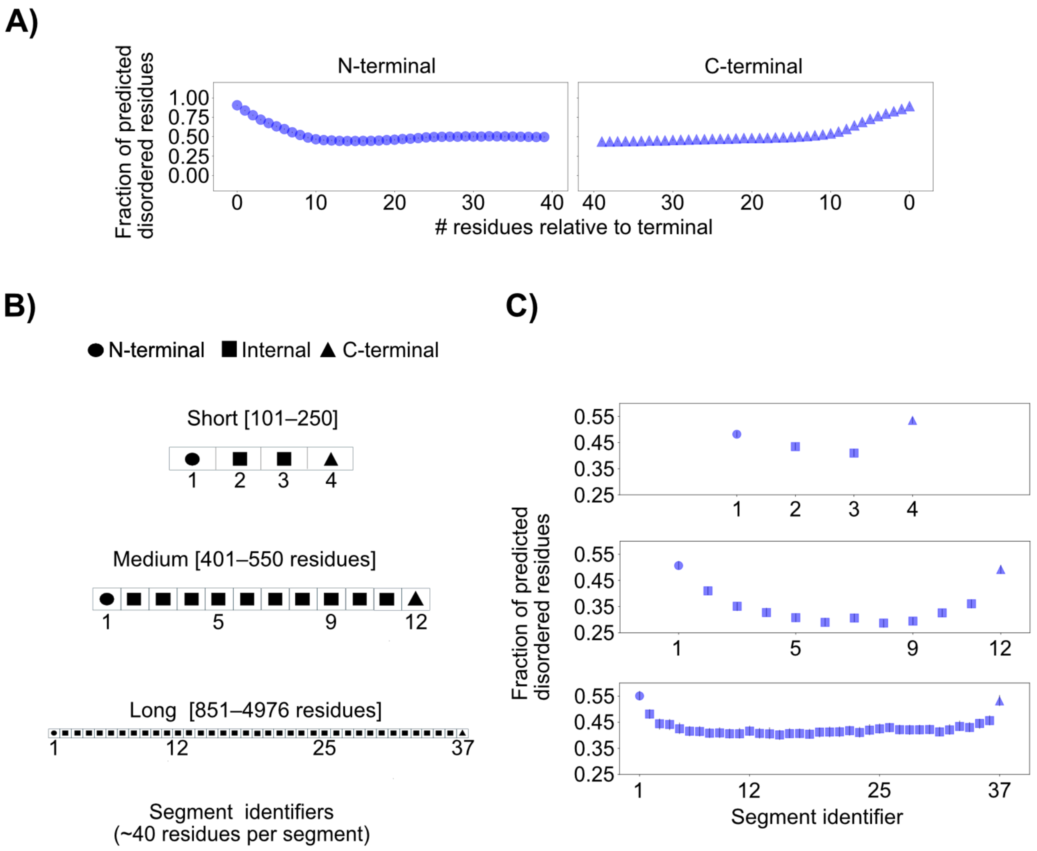

**S9 Fig. Relative solvent accessibility (RSA) values along protein sequences.**
**A)** mean RSA at N- and C- terminal positions. Protein sequences with 80 residues or above are analyzed. **B)** mean RSA for N-terminal, internal and C-terminal segments for each protein length bin. All error bars are 95% confidence intervals calculated among 1000 replicates. Replicates were created by sampling genes with replacement. n=3274, 2171 and 1737 genes for the short, medium and long protein length bins.

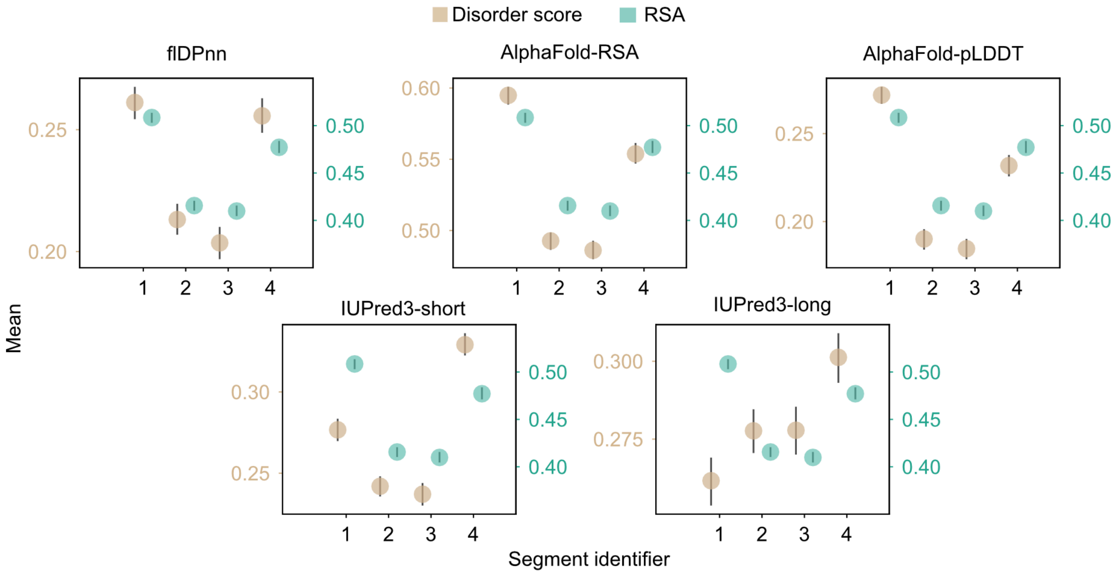

**S10 Fig. Mean disorder scores and RSA values along short protein sequences.** Only protein sequences with data from all disorder prediction approaches were used. All error bars are 95% confidence intervals calculated among 300 replicates. Replicates were created by sampling genes with replacement. n=3274 genes were used.

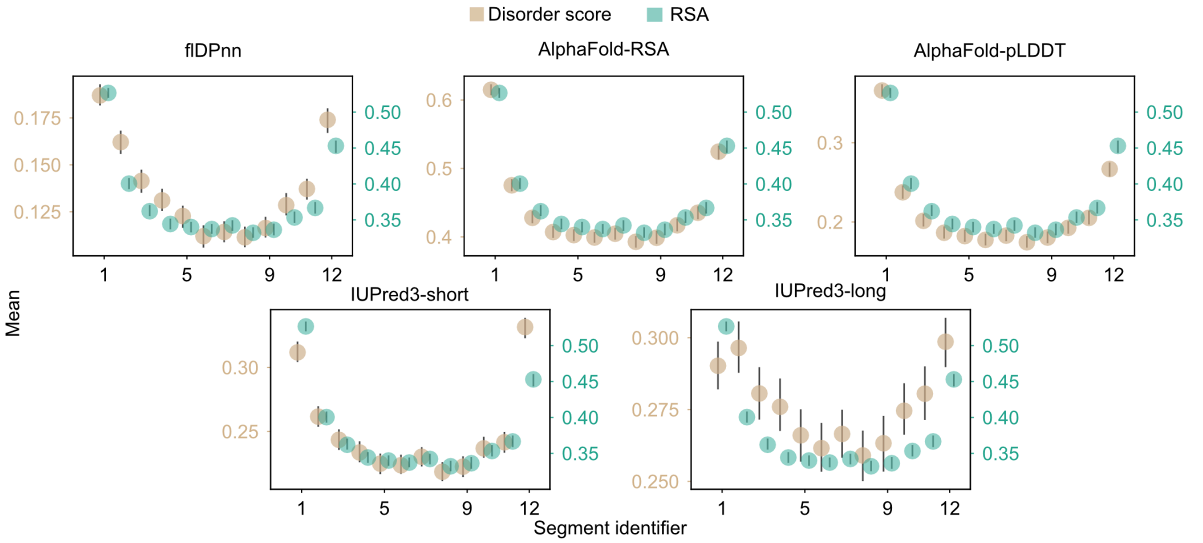

**S11 Fig. Mean disorder scores and RSA values along medium-length protein sequences**. Only protein sequences with data from all disorder prediction approaches were used. All error bars are 95% confidence intervals calculated among 300 replicates. Replicates were created by sampling genes with replacement. n=2171 genes were used.

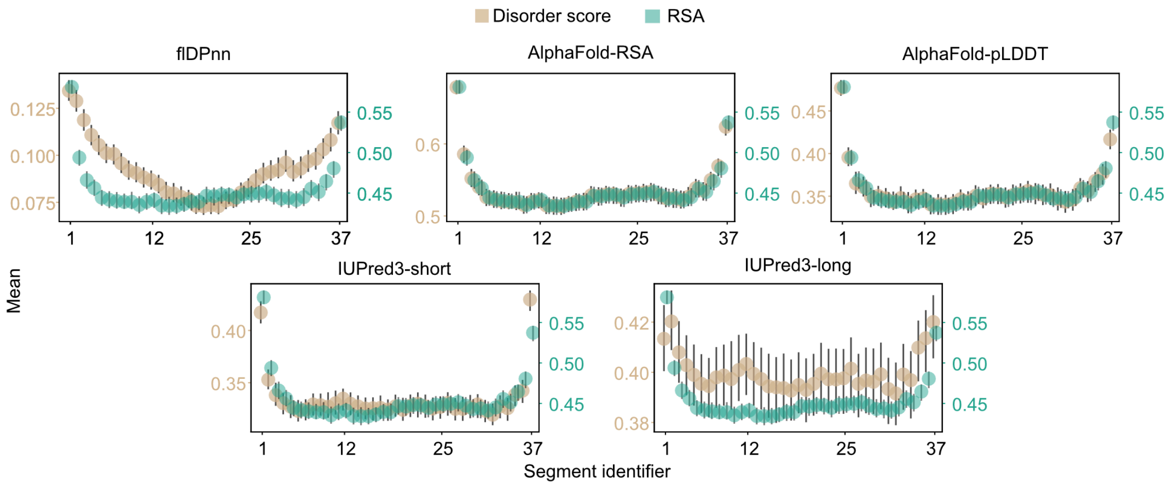

**S12 Fig. Mean disorder scores and RSA values along long-length protein sequences.** Only protein sequences with data from all disorder prediction approaches were used. All error bars are 95% confidence intervals calculated among 300 replicates. Replicates were created by sampling genes with replacement. n=1737 genes were used.

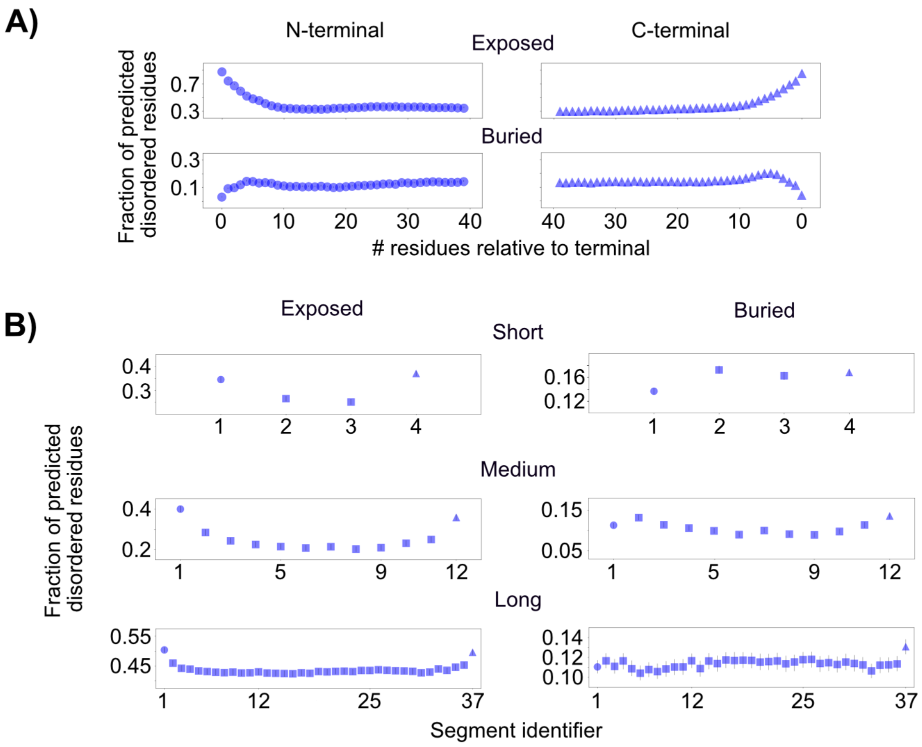

**S13 Fig. Prevalence of intrinsic disorder within terminals and among segments for putative solvent accessibility classes (exposed, buried).**
Values under the non-strict consensus criteria set are shown. Residues were classified, based on relative solvent accessibility (RSA) values, as exposed (RSA≥0.5) and buried (RSA<0.5). The fraction of predicted disordered residues was calculated for each class. **A)** Prevalence of intrinsic disorder at N- and C-terminal positions. Only protein sequences with 80 residues or above were analyzed. n=12472 genes. **B)** Prevalence of intrinsic disorder at terminal and internal segments of protein length bins n=3274, 2171 and 1737 genes for the short, medium and long protein length bins, respectively. In both panels, error bars are 95% confidence intervals calculated among 1000 replicates. Replicates were created by sampling genes with replacement. Statistical tests are presented in S10, S11 and S12 Tables.

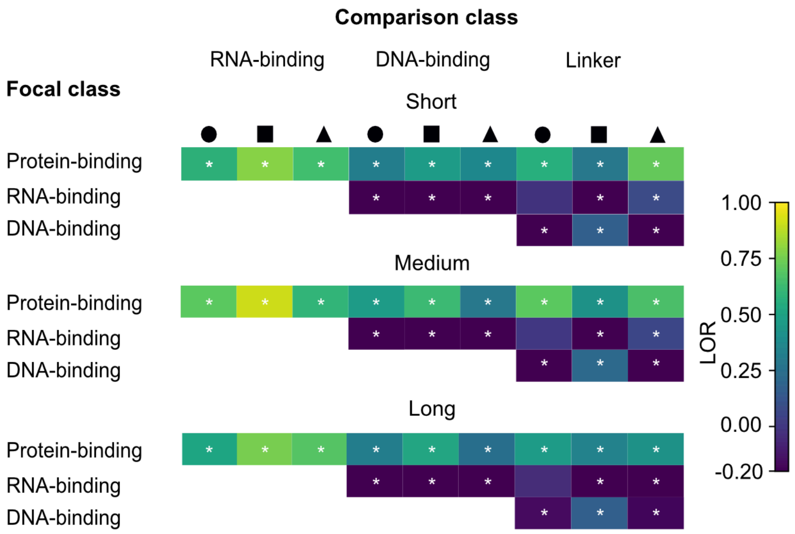

**S14 Fig. Relative prevalence of intrinsic disorder among putative functional classes.** LORs and Z-tests were used to check for differences between "focal" and "comparison" classes, separately for N-terminal (circle), internal (square) and C-terminal (triangle) segments. LOR>0 indicates a higher fraction of predicted disordered residues for the focal class than the comparison class. LOR<0 and LOR=0 indicate a lower fraction and identical equal fractions, respectively. LORs with asterisk are significant at $p<0.05$ and other LORs are non-significant. n=1947, 1851 and 1558 genes for the short, medium and long protein length bins, respectively. Detailed statistical tests results are provided in S13 Table.

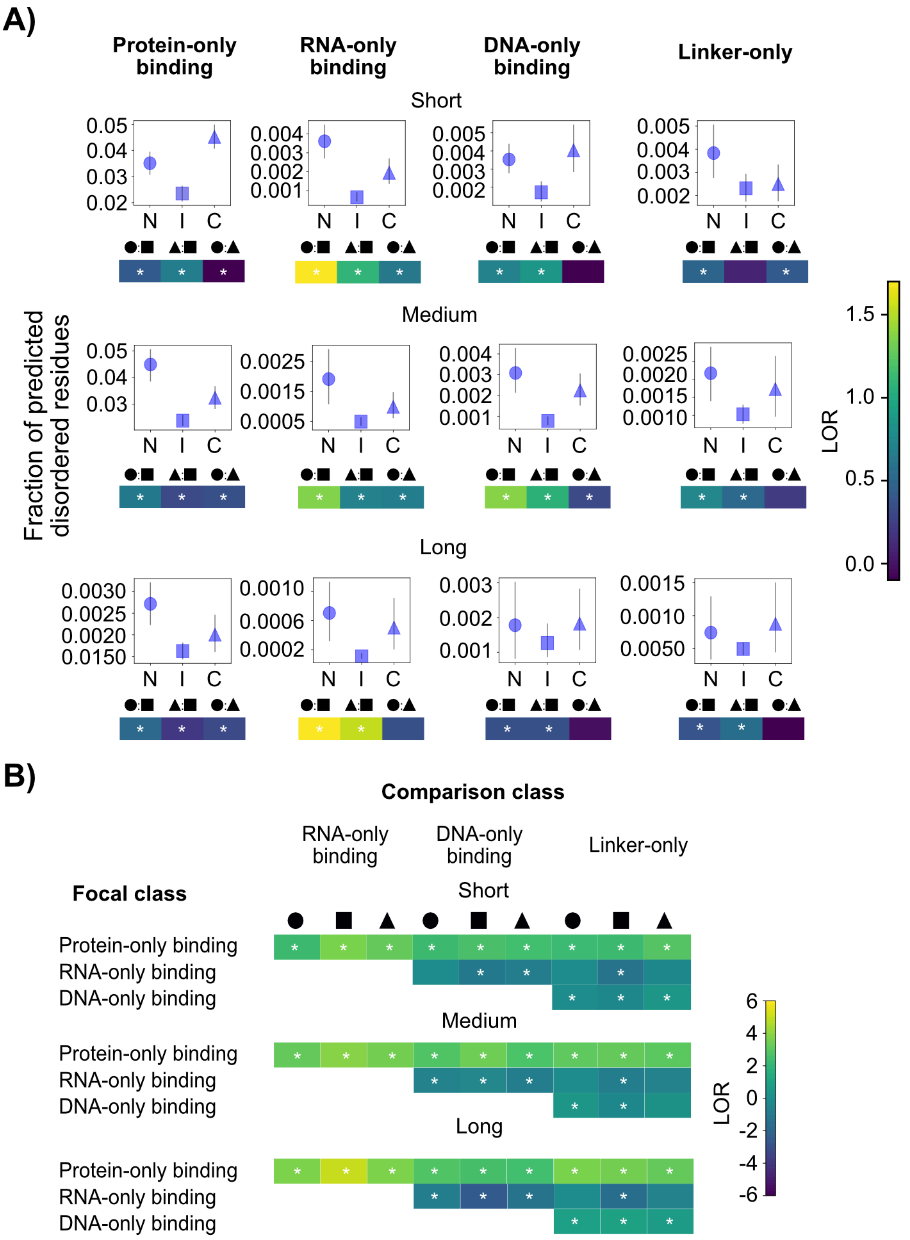

**S15 Fig. Prevalence of single-function classes of intrinsic disorder.**

Analyses are based on the fraction of predicted disordered residues that are exposed (RSA≥0.5) and have been assigned a single function by flDPnn (see *Materials and Methods*). **A)** Prevalence of each class along protein sequences and LOR-based comparisons among terminal (N, C) and internal (I) segment pairings. Error bars are 95% confidence intervals calculated among 500 replicates. Replicates were created by sampling genes with replacement. n=1947, 1851 and 1558 genes for the short, medium and long protein length bins. Given "A:B" symbol, LOR>0 indicates a higher fraction of predicted disordered residues for A than B. LOR<0 and LOR=0 indicate a lower fraction for A and equal fractions, respectively. Asterisk indicates $p<0.05$ for a Z-test. **B)** Relative prevalence of disorder among functional classes. LORs are employed to test differences between "focal vs comparison" classes, separately for each segment. Interpretation of the LOR values follows the same logic as in panel A. In both panels, the Bonferroni sequential method was employed for multiple test corrections among segments and length bins (i.e., nine tests were corrected). Detailed statistical test results are shown in S13 and S14 tables.

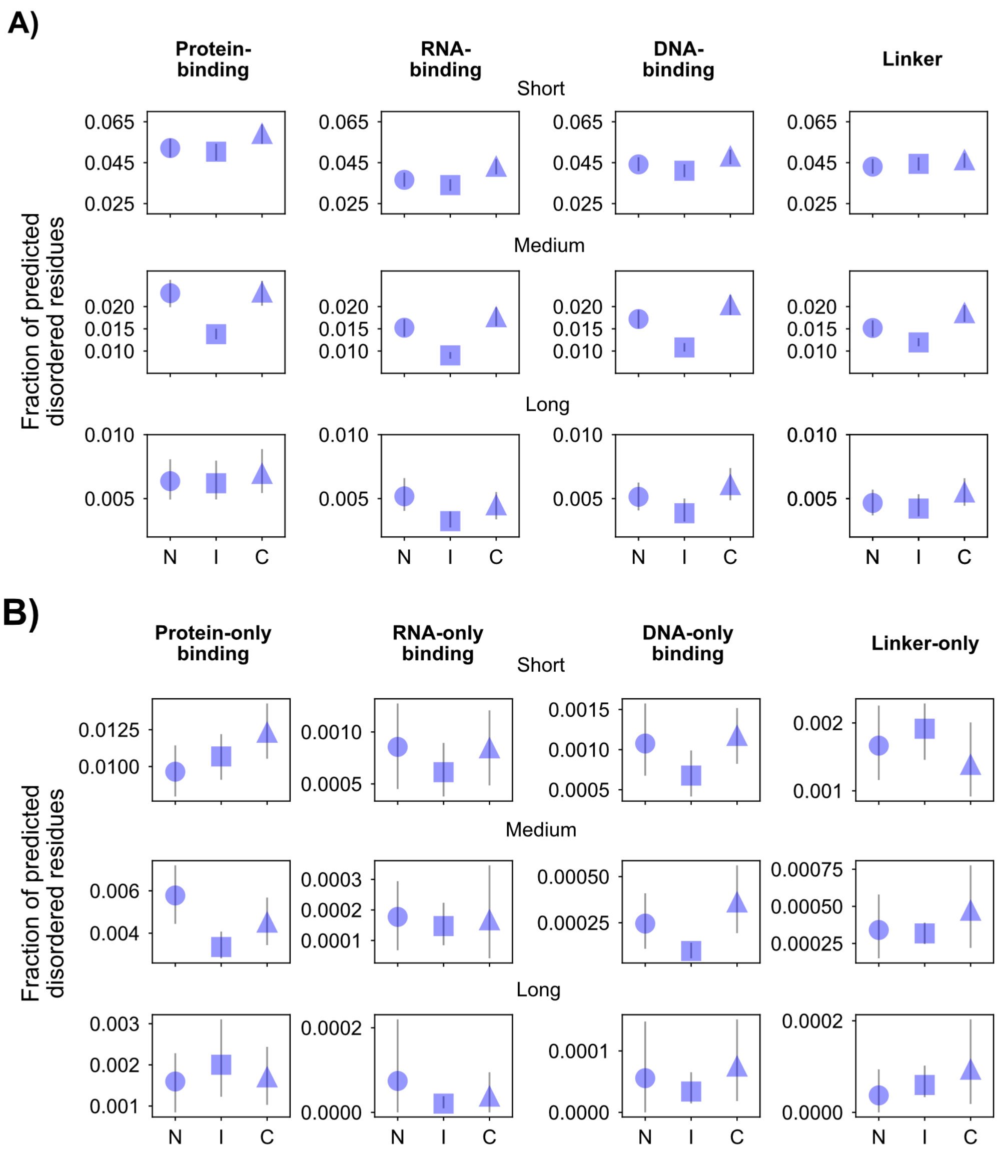

**S16 Fig. Prevalence of intrinsic disorder functional classes in a solvent-buried context.** The fraction of predicted disordered residues for the solvent-buried (RSA<0.5) class, separately for functional class predictions from flDPnn program (see *Materials and Methods*). The fractions are shown for terminal (N, C) and internal (I) segments **A)** Prevalence of each functional class along protein sequences. **B)** Prevalence of single-function classes. In both panels, error bars are 95% confidence intervals calculated among 500 replicates. Replicates were created by sampling genes with replacement. n=1947, 1851 and 1558 genes for the short, medium and long protein length bins, respectively.

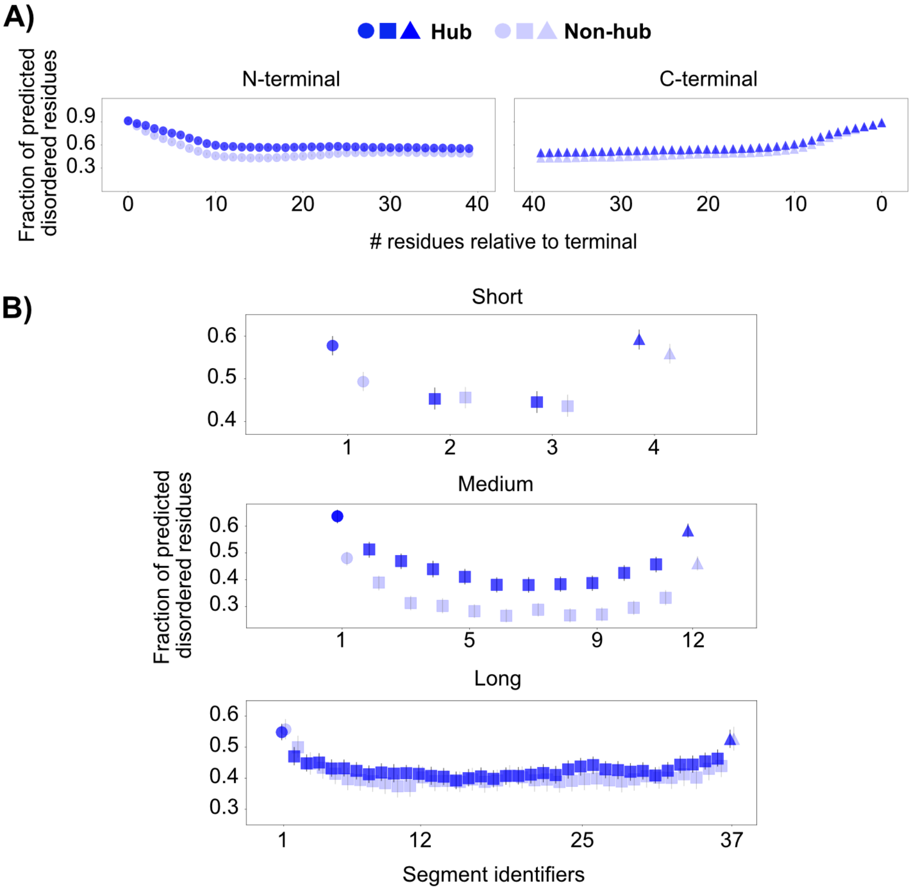

**S17 Fig. Prevalence of intrinsic disorder in protein connectivity classes (hub and non-hub).**

The fraction of predicted disordered residues was calculated separately for hubs (four interactors or above) and non-hubs (below four interactors), as described in *Materials and Methods*. **A)** Prevalence of intrinsic disorder for N- and C- terminal positions of hubs and non-hubs. Only protein sequences with 80 residues or above were analyzed. **B)** Prevalence of intrinsic disorder for terminal (N, C) and internal (I) segments. In both panels, error bars are 95% confidence intervals calculated among 1000 replicates that were created by sampling genes with replacement. n= 943, 805 and 1132 genes for the short, medium and long protein length bins, respectively, for the hub class and n=1043, 800 and 685 for the corresponding bins for the non-hub class. Statistical test results are presented in S15, S16 and S17 Tables.

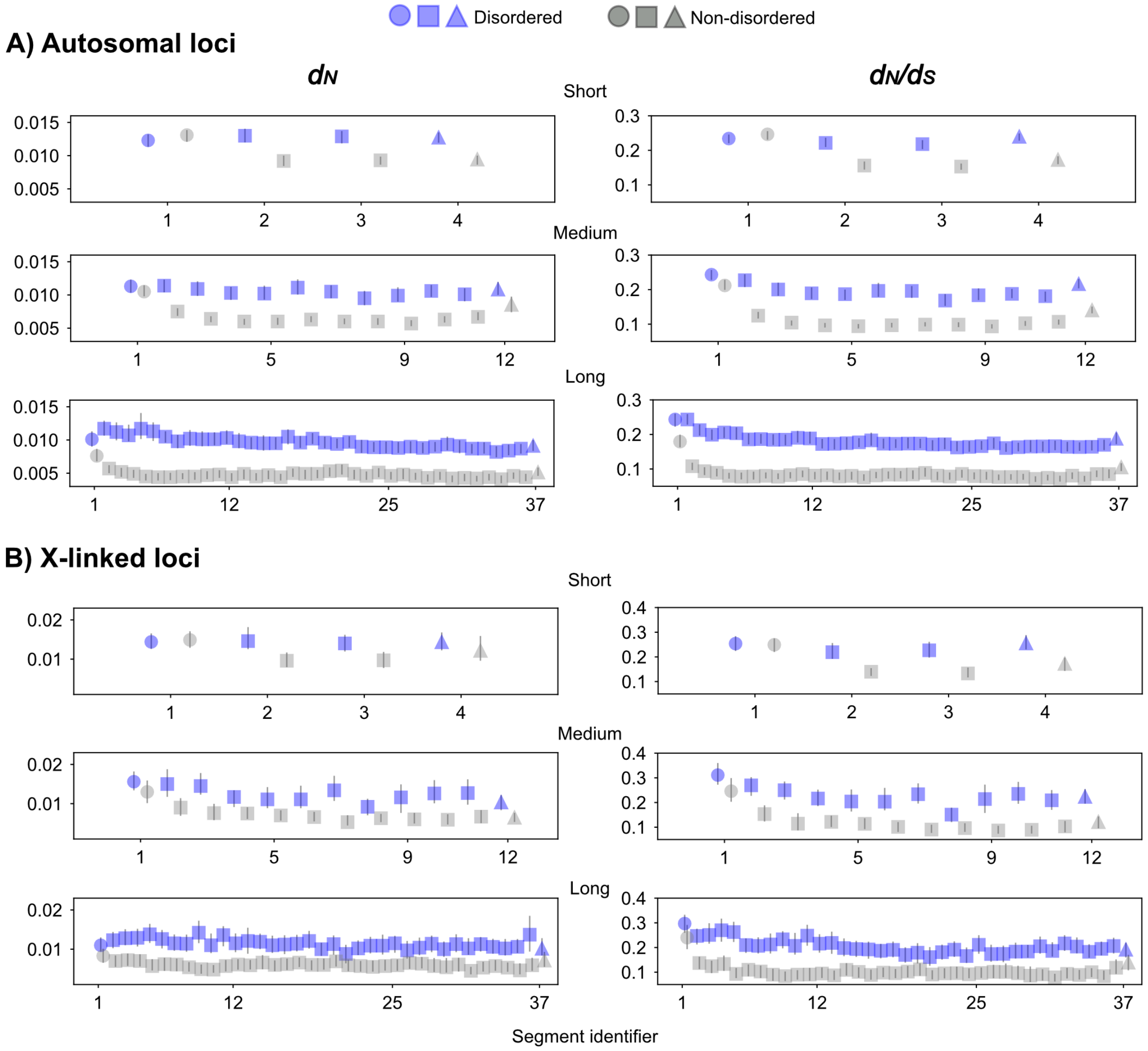

**S18 Fig. Detailed view of $d_N$ and $d_N/d_S$ estimates along protein sequences.**
Estimates of evolutionary parameters were obtained for members of the *D. melanogaster* subgroup and those estimates for the *D. melanogaster* species branch were analyzed (see details in Materials and Methods). $d_N$ and $d_N/d_S$ were calculated under the Yang and Nielsen (1998) codon substitution model implemented in codeml program, separately for N-terminal (N), internal (I) and C-terminal (C) segments. **A)** and **B)** show $d_N$ and $d_N/d_S$ values for autosomal and X-linked loci, respectively, *D. melanogaster* branch. Error bars indicate 95% confidence intervals calculated among 100 replicates that were created by sampling genes with replacement. n=2202, 1694, and 1550 genes for the short, medium and long protein length bins of the autosomal loci and n=378, 282, and 355 genes for the corresponding bins of X-linked loci.

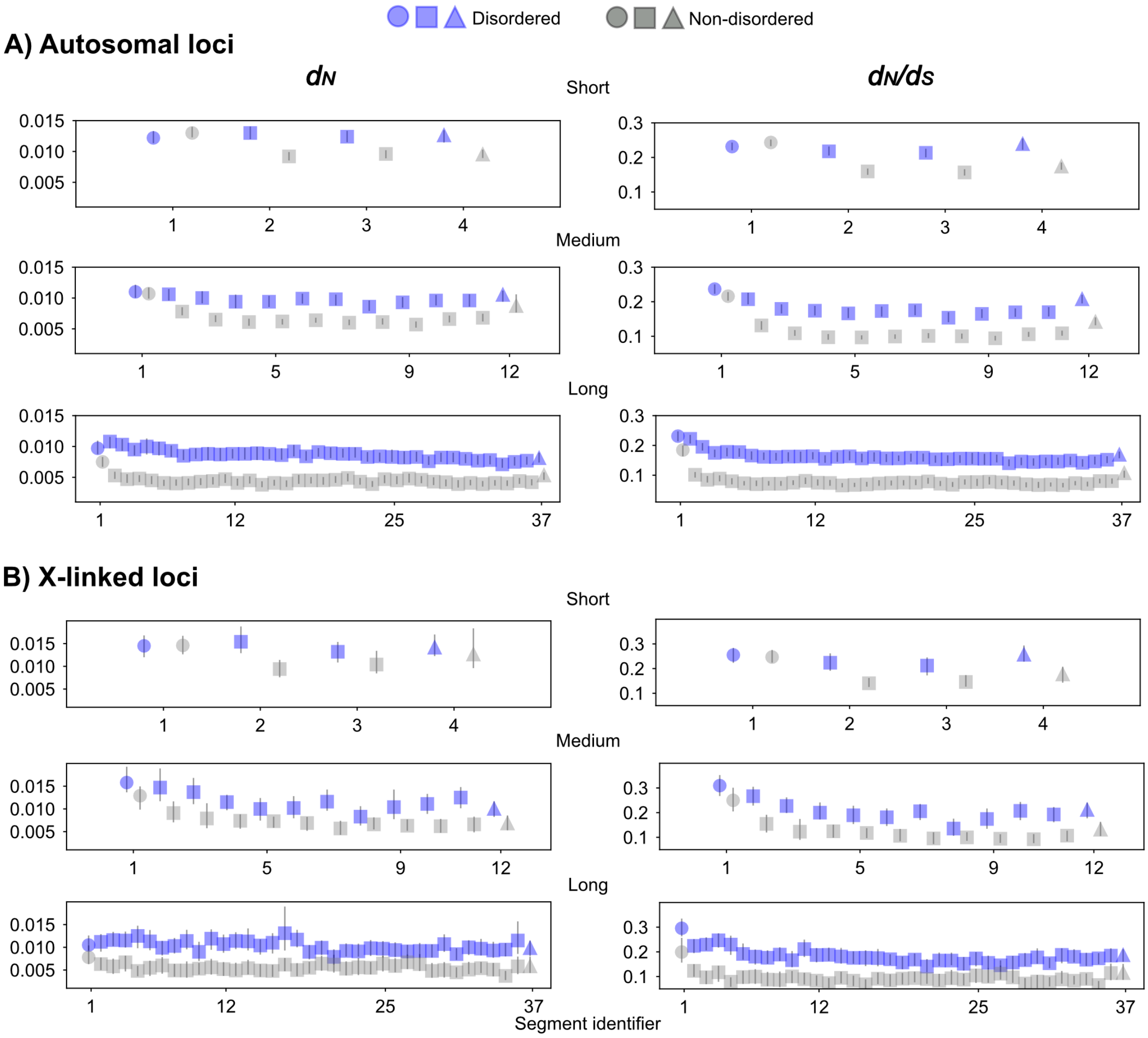

**S19 Fig. Detailed view of $d_N$ and $d_N/d_S$ estimates for residues predicted to be disordered under IUPred3-short.**

Residues were classified as disordered and non-disordered if the IUPred-short disorder score was 0.5 or above and below 0.5, respectively. Estimates of evolutionary parameters were obtained for members of the *D. melanogaster* subgroup and those estimates for the *D. melanogaster* species branch were analyzed (see details in Materials and Methods). $d_N$ and $d_N/d_S$ were calculated under the Yang and Nielsen (1998) codon substitution model implemented in codeml program, separately for N-terminal (N), internal (I) and C-terminal (C) segments.. **A)** and **B)** show $d_N$ and $d_N/d_S$ values for autosomal and X-linked loci, respectively, *D. melanogaster* branch. Error bars indicate 95% confidence intervals calculated among 100 replicates that were created by sampling genes with replacement. n=2202, 1694, and 1550 genes for the short, medium and long protein length bins of the autosomal loci and n=378, 282, and 355 genes for the corresponding bins of X-linked loci.

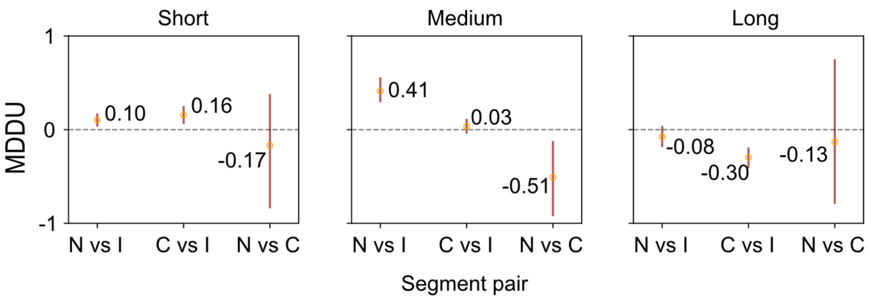

**S20 Fig. Magnitude of disorder distribution unevenness (MDDU) between Actual and Shuffled data.**

MDDU was employed to test Actual and Shuffled data for differences among segment pairings (N-terminal [N], internal [I], and C-terminal [C]) for the fraction of predicted disordered residues (explained in S11 Text). MDDU>0 indicates a greater extent of difference in the Actual data. MDDU<0 and MDDU=0 would indicate lesser and identical extents of difference, respectively. Error bars are 95% confidence intervals from 1000 bootstrap replicates that were created by sampling genes with replacement. n=2005, 1981 and 1956 genes for the short, medium and long protein length bins, respectively. Note that difference statistics are used for MDDU calculation and can result in large variance.

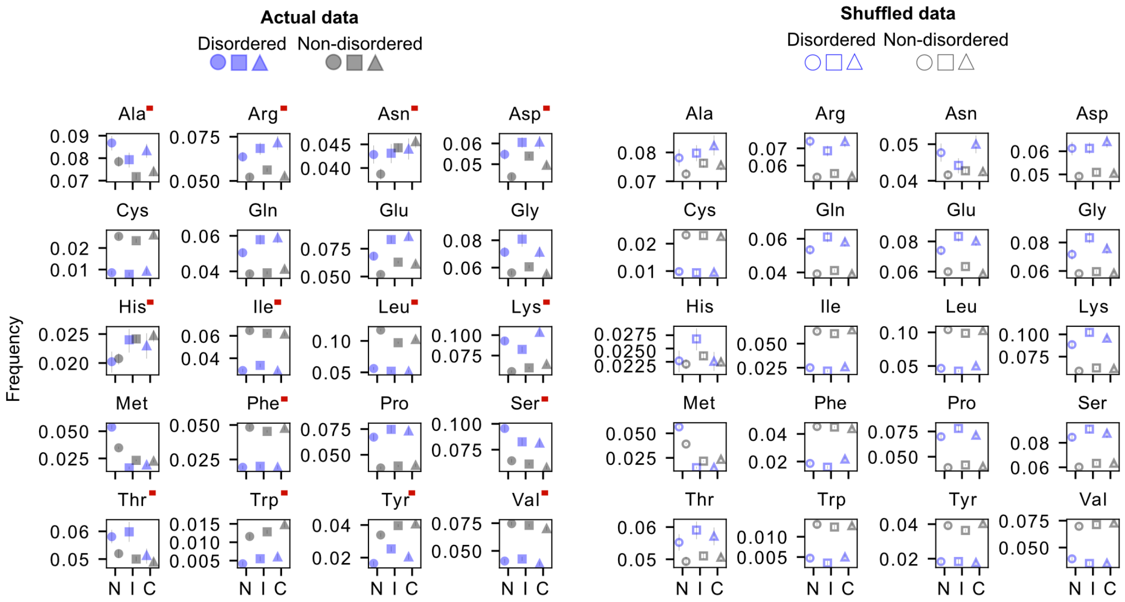

**S21 Fig. Amino acid frequencies in Actual and Shuffled data for the short protein length bin.**

Data for N-terminal (N), internal (I) and C-terminal (C) segments are shown. All error bars are 95% confidence intervals calculated among 1000 replicates created by sampling genes with replacement. n=2005 genes. Red annotations specify cases where amino acid distributions differ (visually) between the Actual and Shuffled data.

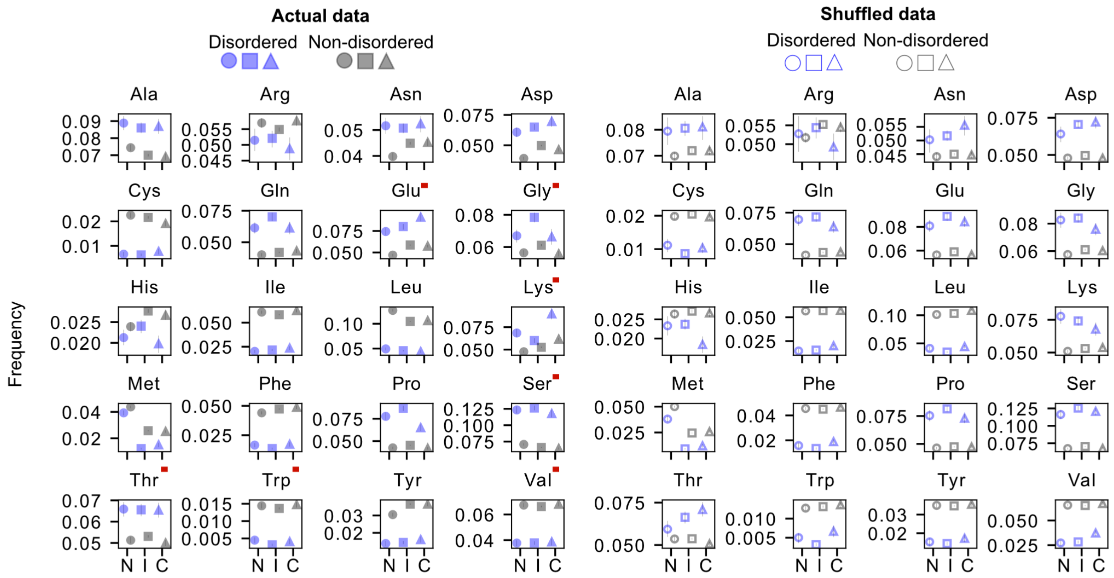

**S22 Fig. Amino acid frequencies in Actual and Shuffled data in the medium protein length bin.**

Data for N-terminal (N), internal (I) and C-terminal (C) segments are shown. All error bars are 95% confidence intervals calculated among 1000 replicates created by sampling genes with replacement. n=1981 genes. Red annotations specify cases where amino acid distributions differ (visually) between the Actual and Shuffled data.

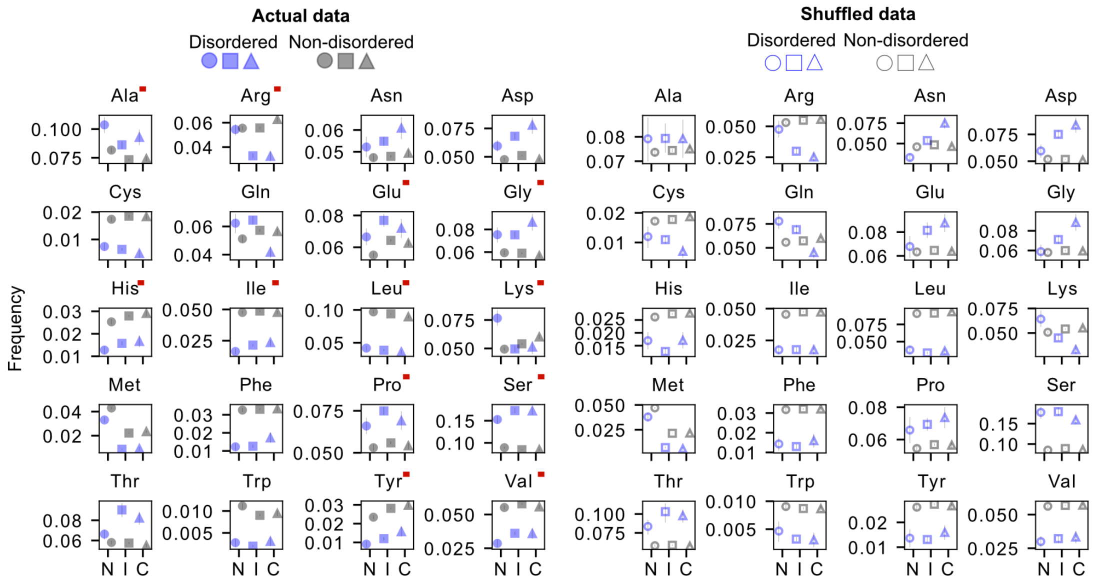

**S23 Fig. Amino acid frequencies in Actual and Shuffled data for the long protein length bin.**

Data for N-terminal (N), internal (I) and C-terminal (C) segments are shown. All error bars are 95% confidence intervals calculated among 1000 replicates created by sampling genes with replacement. n=1956 genes. Red annotations specify cases where amino acid distributions differ (visually) between the Actual and Shuffled data.

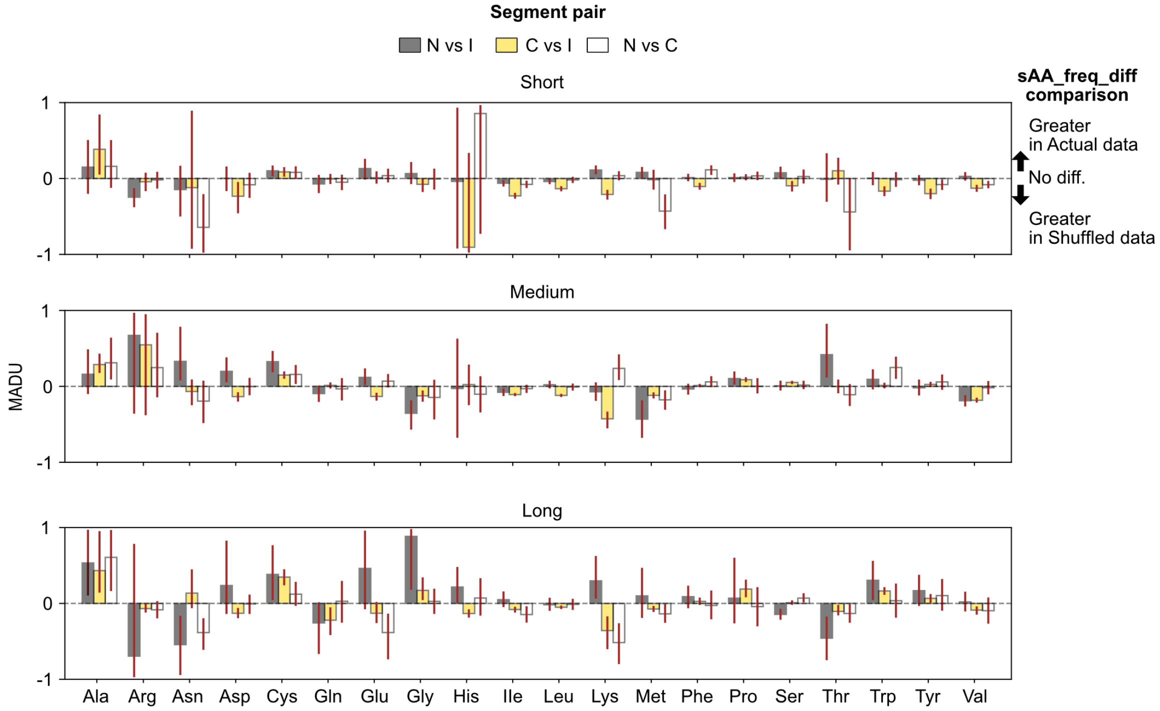

**S24 Fig. Magnitude of amino acid distribution unevenness (MADU) in Actual and Shuffled data.**

MADU was employed to quantify differences between the Actual and Shuffled data for amino acid frequencies at disordered residues along proteins, while accounting for frequencies at non-disordered residues. MADU is calculated for pairings among terminal (N, C) and internal (I) segments. MADU>0 indicates a greater extent of difference in the Actual data. MADU<0 and MADU=0 would indicate lesser and identical extents of difference, respectively. Error bars are 95% confidence intervals calculated among 1000 bootstrap replicates created by sampling genes with replacement. n=2005, 1981 and 1956 genes for the short, medium and long protein length bins, respectively. Note that difference statistics are used for MADU calculation and can result in large variance.

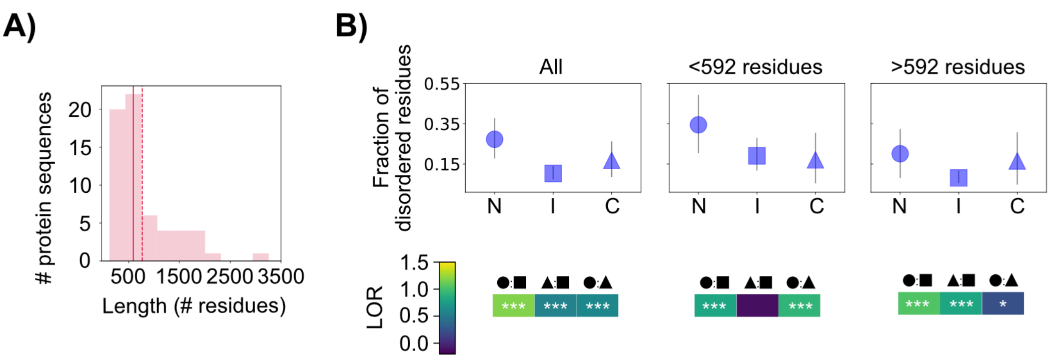

**S25 Fig. Prevalence and distribution of experimentally-determined disordered residues.** Experimentally-validated annotations for intrinsic disorder were obtained from DisProt database. Analyses are based on annotations for *D. melanogaster* protein sequences (n=62. S7 and S9 Texts). **A**) Distribution of lengths for proteins with intrinsic disorder annotations. Solid and dashed lines specify mean (787.9) and median (592) residue counts. The latter count was used as a threshold for binning data into short and longer proteins. **B)** Disorder prevalence along segments. The fraction of disordered residues are shown. Error bars are 95% confidence intervals based on 1000 bootstrap replicates created by sampling protein sequences with replacement. n=62, 31, and 31 protein sequences for the "All", "<592" and ">592" datasets. LORs were calculated and tested for differences among N-terminal (N or circle), internal (I or square) and C-terminal (C or triangle) segments. Given "A:B" symbol, LOR>0 indicates a higher fraction for A than B. LOR<0 and LOR=0 would indicate a lower and identical fractions, respectively.

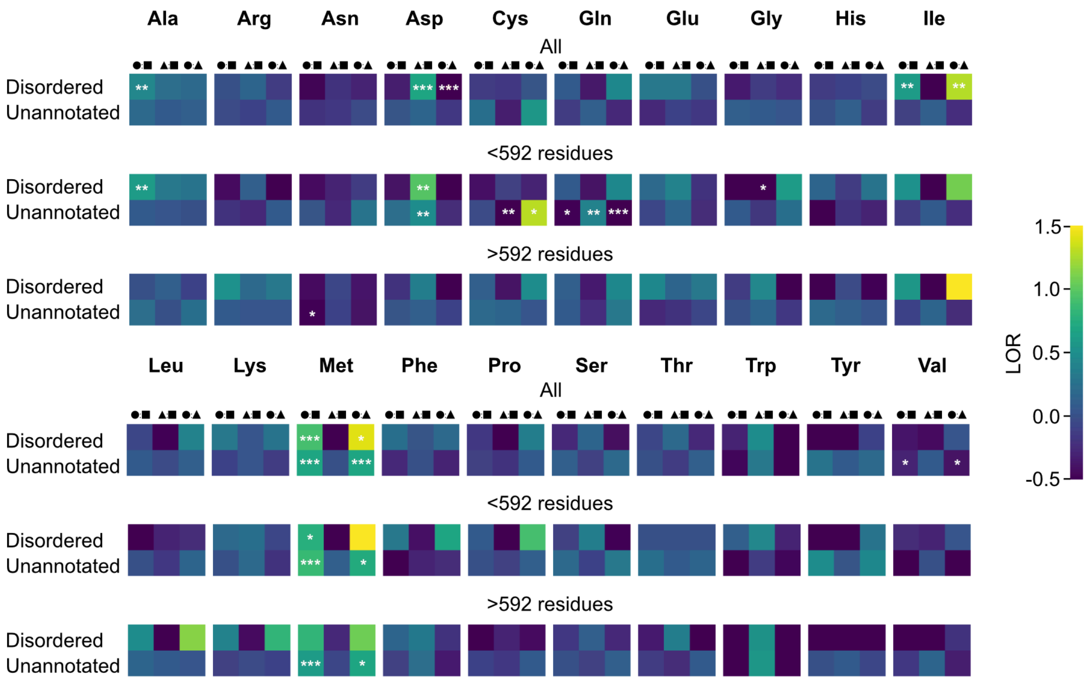

**S26 Fig. Log-odds ratios (LORs) for amino acid frequency comparisons between segments among experimentally-determined disordered residues.**
Intrinsic disorder annotations for *D. melanogaster* were obtained from DisProt database (S7 and S9 Texts). LORs were employed to compare pairings of N-terminal (circle), internal (square), and C-terminal (triangle) segments for amino acid frequencies among disordered residues. Results for cases with disorder annotation – “Unannotated”, are also presented. Note that such annotations could include both disordered and non-disordered residues and are thus treated as absent data. Given “A:B” symbol, LORs>0 and <0 indicate higher and lower fractions, respectively in segment A. Protein sequence counts are shown. n=62, 31, and 31 protein sequences for the “All”, “<592” and “>592” datasets.

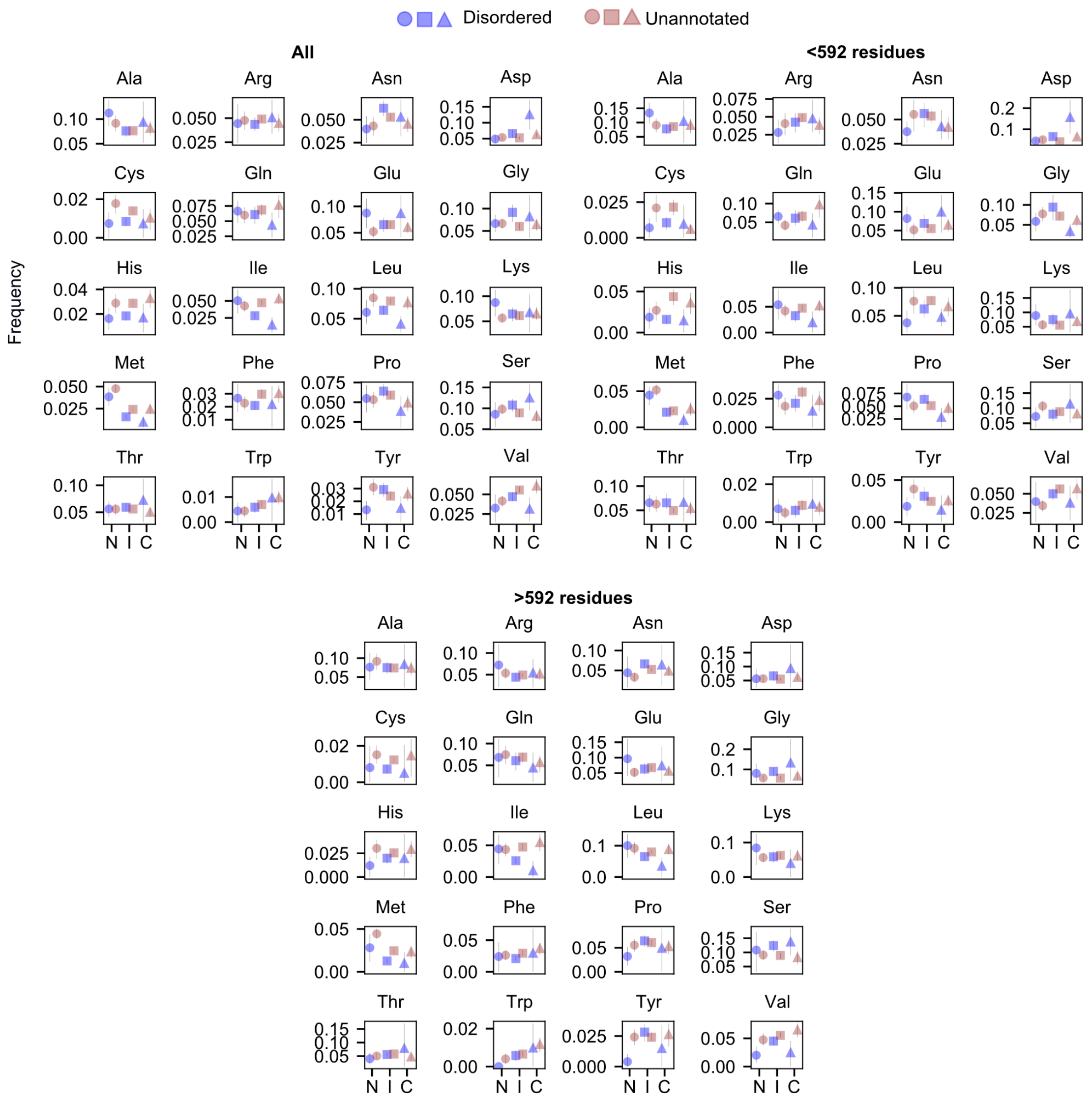

**S27 Fig. Amino acid frequencies for experimentally-determined disordered residues.** Intrinsic disorder annotations for *D. melanogaster* were obtained from DisProt database (S7 and S9 Texts). Amino acid frequencies for the resulting annotations were analyzed, separately for N-terminal (N), internal (I) and C-terminal (C) segments. Analyses are based on all protein sequences and bins for which protein sequences contain below 592 residues ("<592") or above (>592 residues). Frequencies for unannotated residues are included, but note that such cases may include both disordered and non-disordered residues and thus treated as absent data. All error bars specify 95% confidence intervals calculated among 1000 replicates by sampling protein sequences with replacement. n=62, 31, and 31 protein sequences for the "All", "<592" and ">592" datasets.

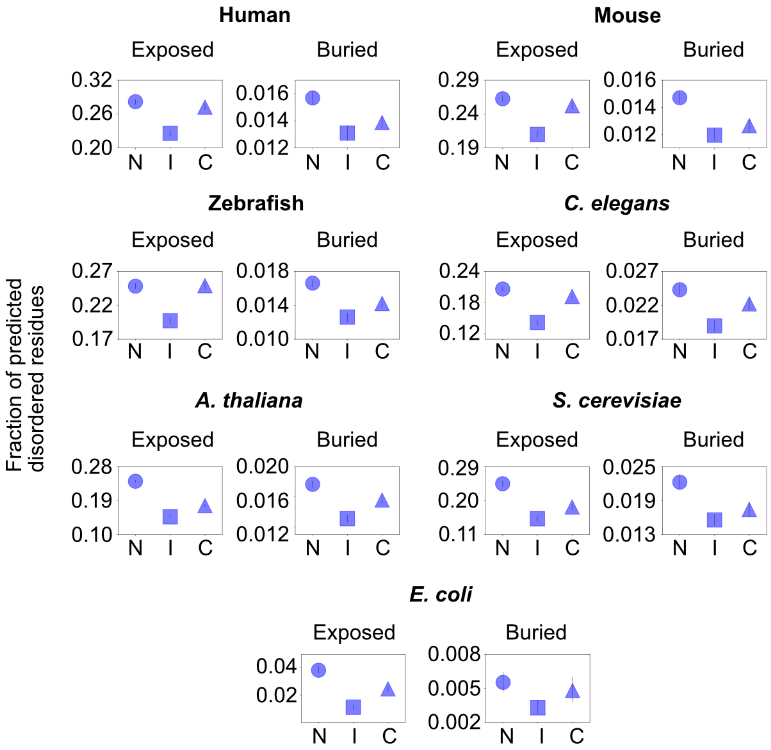

**S28 Fig. AlphaFold-pLDDT-based predictions for intrinsic disorder prevalence in non-*Drosophila* species.**
AlphaFold-pLDDT approach was employed to assign predicted disordered residues (≥0.5 cut-off) for protein sequences obtained from AlphaFold protein structure database (details in S8 and S12 Texts). The resulting assignments were assigned as solvent-exposed (RSA≥0.5) or solvent-buried (RSA<0.5). Protein sequences were split such that the first and last 40 residues corresponded to N-terminal (N) and C-terminal (C) segments, respectively, and the remaining residues were used as the internal (I) segment. Error bars are 95% confidence intervals calculated among 500 replicates created by sampling protein sequences with replacement. n=20227, 21592, 24664, 19690, 27370, 5973 and 4321 protein sequences for Human, Mouse, Zebrafish, *C. elegans*, *A. thaliana*, *S. cerevisiae* and *E. coli*, respectively. Descriptive statistics for protein sequences are shown in S26 Table.

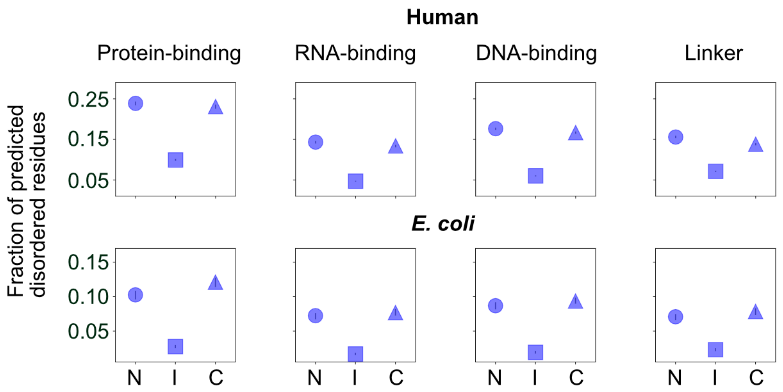

**S29 Fig. Prevalence of putative functional classes of intrinsic disorder in human and *E. coli*.**

Disorder function predictions were obtained from flDPnn program. Note that a given residue can have multiple putative functions (S8 and S12 Texts). The fraction of predicted disordered residues was analyzed for each class. Protein sequences were split such that the first and last 40 residues corresponded to N-terminal (N) and C-terminal (C) segments, respectively, and the remaining residues were used as the internal (I) segment. Error bars are 95% confidence intervals calculated among 100 replicates created by sampling protein sequences with replacement. n=19222 and 3592 protein sequences for human and *E. coli*, respectively.

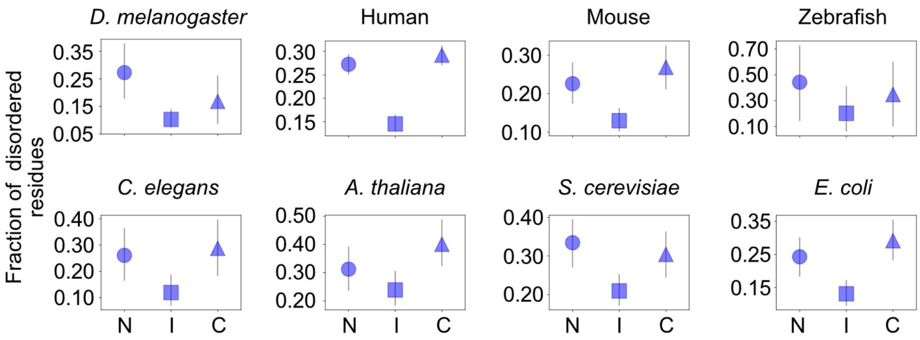

**S30 Fig. Distribution of experimentally-determined disordered residues in *D. melanogaster* and seven non-*Drosophila* species.**

Intrinsic disorder annotations were obtained from the DisProt database (S7 and S8 Text). Protein sequences were split such that the first and last 40 residues corresponded to N-terminal (N) and C-terminal (C) segments, respectively, and the remaining residues were used as the internal (I) segment. Error bars are 95% confidence intervals based on 1000 bootstrap replicates created by sampling protein sequences with replacement. n=62, 1249, 195, 10, 54, 113, 206, 141 protein sequences for *D. melanogaster*, Human, Mouse, Zebrafish, *C. elegans*, *A. thaliana*, *S. cerevisiae* and *E. coli*, respectively.

#### Supplementary Tables

**S1 Table.** Descriptive statistics for disorder scores at experimentally-determined disordered residues.

| Approach | # protein sequences[1] | # disordered residues | Disorder scores | | | | | |
|---|---|---|---|---|---|---|---|---|
| | | | min | 25th percentile | mean | median | 75th percentile | max |
| flDPnn | 57 | 5,062 | 0.003 | 0.162 | 0.363 | 0.333 | 0.553 | 0.982 |
| AlphaFold-RSA | 57 | 4,783 | 0.060 | 0.597 | 0.716 | 0.770 | 0.887 | 0.968 |
| AlphaFold-pLDDT | 57 | 4,783 | 0.011 | 0.316 | 0.468 | 0.537 | 0.641 | 0.811 |
| IUPred3-short | 62 | 5,473 | 0.001 | 0.365 | 0.490 | 0.502 | 0.641 | 1.000 |
| IUPred3-long | 62 | 5,478 | 0.010 | 0.435 | 0.576 | 0.590 | 0.751 | 0.980 |

[1]Intrinsic disorder annotations were obtained for all *D. melanogaster* protein sequences in DisProt (n=62). For flDPnn, protein sequences were excluded if they had low quality predictions. For AlphaFold-based approaches, protein sequences were excluded if they could not be successfully linked to protein structures obtained from AlphaFold protein structure database.

**S2 Table**. Counts of protein sequences used to obtain disorder predictions.

| | | Intrinsic disorder prediction approach | | | | |
|---|---|---|---|---|---|---|
| | | flDPnn[1] | AlphaFold-RSA | AlphaFold-pLDDT | IUPred3-short[2] | IUPred3-long[2] |
| Input[a] | # sequences | 13,937 | 12,922 | 12,922 | 13,937 | 13,937 |
| | # residues | 7,455,625 | 6,188,227 | 6,188,227 | 7,455,625 | 7,455,625 |
| Successfully Processed[b] | # sequences | 13,917 | 12,922 | 12,922 | 13,930 | 13,930 |
| | # residues | 7,281,396 | 6,188,227 | 6,188,227 | 7,455,531 | 7,455,531 |
| Retained[c] | # sequences | 10,629 | 12,922 | 12,922 | 13,930 | 13,930 |
| | # residues | 6,255,769 | 6,188,227 | 6,188,227 | 7,431,738 | 7,453,576 |
| Data coverage | # sequences | 76.265 | 92.717 | 92.717 | 99.950 | 99.950 |
| | # residues | 83.907 | 83.001 | 83.001 | 99.680 | 99.973 |

[a]Note that 13937 protein sequences were employed, but only 12922 protein structures from AlphaFold protein structure database were successfully linked to these sequences (See details in *Materials and Methods*).
[b]For each program, a given process is considered successful if the program returns disorder scores for each residue.
[c] Values are relative to total data (i.e., 13937 protein sequences and 7455625 residues).
[1]3308 successfully processed cases were excluded due to low quality predictions as indicated in flDPnn outputs.
[2]Residues that were assigned unexpected values (below zero or above one) were excluded.

**S3 Table.** Counts of disordered residues under consensus criteria sets.

| Consensus criteria set | DisProt[1] | | | Genome-wide[2] | | |
|---|---|---|---|---|---|---|
| | # sequences | # true positives | # false negatives | # genes | # predicted disordered | # predicted non-disordered |
| Non-strict | 57 | 4,514 | 264 | 13,910 | 3,020,006 | 4,261,296 |
| Strict | 57 | 3,218 | 1,560 | 13,910 | 854,025 | 6,427,277 |

[1]4877 residues were classified.
[2]The longest protein isoform for each gene was used. 7281302 residues were classified.

**S4 Table.** Descriptive statistics for predicted disordered segments.

| Consensus criteria set | # genes[a] | # segments[b] | Predicted segment length (# residues) | | | | | |
|---|---|---|---|---|---|---|---|---|
| | | | min | 25th percentile | mean | median | 75th percentile | max |
| Non-strict | 13,910 | 93,181 | 1 | 2 | 32.410 | 7 | 23 | 2,334 |
| Strict | 13,910 | 50,391 | 1 | 3 | 16.948 | 7 | 21 | 633 |

[a]The longest protein isoform for each gene was used.
[b]Stretches of residues predicted to be disordered or individual residues predicted to be disordered but not adjacent to other such residues.

**S5 Table.** Descriptive statistics for protein length bins used in downstream analysis.

| Data/Bin | # genes[1] | # residues | | | | | |
|---|---|---|---|---|---|---|---|
| | | total | min | mean | median | max | mean per segment identifier[2] |
| All | 13,910 (13746) | 7,281,302 | 19 | 523.46 | 391 | 4,976 | |
| Bin 1 | 836 (827) | 60,921 | 19 | 72.87 | 77 | 100 | |
| Bin 2[a] | 3,361 (3245) | 584,220 | 101 | 173.82 | 173 | 250 | 43.58 |
| Bin 3 | 2,931 (2901) | 944,452 | 251 | 322.23 | 322 | 400 | |
| Bin 4[a] | 2,315 (2312) | 1,094,181 | 401 | 472.65 | 473 | 550 | 39.43 |
| Bin 5 | 1,505 (1500) | 932,860 | 551 | 619.84 | 615 | 700 | |
| Bin 6 | 816 (816) | 626,574 | 701 | 767.86 | 764 | 850 | |
| Bin 7[a] | 2,146 (2145) | 3,038,094 | 851 | 1415.7 | 1209 | 4,976 | 38.26 |

[a] Used as representative bins for short, medium or long protein sequences.

[1] The longest protein isoform for each gene was used. Values between parentheses specify the count of unique protein sequences.

[2]Protein sequences within the representative bins were split into segments to have roughly 40 residues per segment. Short, medium and long proteins were split into 4, 12 and 37 segments, respectively.

**S6 Table.** Spearman's rank correlation coefficients ($r_s$) for the fraction of predicted disordered residues along protein terminals.

| Consensus criteria set | # genes[1] | # positions[2] | Terminal | $r_s$[3] |
|---|---|---|---|---|
| Non-strict | 13,461 | 40 | N | $-0.169^{ns}$ |
| | | | C | $-0.999^{***}$ |
| Strict | 13,461 | 40 | N | $-0.725^{***}$ |
| | | | C | $-0.993^{***}$ |

[1]Only protein sequences containing 80 residues or more were used.
[2]First or last 40 residues in proteins.
[3]The fraction of predicted disordered residues at each position was calculated and the resulting values were used for $r_s$ calculation. The Bonferroni sequential method was used for multiple test corrections for each consensus criteria set (i.e., two tests were corrected). Superscripts ns and *** indicate non-significant and $p<0.001$, respectively. Note that for all cases, $r_s>0.9$ at $p<0.001$ when only the first 20 residues are analyzed.

**S7 Table.** Log-odds ratios (LOR) for differences among protein terminal (N, C) and internal segments for the fraction of predicted disordered residues.

| Predicted residue class | Protein length bins[1] | | | | | | | | |
|---|---|---|---|---|---|---|---|---|---|
| | Short | | | Medium | | | Long | | |
| | N | I | C | N | I | C | N | I | C |
| Disordered | 70,577 | 123,660 | 77,434 | 46,262 | 297,528 | 44,268 | 45,211 | 1,209,043 | 43,376 |
| Non-disordered | 75,905 | 169,304 | 67,340 | 45,028 | 615,372 | 45,723 | 36,914 | 1,665,332 | 38,218 |
| $LOR_{NI}$[a] | 0.241*** | | | 0.754*** | | | 0.523*** | | |
| $LOR_{CI}$[a] | 0.454*** | | | 0.694*** | | | 0.447*** | | |
| $LOR_{NC}$[a] | -0.212*** | | | 0.060*** | | | 0.080*** | | |

[1]Residue count ranges and gene counts for each bin: Short ([101-250], 3361), Medium ([401-550], 2315), Long ([851,4976], 2146). Residue counts for each segment are shown. Note that counts for a given segment are pooled among protein sequences.
[a]Extent of difference between segments for the fraction of predicted disordered residues. Subscript format "AB" specifies the segments being compared. $LOR_{AB}>0$ indicates a higher fraction at segment A than B. $LOR_{AB}<0$ indicates a lower fraction at segment A and $LOR_{AB}=0$ indicates identical fractions. Asterisks specify significance levels for Z-statistics of odds ratios. The Bonferroni sequential method was employed for multiple test corrections across bins (i.e., nine tests were corrected for each segment pairing). Superscript *** indicates $p<0.001$.

**S8 Table.** Log-odds ratios (LOR) for amino acid frequency differences among protein terminal (N, C) and internal (I) for predicted *disordered* residues.

| AA | Protein length bin[1] | | | | | | | | |
|---|---|---|---|---|---|---|---|---|---|
| | Short | | | Medium | | | Long | | |
| | LOR$_{NI}$[2] | LOR$_{CI}$[2] | LOR$_{NC}$[2] | LOR$_{NI}$[2] | LOR$_{CI}$[2] | LOR$_{NC}$[2] | LOR$_{NI}$[2] | LOR$_{CI}$[2] | LOR$_{NC}$[2] |
| Ala | 0.037$^{ns}$ | 0.0$^{ns}$ | 0.038$^{ns}$ | 0.003$^{ns}$ | 0.009$^{ns}$ | -0.006$^{ns}$ | 0.128$^{***}$ | -0.013$^{ns}$ | 0.141$^{***}$ |
| Arg | -0.040$^{ns}$ | 0.073$^{***}$ | -0.113$^{***}$ | 0.008$^{ns}$ | -0.013$^{ns}$ | 0.022$^{ns}$ | 0.087$^{***}$ | 0.107$^{***}$ | -0.020$^{ns}$ |
| Asn | -0.029$^{ns}$ | 0.060$^{*}$ | -0.089$^{**}$ | -0.001$^{ns}$ | 0.052$^{ns}$ | -0.053$^{ns}$ | 0.015$^{ns}$ | 0.078$^{**}$ | -0.064$^{ns}$ |
| Asp | -0.084$^{***}$ | -0.025$^{ns}$ | -0.059$^{ns}$ | -0.085$^{***}$ | 0.031$^{ns}$ | -0.115$^{***}$ | -0.019$^{ns}$ | 0.019$^{ns}$ | -0.038$^{ns}$ |
| Cys | 0.042$^{ns}$ | 0.190$^{***}$ | -0.148$^{*}$ | 0.062$^{ns}$ | 0.139$^{*}$ | -0.077$^{ns}$ | 0.109$^{ns}$ | 0.178$^{**}$ | -0.069$^{ns}$ |
| Gln | -0.119$^{***}$ | -0.033$^{ns}$ | -0.086$^{**}$ | -0.189$^{***}$ | -0.118$^{***}$ | -0.071$^{*}$ | -0.184$^{***}$ | -0.183$^{***}$ | -0.001$^{ns}$ |
| Glu | -0.149$^{***}$ | -0.053$^{**}$ | -0.095$^{**}$ | -0.173$^{***}$ | 0.016$^{ns}$ | -0.189$^{***}$ | -0.135$^{***}$ | -0.109$^{***}$ | -0.026$^{ns}$ |
| Gly | -0.112$^{***}$ | -0.044$^{ns}$ | -0.067$^{*}$ | -0.140$^{***}$ | -0.110$^{***}$ | -0.030$^{ns}$ | 0.001$^{ns}$ | 0.004$^{ns}$ | -0.003$^{ns}$ |
| His | -0.165$^{***}$ | -0.001$^{ns}$ | -0.164$^{***}$ | -0.133$^{***}$ | -0.109$^{**}$ | -0.024$^{ns}$ | -0.109$^{**}$ | 0.017$^{ns}$ | -0.126$^{*}$ |
| Ile | -0.098$^{***}$ | -0.019$^{ns}$ | -0.079$^{*}$ | -0.043$^{ns}$ | 0.066$^{ns}$ | -0.109$^{*}$ | -0.101$^{**}$ | 0.130$^{***}$ | -0.231$^{***}$ |
| Leu | 0.046$^{ns}$ | 0.063$^{**}$ | -0.017$^{ns}$ | 0.027$^{ns}$ | 0.027$^{ns}$ | 0.0$^{ns}$ | -0.100$^{***}$ | -0.004$^{ns}$ | -0.096$^{**}$ |
| Lys | 0.081$^{***}$ | 0.194$^{***}$ | -0.113$^{***}$ | -0.017$^{ns}$ | 0.239$^{***}$ | -0.256$^{***}$ | 0.009$^{ns}$ | 0.098$^{***}$ | -0.089$^{**}$ |
| Met | 1.199$^{***}$ | -0.016$^{ns}$ | 1.215$^{***}$ | 1.219$^{***}$ | 0.130$^{***}$ | 1.089$^{***}$ | 1.165$^{***}$ | 0.088$^{*}$ | 1.077$^{***}$ |
| Phe | -0.006$^{ns}$ | 0.121$^{***}$ | -0.127$^{***}$ | 0.065$^{ns}$ | 0.209$^{***}$ | -0.145$^{**}$ | 0.075$^{ns}$ | 0.262$^{***}$ | -0.187$^{***}$ |
| Pro | -0.084$^{***}$ | -0.068$^{**}$ | -0.016$^{ns}$ | -0.150$^{***}$ | -0.203$^{***}$ | 0.053$^{ns}$ | -0.182$^{***}$ | -0.145$^{***}$ | -0.038$^{ns}$ |
| Ser | 0.093$^{***}$ | -0.076$^{***}$ | 0.169$^{***}$ | 0.048$^{*}$ | -0.051$^{*}$ | 0.099$^{***}$ | -0.027$^{ns}$ | -0.052$^{**}$ | 0.025$^{ns}$ |
| Thr | -0.067$^{**}$ | -0.147$^{***}$ | 0.081$^{**}$ | 0.006$^{ns}$ | -0.079$^{**}$ | 0.085$^{*}$ | -0.053$^{*}$ | -0.098$^{***}$ | 0.045$^{ns}$ |
| Trp | -0.123$^{ns}$ | 0.200$^{***}$ | -0.324$^{***}$ | 0.173$^{*}$ | 0.270$^{***}$ | -0.097$^{ns}$ | 0.429$^{***}$ | 0.434$^{***}$ | -0.006$^{ns}$ |
| Tyr | -0.298$^{***}$ | 0.046$^{ns}$ | -0.344$^{***}$ | -0.180$^{***}$ | 0.093$^{*}$ | -0.273$^{***}$ | -0.195$^{***}$ | 0.246$^{***}$ | -0.441$^{***}$ |
| Val | -0.116$^{***}$ | -0.097$^{***}$ | -0.019$^{ns}$ | -0.003$^{ns}$ | 0.008$^{ns}$ | -0.012$^{ns}$ | -0.108$^{***}$ | 0.064$^{*}$ | -0.173$^{***}$ |

[1]Residue count ranges and gene counts for each bin: Short ([101-250], 3361), Medium ([401-550], 2315), Long ([851,4976], 2146).

[2]Extent of difference between protein segments for amino acid frequencies. Subscript format "AB" specifies the segments being compared. $LOR_{AB}>0$ indicates a higher amino acid frequency at segment A relative to B. $LOR_{AB}<0$ indicates a lower frequency at segment A and $LOR_{AB}=0$ indicates similar frequencies. Asterisks indicate significance levels for Z-statistics of odds ratios. The Bonferroni sequential method was employed for multiple test corrections for each amino acid (i.e., nine tests were corrected for each amino acid). Superscripts ns, *, ** and *** indicate non-significant, $p<0.05$, $p<0.0,1$ and $p<0.001$, respectively.

**S9 Table.** Log-odds ratios (LOR) for amino acid frequency differences among protein terminal (N, C) and internal (I) for predicted *non-disordered* residues.

| AA | Protein length bin[1] | | | | | | | | |
|---|---|---|---|---|---|---|---|---|---|
| | Short | | | Medium | | | Long | | |
| | $LOR_{NI}$[2] | $LOR_{CI}$[2] | $LOR_{NC}$[2] | $LOR_{NI}$[2] | $LOR_{CI}$[2] | $LOR_{NC}$[2] | $LOR_{NI}$[2] | $LOR_{CI}$[2] | $LOR_{NC}$[2] |
| Ala | 0.216$^{***}$ | 0.057$^{**}$ | 0.159$^{***}$ | 0.105$^{***}$ | -0.036$^{ns}$ | 0.141$^{***}$ | 0.121$^{***}$ | 0.022$^{ns}$ | 0.100$^{**}$ |
| Arg | -0.198$^{***}$ | -0.042$^{ns}$ | -0.155$^{***}$ | -0.023$^{ns}$ | 0.036$^{ns}$ | -0.059$^{ns}$ | -0.063$^{*}$ | 0.096$^{***}$ | -0.159$^{***}$ |
| Asn | -0.283$^{***}$ | -0.042$^{ns}$ | -0.241$^{***}$ | -0.237$^{***}$ | -0.042$^{ns}$ | -0.195$^{***}$ | -0.093$^{**}$ | -0.032$^{ns}$ | -0.061$^{ns}$ |
| Asp | -0.313$^{***}$ | -0.100$^{***}$ | -0.214$^{***}$ | -0.358$^{***}$ | -0.106$^{***}$ | -0.253$^{***}$ | -0.167$^{***}$ | -0.099$^{***}$ | -0.068$^{ns}$ |
| Cys | 0.159$^{***}$ | 0.076$^{*}$ | 0.083$^{*}$ | 0.180$^{***}$ | -0.065$^{ns}$ | 0.245$^{***}$ | 0.038$^{ns}$ | 0.019$^{ns}$ | 0.019$^{ns}$ |
| Gln | -0.068$^{*}$ | 0.059$^{ns}$ | -0.128$^{***}$ | -0.069$^{*}$ | -0.023$^{ns}$ | -0.046$^{ns}$ | -0.126$^{***}$ | 0.028$^{ns}$ | -0.154$^{***}$ |
| Glu | -0.270$^{***}$ | 0.019$^{ns}$ | -0.289$^{***}$ | -0.285$^{***}$ | -0.060$^{*}$ | -0.225$^{***}$ | -0.258$^{***}$ | 0.005$^{ns}$ | -0.263$^{***}$ |
| Gly | -0.093$^{***}$ | -0.050$^{ns}$ | -0.043$^{ns}$ | -0.020$^{ns}$ | -0.077$^{**}$ | 0.057$^{ns}$ | 0.024$^{ns}$ | -0.030$^{ns}$ | 0.054 $^{ns}$ |
| His | -0.194$^{***}$ | -0.013$^{ns}$ | -0.181$^{***}$ | -0.231$^{***}$ | -0.086$^{*}$ | -0.145$^{**}$ | -0.110$^{**}$ | 0.033$^{ns}$ | -0.143$^{**}$ |
| Ile | 0.116$^{***}$ | -0.007$^{ns}$ | 0.123$^{***}$ | 0.146$^{***}$ | 0.144$^{***}$ | 0.003$^{ns}$ | 0.068$^{**}$ | -0.043$^{ns}$ | 0.111$^{**}$ |
| Leu | 0.329$^{***}$ | 0.049$^{**}$ | 0.280$^{***}$ | 0.339$^{***}$ | 0.043$^{**}$ | 0.296$^{***}$ | 0.209$^{***}$ | -0.030$^{ns}$ | 0.239$^{***}$ |
| Lys | -0.166$^{***}$ | 0.028$^{ns}$ | -0.194$^{***}$ | -0.115$^{***}$ | 0.153$^{***}$ | -0.268$^{***}$ | -0.114$^{***}$ | 0.063$^{*}$ | -0.177$^{***}$ |
| Met | -0.024$^{ns}$ | -0.015$^{ns}$ | -0.009$^{ns}$ | -0.153$^{***}$ | -0.075$^{ns}$ | -0.079$^{ns}$ | 0.117$^{**}$ | 0.059$^{ns}$ | 0.058$^{ns}$ |
| Phe | 0.114$^{***}$ | 0.020$^{ns}$ | 0.093$^{***}$ | 0.007$^{ns}$ | 0.086$^{***}$ | -0.079$^{*}$ | 0.056$^{ns}$ | 0.008$^{ns}$ | 0.048$^{ns}$ |
| Pro | -0.143$^{***}$ | -0.006$^{ns}$ | -0.137$^{***}$ | -0.119$^{***}$ | -0.156$^{***}$ | 0.037$^{ns}$ | -0.053$^{ns}$ | -0.004$^{ns}$ | -0.049$^{ns}$ |
| Ser | 0.096$^{***}$ | -0.027$^{ns}$ | 0.122$^{***}$ | 0.033$^{ns}$ | -0.077$^{**}$ | 0.110$^{***}$ | 0.049$^{ns}$ | 0.015$^{ns}$ | 0.034$^{ns}$ |
| Thr | 0.001$^{ns}$ | -0.029$^{ns}$ | 0.030$^{ns}$ | -0.112$^{***}$ | -0.079$^{**}$ | -0.033$^{ns}$ | -0.045$^{ns}$ | -0.030$^{ns}$ | -0.015$^{ns}$ |
| Trp | -0.089$^{ns}$ | 0.103$^{*}$ | -0.192$^{***}$ | 0.193$^{***}$ | 0.120$^{**}$ | 0.073$^{ns}$ | 0.301$^{***}$ | 0.064$^{ns}$ | 0.237$^{***}$ |
| Tyr | -0.268$^{***}$ | 0.034$^{ns}$ | -0.302$^{***}$ | -0.121$^{***}$ | 0.056$^{ns}$ | -0.176$^{***}$ | -0.108$^{**}$ | 0.022$^{ns}$ | -0.130$^{**}$ |
| Val | 0.097$^{***}$ | -0.060$^{**}$ | 0.158$^{***}$ | 0.069$^{**}$ | 0.066$^{**}$ | 0.003$^{ns}$ | 0.026$^{ns}$ | -0.069$^{**}$ | 0.094$^{**}$ |

[1]Residue count ranges and gene counts for each bin: Short ([101-250], 3361), Medium ([401-550], 2315), Long ([851,4976], 2146).

[2]Extent of difference between protein segments for amino acid frequencies. Subscript format "AB" specifies the segments being compared. $LOR_{AB}>0$ indicates a higher amino acid frequency at segment A relative to B. $LOR_{AB}<0$ indicates a lower frequency at segment A and $LOR_{AB}=0$ indicates similar frequencies. Asterisks indicate significance levels for Z-statistics of odds ratios. The Bonferroni sequential method was employed for multiple test corrections for each amino acid (i.e., nine tests were corrected for each amino acid). Superscripts ns, *, ** and *** indicate non-significant, $p$<0.05, $p$<0.0,1 and $p$<0.001, respectively.

**S10 Table.** Spearman's rank correlation coefficients ($r_s$) for the fraction of predicted disordered residues in solvent accessibility classes.

| Predicted residue class[1] | Putative solvent accessibility class | | | |
|---|---|---|---|---|
| | Exposed | | Buried | |
| | N | C | N | C |
| Disordered | 207,059 | 201,925 | 61,693 | 75,102 |
| Non-disordered | 97,828 | 57,348 | 132,300 | 164,505 |
| $r_s^a$ | -0.320$^{*}$ | -0.996$^{***}$ | 0.508$^{***}$ | -0.526$^{***}$ |
| $LOR_N^b$ | 1.513$^{***}$ | | | |
| $LOR_C^b$ | 2.043$^{***}$ | | | |

[1] Residue counts for 12472 protein sequences. Only protein sequences containing 80 residues or more were used.

[a] $r_s$ calculated between the fraction of predicted disorder residues versus the first/last 40 residues. The Bonferroni sequential method was employed for multiple test corrections across solvent accessibility classes (i.e., two tests were corrected for each terminal). Superscripts * and *** indicate $p$<0.05 and $p$<0.001, respectively.

[b]Log-odds ratio (LOR) for differences between the exposed and buried classes for the fraction of predicted disordered residues. Subscript "A" specifies a terminal (N or C). LOR>0 indicates a higher fraction for the exposed class, while LOR<0 indicates a lower fraction, and LOR=0 indicates identical fractions between classes. Asterisks indicate significance levels for Z-statistics of odds ratios.

**S11 Table.** Log-odds ratios (LORs) for differences between putative solvent accessibility classes (exposed or buried) for the fraction of predicted disordered residues.

| Predicted residue class | | Protein length bin[1] | | | | | | | | |
|---|---|---|---|---|---|---|---|---|---|---|
| | | Short | | | Medium | | | Long | | |
| | | N | I | C | N | I | C | N | I | C |
| Exposed | Disordered | 49,248 | 73,677 | 52,243 | 34,273 | 195,155 | 30,280 | 33,491 | 874,869 | 32,315 |
| | Non-disordered | 31,028 | 41,914 | 18,509 | 16,699 | 90,094 | 9,528 | 8,163 | 159,035 | 4,468 |
| Buried | Disordered | 19,492 | 47,761 | 23,677 | 9,649 | 88,140 | 11,438 | 6,628 | 237,831 | 7,795 |
| | Non-disordered | 42,825 | 121,834 | 46,475 | 24,918 | 482,001 | 33,027 | 11,713 | 828,090 | 15,042 |
| LOR[a] | | 1.249*** | 1.501*** | 1.712*** | 1.668*** | 2.472*** | 2.217*** | 1.981*** | 2.953*** | 2.636*** |

[1] Residue count ranges and gene counts for each bin: Short ([101-250], 3274), Medium ([401-550], 2171), Long ([851,4976], 1737). Residue counts are provided for terminals (N, C) and internal (I) segments.

[a] Extent of difference between the exposed and buried classes for the fraction of predicted disordered residues. LOR>0 indicates a higher fraction for the exposed class, and LOR<0 and LOR=0 indicate a lower fraction and identical fractions, respectively. Asterisks indicate significance levels for Z-statistics of odds ratios. The Bonferroni sequential method was employed for multiple test corrections across protein length bins (i.e., nine tests were corrected). Superscript *** indicates $p$<0.001.

**S12 Table.** Log-odds ratios (LORs) for differences between terminal (N, C) and internal I) segments for the fraction of predicted disordered residues in solvent accessibility classes (exposed, buried).

| Predicted residue class | Protein length bin[1] | | | | | | | | |
|---|---|---|---|---|---|---|---|---|---|
| | Short | | | Medium | | | Long | | |
| | $LOR_{NI}$[2] | $LOR_{CI}$[2] | $LOR_{NC}$[2] | $LOR_{NI}$[2] | $LOR_{CI}$[2] | $LOR_{NC}$[2] | $LOR_{NI}$[2] | $LOR_{CI}$[2] | $LOR_{NC}$[2] |
| Exposed | $0.415^{***}$ | $0.526^{***}$ | $-0.111^{***}$ | $0.816^{***}$ | $0.640^{***}$ | $0.176^{***}$ | $0.571^{***}$ | $0.505^{***}$ | $0.066^{***}$ |
| Buried | $-0.239^{***}$ | $0.004^{ns}$ | $-0.243^{***}$ | $0.101^{***}$ | $0.313^{***}$ | $-0.211^{***}$ | $-0.028^{ns}$ | $0.163^{***}$ | $-0.191^{***}$ |

[1]Residue count ranges and protein sequence counts for each bin: Short ([101, 250], 3274), Medium ([401, 550], 2171), Long ([851,4976], 1737).

[2]Extent of difference between segments for the fraction of predicted disordered residues. Subscript format "AB" specifies the segments being compared. $LOR_{AB}>0$ indicates a higher fraction of predicted disordered residues at segment A relative to B. $LOR_{AB}<0$ indicates a lower fraction at segment A and $LOR_{AB}=0$ indicates identical fractions. Asterisks indicate significance levels for Z-statistics of odds ratios. The Bonferroni sequential method was employed for multiple test corrections among protein length bins (i.e., nine tests were corrected for each solvent accessibility class). Superscript *** and ns indicate $p<0.001$ and non-significance, respectively.

**S13 Table.** Log-odds ratios (LORs) for differences between putative functional classes for the fraction of predicted disordered residues.

| Focal class[1] | Comparison class | Short: $LOR_N$[3] | Short: $LOR_I$[3] | Short: $LOR_C$[3] | Medium: $LOR_N$[3] | Medium: $LOR_I$[3] | Medium: $LOR_C$[3] | Long: $LOR_N$[3] | Long: $LOR_I$[3] | Long: $LOR_C$[3] |
|---|---|---|---|---|---|---|---|---|---|---|
| Protein-binding | Protein-only binding | 1.850 | 1.436 | 1.553 | 1.353 | 1.253 | 1.554 | 1.364 | 1.302 | 1.618 |
| | RNA-binding | 0.576 | 0.788 | 0.642 | 0.704 | 0.909 | 0.601 | 0.515 | 0.760 | 0.689 |
| | DNA-binding | 0.323 | 0.478 | 0.378 | 0.468 | 0.628 | 0.299 | 0.311 | 0.525 | 0.251 |
| | Linker | 0.562 | 0.300 | 0.715 | 0.702 | 0.419 | 0.667 | 0.473 | 0.351 | 0.425 |
| | Non-protein binding | 1.425 | 1.332 | 1.556 | 1.586 | 1.754 | 1.536 | 1.360 | 1.688 | 1.650 |
| | Non-binding-non-linker | 3.510 | 3.185 | 3.575 | 3.547 | 3.606 | 3.780 | 3.474 | 3.420 | 3.804 |
| Protein-only binding | RNA-only binding | 2.307 | 3.615 | 3.191 | 3.201 | 3.909 | 3.527 | 3.679 | 5.033 | 3.704 |
| | DNA-only binding | 2.330 | 2.642 | 2.458 | 2.721 | 3.449 | 2.701 | 2.751 | 2.565 | 2.414 |
| | Linker-only | 2.248 | 2.343 | 2.941 | 3.074 | 3.160 | 2.960 | 3.628 | 3.507 | 3.149 |
| | Non-binding-non-linker | 1.660 | 1.749 | 2.022 | 2.194 | 2.353 | 2.226 | 2.110 | 2.118 | 2.186 |
| RNA-binding | RNA-only binding | 3.581 | 4.262 | 4.102 | 3.850 | 4.254 | 4.480 | 4.528 | 5.575 | 4.633 |
| | DNA-binding | -0.253 | -0.311 | -0.264 | -0.235 | -0.281 | -0.302 | -0.204 | -0.235 | -0.438 |
| | Linker | -0.014[ns] | -0.488 | 0.073 | -0.002[ns] | -0.490 | 0.066 | -0.04[ns] | -0.408 | -0.264 |
| | Non-RNA binding | -0.098 | -0.551 | -0.171 | -0.289 | -0.617 | -0.128 | -0.055 | -0.402 | -0.273 |
| | Non-binding-non-linker | 2.934 | 2.397 | 2.933 | 2.843 | 2.697 | 3.179 | 2.958 | 2.660 | 3.115 |
| RNA-only binding | DNA-only binding | 0.023[ns] | -0.973 | -0.734 | -0.480 | -0.460 | -0.826 | -0.928 | -2.468 | -1.290 |
| | Linker-only | -0.059[ns] | -1.272 | -0.250[ns] | -0.128[ns] | -0.749 | -0.566 | -0.051[ns] | -1.526 | -0.555[ns] |
| | Non-binding-non-linker | -0.647 | -1.866 | -1.169 | -1.007 | -1.556 | -1.301 | -1.569 | -2.914 | -1.518 |
| DNA-binding | DNA-only binding | 3.857 | 3.600 | 3.632 | 3.606 | 4.074 | 3.956 | 3.804 | 3.342 | 3.781 |
| | Linker | 0.239 | -0.178 | 0.337 | 0.233 | -0.209 | 0.368 | 0.162 | -0.174 | 0.174 |
| | Non-DNA binding | 0.442 | 0.004[ns] | 0.378 | 0.171 | -0.137 | 0.531 | 0.385 | 0.031 | 0.692 |
| | Non-binding-non-linker | 3.187 | 2.707 | 3.197 | 3.079 | 2.978 | 3.481 | 3.163 | 2.895 | 3.553 |
| | Linker-only | -0.082[ns] | -0.299 | 0.484 | 0.353 | -0.289 | 0.260[ns] | 0.877 | 0.942 | 0.736 |
| DNA-only binding | Non-binding-non-linker | -0.670 | -0.893 | -0.435 | -0.527 | -1.096 | -0.474 | -0.641 | -0.447 | -0.228[ns] |
| Linker | Linker-only | 3.536 | 3.479 | 3.779 | 3.725 | 3.994 | 3.848 | 4.518 | 4.457 | 4.343 |
| | Non-linker | -0.072 | 0.393 | -0.305 | -0.285 | 0.297 | -0.250 | 0.029[ns] | 0.419 | 0.254 |
| | Non-binding-non-linker | 2.948 | 2.885 | 2.860 | 2.845 | 3.187 | 3.113 | 3.001 | 3.069 | 3.380 |
| Linker-only | Non-binding-non-linker | -0.588 | -0.594 | -0.919 | -0.880 | -0.807 | -0.734 | -1.518 | -1.389 | -0.963 |

Protein length bin[2]: Short, Medium, Long.

[1]Functional classes based on flDPnn program. Note that only data for putative solvent-exposed residues are compared.

[2]Residue count ranges and protein sequence counts for each bin: Short ([101-250], 1947), Medium ([401-550], 1851), long (851,4976), 1558).

[3]Extent of difference between focal and comparison classes for the fraction of predicted disordered residues at terminal (N, C) and internal (I) segments. LOR>0 indicates a higher fraction for the focal class. LOR<0 and LOR=0 indicate a lower fraction and identical fractions,

respectively. Z-statistics of odds ratios were tested for significance. The Bonferroni sequential method was employed for multiple test corrections among bins, separately for each comparison (i.e., nine tests were corrected per comparison). Superscript "ns" indicates non-significance. $p<0.05$ for all other LOR values.

**S14 Table.** Log-odds ratios (LORs) for differences among terminal (N, C) and internal (I) segments for the fraction of predicted disordered residues at putative solvent-exposed functional classes.

| Predicted class | Protein length bin[1] | | | | | | | | |
|---|---|---|---|---|---|---|---|---|---|
| | Short | | | Medium | | | Long | | |
| | $LOR_{NI}$[2] | $LOR_{CI}$[2] | $LOR_{NC}$[2] | $LOR_{NI}$[2] | $LOR_{CI}$[2] | $LOR_{NC}$[2] | $LOR_{NI}$[2] | $LOR_{CI}$[2] | $LOR_{NC}$[2] |
| Protein-binding | 0.827 | 0.793 | 0.034 | 0.757 | 0.619 | 0.138 | 0.589 | 0.531 | 0.058 |
| Protein-only binding | 0.413 | 0.676 | -0.263 | 0.657 | 0.318 | 0.339 | 0.527 | 0.214 | 0.313 |
| RNA-binding | 1.040 | 0.939 | 0.100 | 0.962 | 0.926 | 0.035$^{ns}$ | 0.833 | 0.601 | 0.232 |
| RNA-only binding | 1.721 | 1.100 | 0.621 | 1.365 | 0.700 | 0.665 | 1.880 | 1.543 | 0.338$^{ns}$ |
| DNA-binding | 0.981 | 0.893 | 0.089 | 0.916 | 0.948 | -0.031$^{ns}$ | 0.803 | 0.805 | -0.002$^{ns}$ |
| DNA-only binding | 0.725 | 0.860 | -0.136$^{ns}$ | 1.385 | 1.067 | 0.319 | 0.340 | 0.365 | -0.025$^{ns}$ |
| Linker | 0.565 | 0.378 | 0.187 | 0.474 | 0.371 | 0.103 | 0.467 | 0.457 | 0.010$^{ns}$ |
| Linker-only | 0.508 | 0.077$^{ns}$ | 0.430 | 0.743 | 0.517 | 0.226$^{ns}$ | 0.406 | 0.572 | -0.166$^{ns}$ |
| Non-binding-non-linker | 0.502 | 0.403 | 0.099$^{ns}$ | 0.816 | 0.445 | 0.371 | 0.535 | 0.146$^{ns}$ | 0.389 |
| Non-protein binding | 0.734 | 0.569 | 0.165 | 0.925 | 0.837 | 0.088 | 0.918 | 0.569 | 0.349 |
| Non-RNA binding | 0.586 | 0.559 | 0.027$^{ns}$ | 0.633 | 0.437 | 0.196 | 0.487 | 0.472 | 0.014$^{ns}$ |
| Non-DNA binding | 0.543 | 0.518 | 0.025$^{ns}$ | 0.609 | 0.280 | 0.328 | 0.449 | 0.144 | 0.305 |
| Non-linker | 1.029 | 1.076 | -0.046 | 1.056 | 0.918 | 0.138 | 0.857 | 0.622 | 0.235 |

[1]Residue count ranges and protein sequence counts for each bin: Short ([101-250], 1947), Medium ([401-550], 1851), long ([851,4976], 1558).
[2]Extent of difference for the fraction of predicted disordered residues between a pair of protein segments. Subscript format "AB" specifies the segments being compared. $LOR_{AB}>0$ indicates a higher fraction at segment A relative to B. $LOR_{AB}<0$ indicates a lower frequency at segment A and $LOR_{AB}=0$ indicates a similar fraction. Asterisks indicate significance levels for Z-statistics of odds ratios. The Bonferroni sequential method was employed for multiple test corrections for each amino acid (i.e., nine tests were corrected for each amino acid). Superscript "ns" indicates non-significance. $p<0.05$ for all other LOR values.

**S15 Table**. Spearman's rank correlation coefficients ($r_s$) for the fraction of predicted disordered residues at terminals for protein connectivity classes.

| Predicted residue class[1] | Protein connectivity class | | | |
|---|---|---|---|---|
| | Hub | | Non-hub | |
| | N | C | N | C |
| Disordered | 120,501 | 115,807 | 91,933 | 94,063 |
| Non-disordered | 74,259 | 78,953 | 82,867 | 80,737 |
| $r_s$[a] | -0.886*** | -0.998*** | -0.188[ns] | -0.999*** |
| $LOR_N$[b] | 0.380*** | | | |
| $LOR_C$[b] | 0.230*** | | | |

[1] Counts from 4869 and 4370 protein sequences for "hub" and "non-hub" classes, respectively. Only protein sequences containing 80 residues or more were used.

[a] Correlation coefficient for the fraction of disordered residues at the first/last 40 residues of proteins. The Bonferroni sequential method was employed for multiple test corrections for each terminal (i.e., two tests were corrected for each terminal). Superscript *** indicates significance at $p<0.001$, and "ns" indicates non-significance.

[b] Log-odds ratio (LOR) for differences between hub and non-hub classes for the fraction of predicted disordered residues. Subscripts specify terminals and LOR>0 indicates a higher fraction for the hub class than the non-hub class. LOR<0 indicates a lower fraction, and LOR=0 would indicate identical fractions. Significance levels for Z-statistics of odds ratios are shown. Multiple test correction was done as described above, and superscript *** indicates $p<0.001$.

**S16 Table.** Log-odds ratios (LORs) for differences between protein connectivity classes for the fraction of predicted disordered residues.

| Predicted residue class | | Protein length bin[1] | | | | | | | | |
|---|---|---|---|---|---|---|---|---|---|---|
| | | Short | | | Medium | | | Long | | |
| | | N | I | C | N | I | C | N | I | C |
| Hub | Disordered | 24,099 | 37,473 | 24,437 | 20,141 | 134,402 | 18,222 | 24,568 | 665,026 | 23,453 |
| | Non-disordered | 17,607 | 45,939 | 16,785 | 11,478 | 181,788 | 12,985 | 20,248 | 903,534 | 21,116 |
| Non-hub | Disordered | 22,754 | 41,155 | 25,495 | 15,200 | 95,159 | 14,385 | 14,415 | 365,061 | 13,539 |
| | Non-disordered | 23,370 | 51,093 | 20,085 | 16,431 | 221,151 | 16,780 | 11,460 | 540,564 | 12,182 |
| LOR[a] | | $0.341^{***}$ | $0.013^{ns}$ | $0.137^{***}$ | $0.640^{***}$ | $0.541^{***}$ | $0.493^{***}$ | $-0.036^{ns}$ | $0.086^{***}$ | $-0.001^{ns}$ |

[1]Residue count ranges and protein sequence counts (hub and non-hub) for each bin: Short ([101-250],943,1043), Medium ([401-550], 805,800), long ([851,4976], 1132,685). Residue counts are shown separately for terminal (N, C) and internal (I) segments.

[a]Extent of difference in the fraction of disordered residues between hub and non-hub classes. LOR>0 indicates a higher fraction for hub than non-hub. LOR<0 or LOR=0 indicate lower and identical fractions, respectively. Superscripts indicate significance levels for Z-statistics of odds ratios. The Bonferroni sequential method was employed for multiple test corrections across protein length bins (i.e., nine tests were corrected for each protein connectivity class). Superscripts *** and ns indicate $p<0.001$ and non-significance, respectively.

**S17 Table.** Log-odds ratios (LORs) for differences among protein segments for the fraction of predicted disordered residues in protein connectivity classes.

| Protein connectivity class | Protein length bin[1] | | | | | | | | |
|---|---|---|---|---|---|---|---|---|---|
| | Short[1] | | | Medium[1] | | | Long[1] | | |
| | $LOR_{NI}$[2] | $LOR_{CI}$[2] | $LOR_{NC}$[2] | $LOR_{NI}$[2] | $LOR_{CI}$[2] | $LOR_{NC}$[2] | $LOR_{NI}$[2] | $LOR_{CI}$[2] | $LOR_{NC}$[2] |
| Hub | 0.518*** | 0.579*** | -0.062*** | 0.864*** | 0.641*** | 0.223*** | 0.500*** | 0.411*** | 0.088*** |
| Non-hub | 0.190*** | 0.455*** | -0.265*** | 0.765*** | 0.689*** | 0.076*** | 0.622*** | 0.498*** | 0.124*** |

[1]Residue count ranges and protein sequences counts (hub and non-hub) for each bin: Short ([101-250],943,1043), Medium ([401-550], 805,800), long ([851,4976], 1132,685).
[2]Extent of difference among protein segment pairings for the fraction of predicted disordered residues. Subscript format "AB" specifies the segments being compared. $LOR_{AB}>0$ indicates a higher fraction for segment A than B. $LOR_{AB}<0$ indicates a lower fraction at segment A and $LOR_{AB}=0$ indicates similar fractions. Asterisks indicate significance levels for Z-statistics of odds ratios. The Bonferroni sequential method was employed for multiple test corrections across protein length bins (i.e., nine tests were corrected for each protein connectivity class). Superscript *** indicates $p<0.001$.

**S18 Table**. Top 5 segment-specific GO term (ten genes or above) enrichments based on the fraction of predicted disordered residue.

| Segment ID/type[a] | Term ID | Term name[b] | # genes | Fraction of predicted disordered residues[c] |
|---|---|---|---|---|
| 1 (N terminal) | 0035059 | C: RCAF complex | 47 | 0.964*** |
| | 0106222 | F: lncRNA binding | 11 | 0.762*** |
| | 0000900 | F: mRNA regulatory element binding translation repressor activity | 12 | 0.647** |
| | 0030532 | C: small nuclear ribonucleoprotein complex | 10 | 0.623*** |
| | 0048137 | P: spermatocyte division | 11 | 0.599** |
| 2 (Internal) | 0010507 | P: negative regulation of autophagy | 13 | 0.474** |
| | 0030426 | C: growth cone | 11 | 0.451** |
| | 0046329 | P: negative regulation of JNK cascade | 25 | 0.438* |
| | 0007314 | P: oocyte anterior/posterior axis specification | 16 | 0.430** |
| | 0031252 | C: cell leading edge | 11 | 0.424*** |
| 2 (Internal) | 0032456 | P: endocytic recycling | 21 | 0.413*** |
| | 0030713 | P: follicle cell of egg chamber stalk formation | 11 | 0.391** |
| | 0048515 | P: spermatid differentiation | 13 | 0.354* |
| | 0006898 | P: receptor-mediated endocytosis | 13 | 0.306*** |
| | 0032039 | C: integrator complex | 15 | 0.300** |
| 4 (C terminal) | 0008188 | F: neuropeptide receptor activity | 12 | 0.794*** |
| | 0003924 | F: GTPase activity | 24 | 0.712*** |
| | 0007189 | P: adenylate cyclase-activating G protein-coupled receptor signaling pathway | 12 | 0.710*** |
| | 0008306 | P: associative learning | 15 | 0.708*** |
| | 0009649 | P: entrainment of circadian clock | 13 | 0.667*** |

[a]Data for genes belonging to a given GO term were pooled. Protein sequences were split into four parts such that each part is proportional to the protein length.

[b]Prefixes C, F, P specify “Cellular component”, “Molecular Function” and “Biological Process”, respectively.

[c]Superscripts *, ** and *** indicate $p<0.05$, $p<0.01$, $p<0.001$, respectively, based on hypergeometric tests.

**S19 Table.** Log-odds ratios (LOR) for differences between Actual and Shuffled data for the fraction of predicted disordered residues for protein segments.

| Data and residue class | | Protein length bins[1] | | | | | | | | |
|---|---|---|---|---|---|---|---|---|---|---|
| | | Short | | | Medium | | | Long | | |
| | | N | I | C | N | I | C | N | I | C |
| Actual | Disordered | 30,106 | 35,574 | 27,823 | 17,738 | 90,024 | 15,995 | 9,020 | 175,130 | 7,840 |
| | Non-disordered | 59,027 | 14,2692 | 60,299 | 60,793 | 695,286 | 61,380 | 66,337 | 246,2365 | 67,181 |
| Shuffled | Disordered | 22,446 | 29,327 | 20,187 | 7,162 | 54,405 | 9,276 | 3,006 | 52,506 | 3,573 |
| | Non-disordered | 66,687 | 148,939 | 67,935 | 71,369 | 730,905 | 68,099 | 72,351 | 2,584,989 | 71,448 |
| | $LOR_N$[a] | 0.294*** | | | 0.907*** | | | 1.099*** | | |
| | $LOR_I$[a] | 0.193*** | | | 0.504*** | | | 1.206*** | | |
| | $LOR_C$[a] | 0.321*** | | | 0.545*** | | | 0.786*** | | |

[1]Residue count ranges and gene counts for each bin: Short ([101-250], 2005), Medium ([401-550], 1981), Long ([851,4976], 1951). Residue counts for each segment are shown. Note that counts for a given segment are pooled among protein sequences.

[a]Extent of differences between Actual and Shuffled data for the fraction of predicted disordered residues for each segment. LOR>0 indicates a higher fraction for the Actual data, and LOR<0 and LOR=0 indicate lower or identical fractions, respectively. Asterisks indicate significance levels for Z-statistics of odds ratios. The Bonferroni sequential method was employed for multiple test corrections among bins (i.e., nine tests were corrected for each segment). Superscript *** indicates $p<0.001$.

**S20 Table.** Z-statistics for comparisons between Actual and Shuffled data for differences in the distribution of predicted disordered residues.

| Dataset | Protein length bin[1] | | | | | | | | |
|---|---|---|---|---|---|---|---|---|---|
| | Short[1] | | | Medium[1] | | | Long[1] | | |
| | $LOR_{NI}$[2] | $LOR_{CI}$[2] | $LOR_{NC}$[2] | $LOR_{NI}$[2] | $LOR_{CI}$[2] | $LOR_{NC}$[2] | $LOR_{NI}$[2] | $LOR_{CI}$[2] | $LOR_{NC}$[2] |
| Actual | $0.716^{***}$ | $0.616^{***}$ | $0.100^{***}$ | $0.812^{***}$ | $0.699^{***}$ | $0.113^{***}$ | $0.648^{***}$ | $0.495^{***}$ | $0.153^{***}$ |
| Shuffled | $0.536^{***}$ | $0.412^{***}$ | $0.125^{***}$ | $0.299^{***}$ | $0.604^{***}$ | $-0.306^{***}$ | $0.716^{***}$ | $0.901^{***}$ | $-0.185^{***}$ |
| Z[3] | $13.187^{***}$ | $14.703^{***}$ | $-1.624^{ns}$ | $31.936^{***}$ | $6.225^{***}$ | $20.239^{***}$ | $-3.029^{**}$ | $-18.882^{***}$ | $11.219^{***}$ |

[1]Residue count ranges and gene counts for each bin: Short ([101-250], 2005), Medium ([401-550], 1981), Long ([851,4976], 1951).

[2]Extent of difference between segment pairs for the fraction of predicted disordered residues. Subscript format "AB" specifies the segments being compared, and $LOR_{AB}>0$ indicates a higher fraction for segment A than B. $LOR_{AB}<0$ indicates a lower fraction at segment A and $LOR_{AB}=0$ indicates identical fractions. Asterisks indicate significance levels for Z-statistics of odds ratios. The Bonferroni sequential method was employed for multiple test corrections (i.e., nine tests were corrected for each dataset). Superscript *** indicates $p<0.001$.

[3]Wald test statistic for a comparison of LORs between Actual and Shuffled data. Superscripts ns, ** and *** indicate non-significance, $p<0.01$ and $p<0.001$, respectively.

**S21 Table.** Log-odds ratios (LOR) for differences among protein segments for amino acid frequencies in *Actual* data.

| AA | Protein length bin[1] | | | | | | | | |
|---|---|---|---|---|---|---|---|---|---|
| | Short | | | Medium | | | Long | | |
| | $LOR_{NI}$[2] | $LOR_{CI}$[2] | $LOR_{NC}$[2] | $LOR_{NI}$[2] | $LOR_{CI}$[2] | $LOR_{NC}$[2] | $LOR_{NI}$[2] | $LOR_{CI}$[2] | $LOR_{NC}$[2] |
| Ala | 0.098$^{**}$ | 0.053 | 0.044 | 0.036 | 0.011 | 0.024 | 0.203$^{***}$ | 0.08 | 0.124 |
| Arg | -0.078 | 0.049 | -0.126 | -0.013 | -0.069 | 0.057 | 0.527$^{***}$ | -0.014 | 0.542$^{***}$ |
| Asn | -0.007 | 0.021 | -0.028 | 0.018 | 0.034 | -0.015 | -0.052 | 0.11 | -0.162 |
| Asp | -0.106$^{**}$ | 0.002 | -0.109$^{**}$ | -0.072 | 0.073 | -0.145$^{**}$ | -0.144$^{**}$ | 0.144$^{**}$ | -0.287$^{***}$ |
| Cys | 0.093 | 0.168 | -0.076 | 0.048 | 0.203 | -0.155 | 0.155 | -0.274 | 0.429 |
| Gln | -0.143 | 0.019 | -0.162$^{***}$ | -0.143 | -0.144 | 0.001 | -0.037 | -0.460$^{***}$ | 0.423$^{***}$ |
| Glu | -0.214$^{***}$ | 0.035 | -0.248$^{***}$ | -0.081$^{*}$ | 0.139$^{***}$ | -0.220$^{***}$ | -0.156$^{**}$ | -0.072 | -0.084 |
| Gly | -0.135$^{***}$ | -0.136$^{***}$ | 0 | -0.170$^{***}$ | -0.181$^{***}$ | 0.011 | 0.002 | 0.150$^{**}$ | -0.148$^{*}$ |
| His | -0.175$^{**}$ | -0.045 | -0.13 | -0.125 | -0.200$^{**}$ | 0.075 | -0.214 | 0.053 | -0.267 |
| Ile | -0.151$^{**}$ | -0.165$^{**}$ | 0.014 | -0.068 | 0.091 | -0.159 | -0.275$^{**}$ | 0.09 | -0.365$^{**}$ |
| Leu | 0.085 | -0.007 | 0.092 | 0.079 | -0.046 | 0.124 | 0.085 | -0.078 | 0.163 |
| Lys | 0.129$^{***}$ | 0.248$^{***}$ | -0.119$^{***}$ | 0.140$^{***}$ | 0.434$^{***}$ | -0.294$^{***}$ | 0.465$^{***}$ | 0.034 | 0.431$^{***}$ |
| Met | 1.233$^{***}$ | 0.154$^{*}$ | 1.078$^{***}$ | 1.198$^{***}$ | 0.212$^{**}$ | 0.986$^{***}$ | 1.359$^{***}$ | 0.054 | 1.305$^{***}$ |
| Phe | -0.038 | -0.04 | 0.002 | 0.199$^{*}$ | 0.240$^{**}$ | -0.041 | -0.025 | 0.315$^{**}$ | -0.341$^{*}$ |
| Pro | -0.115 | -0.023 | -0.092$^{*}$ | -0.12 | -0.313$^{***}$ | 0.193$^{***}$ | -0.137$^{**}$ | -0.089 | -0.048 |
| Ser | 0.159$^{***}$ | -0.017 | 0.176$^{***}$ | -0.026 | -0.080$^{*}$ | 0.053 | -0.148$^{***}$ | -0.006 | -0.142$^{**}$ |
| Thr | -0.032 | -0.163$^{***}$ | 0.131$^{**}$ | 0.006 | -0.003 | 0.008 | -0.335$^{***}$ | -0.103 | -0.231 |
| Trp | -0.284 | 0.086 | -0.370$^{*}$ | 0.352$^{*}$ | 0.247 | 0.105 | 0.339 | 0.439 | -0.1 |
| Tyr | -0.442$^{***}$ | -0.222 | -0.220$^{**}$ | -0.043 | 0.106 | -0.149 | -0.28 | 0.272$^{*}$ | -0.552 |
| Val | -0.048 | -0.113$^{*}$ | 0.065 | -0.003 | 0.02 | -0.023 | -0.240$^{**}$ | -0.007 | -0.233 |

[1]Residue count ranges and gene counts for each bin: Short ([101-250], 2005), Medium ([401-550], 1981), Long ([851,4976], 1951).

[2]Extent of difference between a pair of segments for amino acid frequencies. Subscript format "AB" specifies the segments being compared. $LOR_{AB}>0$ indicates a higher amino acid frequency at segment A relative to B. $LOR_{AB}<0$ indicates a lower frequency at segment A and $LOR_{AB}=0$ indicates similar frequencies. Asterisks indicate significance levels for Z-statistics of odds ratios. The Bonferroni sequential method was employed for multiple test corrections for each amino acid (i.e., nine tests were corrected for each amino acid). Superscripts ns, *, **, and *** indicate non-significance, $p<0.05$, $p<0.01$, $p<0.001$, respectively.

**S22 Table.** Log-odds ratios (LOR) for differences among protein segments for amino acid frequencies in *Shuffled* data.

| AA | Protein length bin[1] | | | | | | | | |
|---|---|---|---|---|---|---|---|---|---|
| | Short | | | Medium | | | Long | | |
| | $LOR_{NI}$[2] | $LOR_{CI}$[2] | $LOR_{NC}$[2] | $LOR_{NI}$[2] | $LOR_{CI}$[2] | $LOR_{NC}$[2] | $LOR_{NI}$[2] | $LOR_{CI}$[2] | $LOR_{NC}$[2] |
| Ala | -0.023 | 0.035 | -0.057 | -0.015 | 0.006 | -0.022 | -0.002 | -0.002 | -0.001 |
| Arg | 0.084 | 0.081 | 0.003 | -0.031 | -0.104 | 0.073 | 0.484$^{***}$ | -0.175 | 0.659$^{***}$ |
| Asn | 0.079 | 0.129$^{*}$ | -0.05 | -0.03 | 0.077 | -0.107 | -0.510$^{***}$ | 0.358$^{***}$ | -0.868$^{***}$ |
| Asp | 0 | 0.049 | -0.049 | -0.102 | 0.025 | -0.127 | -0.248$^{*}$ | 0.118 | -0.367$^{**}$ |
| Cys | 0.045 | 0.02 | 0.025 | 0.259 | 0.182 | 0.077 | 0.089 | -0.453 | 0.542 |
| Gln | -0.141$^{**}$ | -0.053 | -0.088 | -0.031 | -0.123$^{*}$ | 0.092 | 0.13 | -0.451$^{***}$ | 0.581$^{***}$ |
| Glu | -0.129 | -0.04 | -0.089 | -0.103 | -0.057 | -0.046 | -0.197 | 0.08 | -0.277$^{*}$ |
| Gly | -0.164$^{***}$ | -0.104$^{*}$ | -0.06 | -0.019 | -0.110$^{*}$ | 0.091 | -0.202$^{*}$ | 0.237 | -0.439$^{***}$ |
| His | -0.171$^{*}$ | -0.175$^{*}$ | 0.004 | -0.018 | -0.258$^{*}$ | 0.24 | 0.291 | 0.297 | -0.007 |
| Ile | 0.153 | 0.192$^{*}$ | -0.039 | -0.059 | 0.260$^{*}$ | -0.319 | -0.008 | -0.008 | 0 |
| Leu | 0.099 | 0.178 | -0.079 | 0.181$^{*}$ | 0.255$^{***}$ | -0.074 | 0.124 | 0.081 | 0.043 |
| Lys | -0.160$^{***}$ | -0.073 | -0.087$^{*}$ | 0.051 | -0.095 | 0.146 | 0.376$^{***}$ | -0.322$^{**}$ | 0.697$^{***}$ |
| Met | 1.369$^{***}$ | 0.014 | 1.355$^{***}$ | 1.446$^{***}$ | 0.274$^{*}$ | 1.173$^{***}$ | 1.715$^{***}$ | -0.182 | 1.897$^{***}$ |
| Phe | 0.161 | 0.322$^{***}$ | -0.161 | 0.164 | 0.346 | -0.182 | 0.099 | 0.21 | -0.111 |
| Pro | -0.120$^{**}$ | -0.097$^{*}$ | -0.023 | -0.083 | -0.117$^{*}$ | 0.034 | -0.056 | 0.064 | -0.12 |
| Ser | -0.083 | -0.042 | -0.041 | -0.087 | -0.044 | -0.042 | -0.012 | -0.145$^{*}$ | 0.133 |
| Thr | -0.071 | -0.034 | -0.036 | -0.119 | 0.072 | -0.191$^{*}$ | -0.233$^{**}$ | -0.06 | -0.173 |
| Trp | 0.297 | 0.355 | -0.058 | 0.454 | 0.741$^{***}$ | -0.287 | 0.371 | -0.044 | 0.415 |
| Tyr | -0.002 | -0.034 | 0.032 | 0.061 | 0.179 | -0.118 | 0.042 | 0.201 | -0.159 |
| Val | 0.12 | 0.008 | 0.112 | -0.039 | 0.286$^{***}$ | -0.325$^{**}$ | -0.074 | 0.036 | -0.11 |

[1]Residue count ranges and gene counts for each bin: Short ([101-250], 2005), Medium ([401-550], 1981), Long ([851,4976], 1951).

[2]Extent of difference between a pair of segments for amino acid frequencies. Subscript format "AB" specifies the segments being compared. $LOR_{AB}>0$ indicates a higher amino acid frequency at segment A relative to B. $LOR_{AB}<0$ indicates a lower frequency at segment A and $LOR_{AB}=0$ indicates similar frequencies. Asterisks indicate significance levels for Z-statistics of odds ratios. The Bonferroni sequential method was employed for multiple test corrections for each amino acid (i.e., nine tests were corrected for each amino acid). Superscripts ns, *, **, and *** indicate non-significance, $p<0.05$, $p<0.01$, and $p<0.001$, respectively.

**S23 Table.** Log-odds ratios (LOR) for differences between Actual and Shuffled data for the fraction of predicted disordered residues among functional classes.

| Predicted residue class | Protein length bins[1] | | | | | | | | |
|---|---|---|---|---|---|---|---|---|---|
| | Short | | | Medium | | | Long | | |
| | $LOR_N^a$ | $LOR_I^a$ | $LOR_C^a$ | $LOR_N^a$ | $LOR_I^a$ | $LOR_C^a$ | $LOR_N^a$ | $LOR_I^a$ | $LOR_C^a$ |
| Protein-binding | 0.271*** | 0.116*** | 0.296*** | 0.859*** | 0.472*** | 0.471*** | 0.989*** | 1.215*** | 0.756*** |
| RNA-binding | 0.248*** | 0.134*** | 0.279*** | 0.780*** | 0.338*** | 0.521*** | 0.992*** | 1.046*** | 0.649*** |
| DNA-binding | 0.250*** | 0.175*** | 0.257*** | 0.737*** | 0.308*** | 0.496*** | 1.024*** | 1.079*** | 0.750*** |
| Linker | 0.280*** | 0.202*** | 0.249*** | 0.623*** | 0.374*** | 0.412*** | 1.054*** | 1.112*** | 0.794*** |

[1]Residue count ranges and gene counts for each bin: Short ([101-250], 2005), Medium ([401-550], 1981), Long ([851,4976], 1951).

[a]Extent of difference between Actual and Shuffled data for the fraction of predicted disordered residues for a given functional class. Subscripts specify protein segments and $LOR_A>0$ indicates a higher fraction in the Actual data while $LOR_A<0$ and $LOR_A=0$ indicate lower, and equal fractions, respectively. Asterisks indicate significance levels for Z-statistics of odds ratios. The Bonferroni sequential method was employed for multiple test corrections within each functional class and nine tests were corrected (among protein length bins). Superscript *** indicates $p<0.001$.

**S24 Table.** Z-statistics for comparisons between Actual and Shuffled data for differences in the distribution of predicted disordered residues among functional classes.

| Predicted residue class | | Protein length bins[1] | | | | | | | | |
|---|---|---|---|---|---|---|---|---|---|---|
| | | Short | | | Medium | | | Long | | |
| | | $LOR_{NI}$[a] | $LOR_{CI}$[a] | $LOR_{NC}$[a] | $LOR_{NI}$[a] | $LOR_{CI}$[a] | $LOR_{NC}$[a] | $LOR_{NI}$[a] | $LOR_{CI}$[a] | $LOR_{NC}$[a] |
| Protein-binding | Actual | $0.687^{***}$ | $0.622^{***}$ | $0.065^{***}$ | $0.759^{***}$ | $0.641^{***}$ | $0.118^{***}$ | $0.567^{***}$ | $0.466^{***}$ | $0.101^{***}$ |
| | Shuffled | $0.469^{***}$ | $0.378^{***}$ | $0.092^{***}$ | $0.283^{***}$ | $0.613^{***}$ | $-0.330^{***}$ | $0.775^{***}$ | $0.928^{***}$ | $-0.153^{***}$ |
| | $Z^b$ | $14.375^{***}$ | $15.902^{***}$ | $-1.621^{ns}$ | $26.992^{***}$ | $1.700^{ns}$ | $19.751^{***}$ | $-8.513^{***}$ | $-19.496^{***}$ | $7.682^{***}$ |
| RNA-binding | Actual | $0.739^{***}$ | $0.664^{***}$ | $0.075^{***}$ | $0.925^{***}$ | $0.914^{***}$ | $0.011^{ns}$ | $0.802^{***}$ | $0.561^{***}$ | $0.241^{***}$ |
| | Shuffled | $0.594^{***}$ | $0.487^{***}$ | $0.107^{***}$ | $0.433^{***}$ | $0.700^{***}$ | $-0.267^{***}$ | $0.837^{***}$ | $0.960^{***}$ | $-0.123^{***}$ |
| | $Z^b$ | $7.902^{***}$ | $9.485^{***}$ | $-1.672^{ns}$ | $21.966^{***}$ | $10.090^{***}$ | $9.815^{***}$ | $-1.146^{ns}$ | $-13.128^{***}$ | $8.718^{***}$ |
| DNA-binding | Actual | $0.745^{***}$ | $0.657^{***}$ | $0.088^{***}$ | $0.883^{***}$ | $0.919^{***}$ | $-0.036^{*}$ | $0.744^{***}$ | $0.736^{***}$ | $0.008^{ns}$ |
| | Shuffled | $0.635^{***}$ | $0.543^{***}$ | $0.092^{***}$ | $0.394^{***}$ | $0.691^{***}$ | $-0.297^{***}$ | $0.777^{***}$ | $1.058^{***}$ | $-0.281^{***}$ |
| | $Z^b$ | $6.498^{***}$ | $6.614^{***}$ | $-0.226^{ns}$ | $24.285^{***}$ | $12.094^{***}$ | $10.237^{***}$ | $-1.151^{ns}$ | $-12.233^{***}$ | $7.684^{***}$ |
| Linker | Actual | $0.438^{***}$ | $0.268^{***}$ | $0.170^{***}$ | $0.465^{***}$ | $0.416^{***}$ | $0.048^{**}$ | $0.453^{***}$ | $0.450^{***}$ | $0.003^{ns}$ |
| | Shuffled | $0.333^{***}$ | $0.208^{***}$ | $0.125^{***}$ | $0.184^{***}$ | $0.366^{***}$ | $-0.183^{***}$ | $0.497^{***}$ | $0.767^{***}$ | $-0.269^{***}$ |
| | $Z^b$ | $6.117^{***}$ | $3.418^{**}$ | $2.283^{*}$ | $13.368^{***}$ | $2.453^{*}$ | $8.387^{***}$ | $-1.459^{ns}$ | $-11.300^{***}$ | $6.749^{***}$ |

[1]n=2005, 1981, and 1951 protein sequences for the short, medium and long protein length bins, respectively.

[a]Log-odds ratio (LOR) for differences between Actual and Shuffled data for the fraction of predicted disordered residues for a pair of segments. Subscript format "AB" specifies the segments being compared. $LOR_{AB}>0$ indicates a higher fraction for segment A than B. $LOR_{AB}<0$ indicates a lower fraction at segment A and $LOR_{AB}=0$ indicates identical fractions. Asterisks indicate significance levels for Z-statistics of odds ratios. The Bonferroni sequential method was employed for multiple test corrections (i.e., nine tests were corrected for each dataset). Superscript *** indicates $p<0.001$, respectively.

[b] Wald test statistic for a comparison of LORs between Actual and Shuffled data. Superscripts ns, *, ** and *** indicate non-significance, $p<0.05$, $p<0.01$ and $p<0.001$, respectively.

**S25 Table.** Log-odds ratios (LORs) for between putative functional classes for the fraction of predicted disordered residues.

| Focal class | Comparison class | | Protein length bin[1] | | | | | | | | |
|---|---|---|---|---|---|---|---|---|---|---|---|
| | | | Short | | | Medium | | | Long | | |
| | | | $LOR_N^a$ | $LOR_I^a$ | $LOR_C^a$ | $LOR_N^a$ | $LOR_I^a$ | $LOR_C^a$ | $LOR_N^a$ | $LOR_I^a$ | $LOR_C^a$ |
| Protein-binding | RNA-binding | Actual | 0.568$^{***}$ | 0.620$^{***}$ | 0.578$^{***}$ | 0.670$^{***}$ | 0.836$^{***}$ | 0.563$^{***}$ | 0.510$^{***}$ | 0.744$^{***}$ | 0.649$^{***}$ |
| | | Shuffled | 0.509$^{***}$ | 0.634$^{***}$ | 0.525$^{***}$ | 0.527$^{***}$ | 0.677$^{***}$ | 0.590$^{***}$ | 0.489$^{***}$ | 0.551$^{***}$ | 0.519$^{***}$ |
| | | $Z^b$ | 3.277$^{**}$ | -0.890$^{ns}$ | 2.854$^{**}$ | 5.377$^{***}$ | 15.018$^{***}$ | -1.074$^{ns}$ | 0.539$^{ns}$ | 20.905$^{***}$ | 3.488$^{**}$ |
| | DNA-binding | Actual | 0.322$^{***}$ | 0.380$^{***}$ | 0.345$^{***}$ | 0.444$^{***}$ | 0.568$^{***}$ | 0.290$^{***}$ | 0.333$^{***}$ | 0.509$^{***}$ | 0.239$^{***}$ |
| | | Shuffled | 0.276$^{***}$ | 0.442$^{***}$ | 0.276$^{***}$ | 0.270$^{***}$ | 0.381$^{***}$ | 0.303$^{***}$ | 0.353$^{***}$ | 0.354$^{***}$ | 0.225$^{***}$ |
| | | $Z^b$ | 2.657$^{*}$ | -4.161$^{***}$ | 3.872$^{***}$ | 6.982$^{***}$ | 19.223$^{***}$ | -0.539$^{ns}$ | -0.554$^{ns}$ | 17.790$^{***}$ | 0.422$^{ns}$ |
| | Linker | Actual | 0.484$^{***}$ | 0.235$^{***}$ | 0.588$^{***}$ | 0.678$^{***}$ | 0.384$^{***}$ | 0.609$^{***}$ | 0.467$^{***}$ | 0.353$^{***}$ | 0.368$^{***}$ |
| | | Shuffled | 0.467$^{***}$ | 0.330$^{***}$ | 0.500$^{***}$ | 0.369$^{***}$ | 0.269$^{***}$ | 0.516$^{***}$ | 0.513$^{***}$ | 0.235$^{***}$ | 0.396$^{***}$ |
| | | $Z^b$ | 0.960$^{ns}$ | -6.598$^{***}$ | 4.781$^{***}$ | 12.006$^{***}$ | 12.208$^{***}$ | 3.779$^{***}$ | -1.202$^{ns}$ | 14.018$^{***}$ | -0.778$^{ns}$ |
| RNA-binding | DNA-binding | Actual | -0.246$^{***}$ | -0.240$^{***}$ | -0.233$^{***}$ | -0.226$^{***}$ | -0.267$^{***}$ | -0.273$^{***}$ | -0.177$^{***}$ | -0.235$^{***}$ | -0.410$^{***}$ |
| | | Shuffled | -0.233$^{***}$ | -0.192$^{***}$ | -0.249 | -0.257$^{***}$ | -0.296$^{***}$ | -0.287$^{***}$ | -0.136$^{***}$ | -0.196$^{***}$ | -0.294$^{***}$ |
| | | $Z^b$ | -0.706$^{ns}$ | -2.874$^{**}$ | 0.812$^{ns}$ | 1.136$^{*}$ | 2.520$^{ns}$ | 0.543$^{*}$ | -1.003$^{ns}$ | -3.861$^{***}$ | -2.980$^{**}$ |
| | Linker | Actual | -0.084$^{**}$ | -0.385$^{***}$ | 0.011$^{ns}$ | 0.008$^{***}$ | -0.452$^{***}$ | 0.045$^{**}$ | -0.042$^{ns}$ | -0.391$^{***}$ | -0.281$^{**}$ |
| | | Shuffled | -0.043$^{***}$ | -0.303$^{***}$ | -0.025$^{ns}$ | -0.159$^{ns}$ | -0.408$^{***}$ | -0.074$^{*}$ | 0.024$^{ns}$ | -0.316$^{***}$ | -0.122$^{***}$ |
| | | $Z^b$ | -2.185$^{ns}$ | -4.995$^{***}$ | 1.777$^{ns}$ | 5.833$^{***}$ | -3.952$^{***}$ | 4.397$^{***}$ | -1.578$^{ns}$ | -7.781$^{***}$ | -3.940$^{***}$ |
| DNA-binding | Linker | Actual | 0.162$^{***}$ | -0.145$^{***}$ | 0.244$^{***}$ | 0.234$^{***}$ | -0.184$^{***}$ | 0.318$^{***}$ | 0.135$^{***}$ | -0.156$^{***}$ | 0.129$^{***}$ |
| | | Shuffled | 0.191$^{***}$ | -0.111$^{***}$ | 0.224$^{***}$ | 0.098$^{***}$ | -0.112$^{***}$ | 0.213$^{***}$ | 0.160$^{***}$ | -0.119$^{***}$ | 0.172$^{***}$ |
| | | $Z^b$ | -1.556$^{ns}$ | -2.156$^{*}$ | 1.034$^{ns}$ | 4.975$^{***}$ | -7.029$^{***}$ | 4.093$^{***}$ | -0.629$^{ns}$ | -4.031$^{***}$ | -1.133$^{ns}$ |

[1]n=2005, 1981, and 1951 protein sequences for the short, medium and long protein length bins, respectively.

[a]Extent of difference between putative functional classes for the fraction of predicted disordered residues. Subscripts specify segments and LOR>0 indicates a higher fraction for the focal class, while LOR<0 and LOR=0 indicate lower and equal fractions, respectively. Asterisks indicate significance levels for Z-statistics of odds ratios. The Bonferroni sequential method was employed for multiple test corrections (i.e., nine tests were corrected for each dataset). Superscripts ns, *, **, and *** indicate non-significance, $p<0.05$, $p<0.01$, $p<0.001$, respectively.

[b]Wald test statistic for a comparison of LORs between Actual and Shuffled data. Multiple test corrections and significance levels are implemented similarly as described in footnote a.

**S26 Table.** Statistics for AlphaFold database protein structures for non-*Drosophila* organisms.

| Organism | # PDB files | # protein sequences[1] | # residues | | | | |
|---|---|---|---|---|---|---|---|
| | | | total | min | mean | median | max |
| Human | 23,391 | 20,227 | 10,520,318 | 16 | 520.113 | 409 | 2,699 |
| Mouse | 21,615 | 21,592 | 10,878,190 | 16 | 503.807 | 382 | 2,698 |
| Zebrafish | 24,664 | 24,664 | 12,660,481 | 16 | 513.318 | 398 | 2,694 |
| *C. elegans* | 19,694 | 19,690 | 7,811,001 | 16 | 396.699 | 330 | 2,697 |
| *A. thaliana* | 27,434 | 27,370 | 11,015,547 | 17 | 402.468 | 347 | 2,696 |
| *S. cerevisiae* | 6,039 | 5,973 | 2,879,811 | 16 | 482.138 | 397 | 2,672 |
| *E. coli* | 4,363 | 4,321 | 1,340,642 | 16 | 310.262 | 272 | 2,358 |

[1]Cases where a given protein sequence was split into multiple PDB files were excluded. If the same protein sequence had multiple PDB files, one PDB file was chosen at random.
